# Loop statistics and fountain geometry reveal effective two-sided cohesin extrusion and crowding-induced arm desynchronization

**DOI:** 10.64898/2026.08.20.745948

**Authors:** Anastasia Chervinskaya, Mikhail S. Gelfand, Ralf Metzler, Kirill E. Polovnikov

## Abstract

Cohesin-driven loop extrusion shapes chromosome organization, yet what determines loop lengths and coordination between the extruding arms *in vivo* remains unclear. Broad extrusion “fountains” in Hi-C maps point to arm desynchronization, whose physical origin is unknown. Here we develop a kinetic theory of extrusion through transient chromatin roadblocks and neighbouring cohesins. Roadblock abundance, lifetime and partial permeability, together with cohesin crowding, combine into an effective obstacle density that renormalizes processivity and sets the mean loop length. Full loop-length distributions reveal extrusion symmetry: one-sided extrusion remains exponential, whereas effectively two-sided extrusion can generate a finite-length peak. ChIA-PET and MNase HiChIP data match the two-sided predictions, disfavouring purely one-sided extrusion. Cohesin crowding further desynchronizes the arms while leaving a residual correlation approaching *ρ* = 1/4. Fountain anisotropy across three vertebrates yields correlations near this value, consistent with local cohesin crowding as a sufficient mechanism for fountain formation. Fits further reveal 30–100-kb cohesin-loading regions and, in *Danio rerio*, a loading width comparable to the extent of enhancer enrichment, motivating a model in which an enhancer-rich loading platform acts as a collective barrier sustaining outward extrusion toward surrounding promoters. Our framework shows how cohesin and roadblock kinetics jointly determine loop statistics, shape chromosome-contact patterns, and may facilitate enhancer–promoter search.

## Introduction

Eukaryotic genomes are compacted into micrometre-sized nuclei while remaining accessible for transcription, enhancer–promoter communication, replication, and DNA repair. Chromosome folding is therefore an integral component of genome regulation [1, 2]. A major contribution to this organization comes from loop extrusion, in which cohesin—a chromosome-organizing motor of the structural-maintenance-of-chromosomes (SMC) family—loads onto chromatin and enlarges a DNA loop until it dissociates or encounters an obstacle. CTCF is the best-characterized sequence-specific barrier to cohesin, with blocking strongly dependent on CTCF orientation [3–8], but extrusion is also affected by other chromatin-bound proteins and by collisions with neighbouring cohesins [9, 10]. The kinetics of these encounters should therefore shape both loop sizes and the chromosome-contact patterns generated by extrusion.

Cohesin-mediated extrusion was proposed to explain topologically associating domains (TADs) and focal CTCF-dependent loops [3, 4] and was subsequently observed directly in single-molecule experiments [11–13]. Yet two fundamental properties of cohesin motion *in vivo* remain unresolved. First, does cohesin extrude from one side or from both sides of its loading site? Recent single-molecule studies indicate that extrusion can be asymmetric and switch direction, potentially in connection with NIPBL turnover or encounters with CTCF [14, 15], while regulator-resolved models predict intermittent extrusion controlled by cohesin-associated factors [16]. We therefore use *effectively two-sided* to describe extrusion in which both genomic directions contribute appreciably to loop growth over a cohesin residence time, irrespective of whether the two arms move simultaneously or alternate in time. Second, if both directions contribute, how strongly are the accumulated extensions of the two arms coordinated? Extrusion symmetry and arm coordination are related but distinct: the former determines which directions contribute to loop growth, whereas the latter describes how the two extensions fluctuate relative to one another.

Loop extrusion can be viewed as directed cohesin motion through a heterogeneous chromatin environment in which transiently bound obstacles and neighbouring extruders generate kinetic disorder. Most theoretical studies have focused on the three-dimensional consequences of extrusion, including chromosome-contact patterns, TAD formation, and the mechanics of loop-organized polymers [16–28]. Less is known about the statistics generated directly by the underlying one-dimensional extrusion process. In particular, how obstacle abundance, residence time, orientation-dependent permeability, and cohesin crowding jointly determine loop length, extrusion symmetry, and coordination between the two arms remains poorly understood.

Extrusion “fountains” (also termed jets) provide a complementary structural signature of these dynamics. These anisotropic contact-map features have been observed across several vertebrates and are associated with facilitated cohesin loading and enhancer-rich chromatin [29–32]. Unlike CTCF-anchored domains, fountains do not require fixed barriers and are thought to arise from outward extrusion initiated within regions of elevated cohesin loading [29–31]. Their transverse broadening implies incomplete coordination between the two extrusion directions, but the microscopic origin of this desynchronization and its relationship to cohesin crowding remain unclear.

Here we develop a kinetic theory of loop extrusion through transient, partially permeable chromatin roadblocks and neighbouring cohesins (Fig. 1). We first derive a universal expression for the mean loop length, in which diverse sources of kinetic disorder renormalize the bare cohesin processivity through a single effective obstacle density. The mean alone, however, cannot uniquely distinguish extrusion symmetry. We show that the full loop-length distribution provides a stronger diagnostic: strictly one-sided extrusion remains exponential, whereas effectively two-sided extrusion can develop a finite-length maximum because loop growth continues after one arm becomes blocked. Peaked loop-length distributions in RAD21 ChIA-PET, in CTCF-directed ChIA-PET and MNase HiChIP are quantitatively consistent with the corresponding two-sided predictions using independently constrained kinetic parameters.

**Figure 1:**
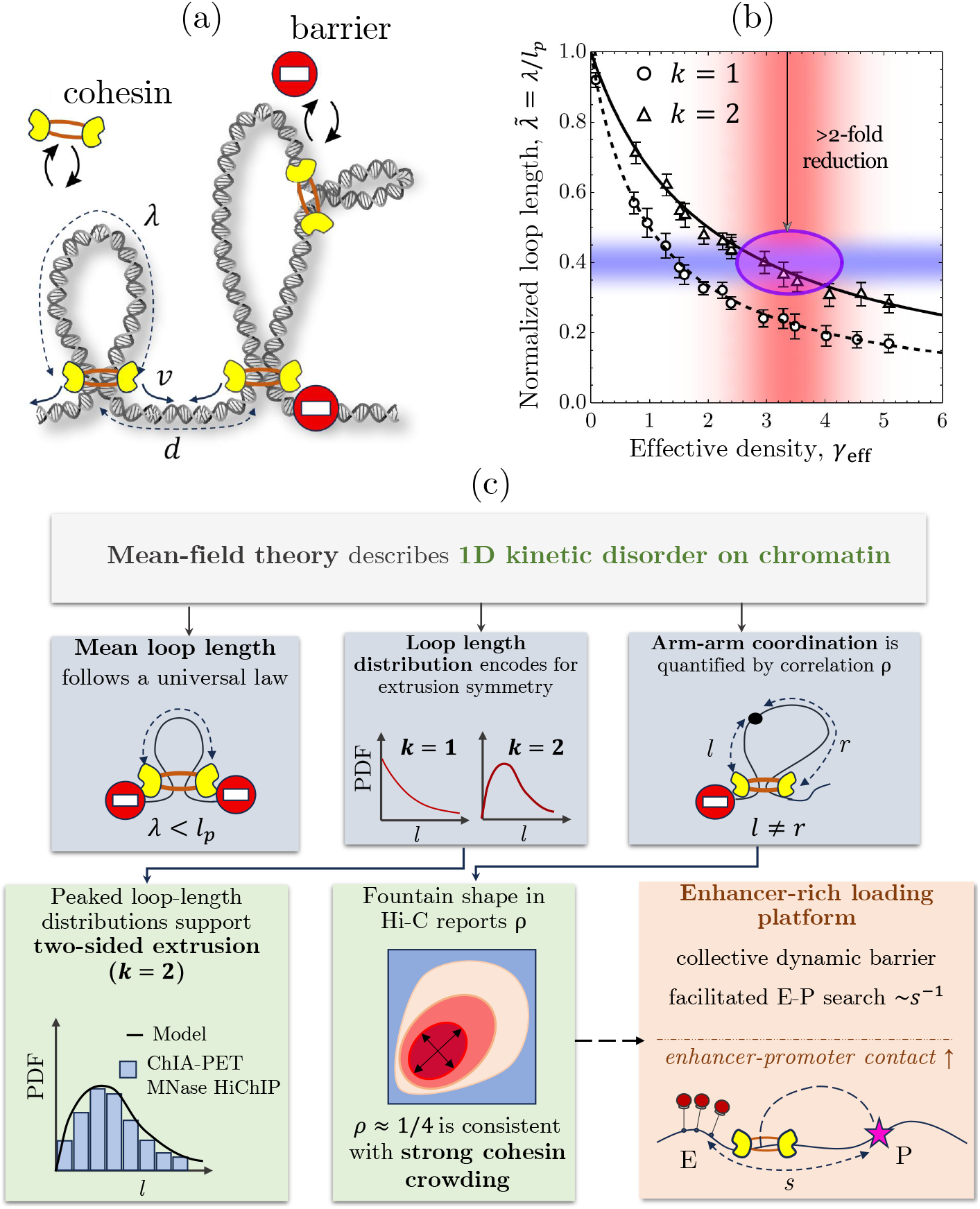
Chromatin disorder limits loop growth and links extrusion dynamics to chromosome organization. (a) Schematic of two-sided loop extrusion through transient chromatin roadblocks and neighbouring cohesins. A cohesin with residence time *τ* and bare processivity *l*_*p*_ = *vτ* extrudes at total speed *v*; encounters with obstacles reduce the mean loop length to *λ < l*_*p*_. Cohesin crowding is quantified by *γ* = *l*_*p*_*/d*, where *d* is the mean cohesin spacing. (b) Processivity-normalized mean loop length, 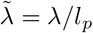, as a function of effective obstacle density *γ*_eff_. Solid and dashed curves show two-sided and one-sided extrusion, respectively; symbols denote stochastic simulations. The blue band marks the experimentally inferred 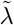 range in HeLa cells [6, 11, 12], and the red band the *γ*_eff_ range inferred from measured barrier kinetics (Table S2). Their overlap lies on the two-sided branch and implies a more than twofold disorder-induced reduction of loop length relative to *l*_*p*_. (c) Summary of the framework and experimental readouts. Mean loop length is controlled by *γ*_eff_, whereas the full loop-length distribution reports effective extrusion symmetry. Peaked ChIA-PET and MNase HiChIP loop-length distributions support effectively two-sided extrusion. Fountain anisotropy reports arm–arm coordination, with correlations near *ρ* = 1/4 consistent with strong local cohesin crowding. Enhancer-rich loading regions provide a potential genomic mechanism for generating this crowded state and promoting outward enhancer–promoter contacts.

The theory further identifies the correlation between the two arm extensions as a distinct dynamical observable. Cohesin crowding progressively desynchronizes the arms but leaves a characteristic residual correlation, *ρ* = 1/4, whereas sufficiently persistent chromatin roadblocks can reduce it further. We show that this otherwise hidden variable is encoded in fountain anisotropy: full kinetic-model fits to fountains from three vertebrates yield arm correlations close to the crowding limit and require strong local cohesin accumulation. The inferred cohesin-loading regions span tens of kilobases and, in *Danio rerio*, closely match the spatial extent of enhancer enrichment. These observations motivate a model in which enhancer-rich regions act as distributed cohesin-loading platforms that both generate local crowding and provide a collective barrier to inward extrusion (Fig. 1c). Such platforms could retain one cohesin arm while the other extrudes outward, changing the optimized promoter-capture envelope from ~ *s*^−2^ for an isolated enhancer to ~ *s*^−1^ for platform-mediated retention, where *s* is the genomic distance to promoter. Together, loop statistics and fountain geometry provide complementary readouts of effective extrusion symmetry and arm coordination and connect these microscopic dynamics to regulatory chromosome organization.

## Results

### A kinetic framework for cohesin extrusion on a disordered chromatin track

We consider a single loop-extruding cohesin motor moving on chromatin in a steady-state background of DNA-bound obstacles and other cohesins. Upon loading, cohesin remains bound for a mean residence time *τ* and enlarges a loop of length *x* at total speed *v* until it dissociates or is blocked by an obstacle [33]. We distinguish strictly one-sided extrusion (*k* = 1), in which only one cohesin arm is active, from symmetric two-sided extrusion (*k* = 2), in which both arms move simultaneously; in both cases *v* denotes the total loop-growth speed. In the absence of obstacles, the intrinsic processivity

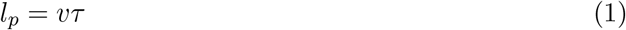

sets both the characteristic loop scale and the exponential loop-length distribution.

*In vivo*, extrusion proceeds through a heterogeneous landscape of chromatin-bound roadblocks. For generality, we consider *n* classes of transiently bound obstacles, *i* = 1, …, *n*, each characterized by a mean residence time *τ*_*i*_ and a mean spacing *d*_*i*_ between successful blocking events (Fig. 1a). Thus, *d*_*i*_ already incorporates partial permeability: a roadblock population with blocking probability *p*_*i*_ *<* 1 and raw spacing 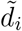 is represented as an effectively sparser population of fully blocking barriers, with 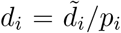. Such permeability can arise, for example, from CTCF binding polarity and the resulting orientation-dependent blocking [7, 8]. Neighboring cohesins constitute a separate obstacle class with mean spacing *d*, which likewise absorbs, at the mean-field level, the probability of mutual bypassing.

This kinetically disordered environment is conveniently described by a small set of dimensionless control parameters,

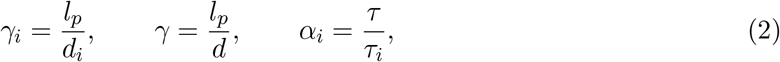

where *γ*_*i*_ and *γ* quantify the crowding by static roadblocks and neighboring cohesins, respectively, while *α*_*i*_ measures cohesin residence relative to roadblock persistence. Long-lived obstacles correspond to *α*_*i*_ ≪ 1, whereas short-lived obstacles satisfy *α*_*i*_ ≳ 1. For cohesin obstacles, *α*_*c*_ = 1 by definition. Thus, the extrusion kinetics is controlled by three biologically meaningful ingredients: the intrinsic motor processivity, the density of successful blocking events, and the lifetime of the blocking species.

To compute stationary loop statistics, we formulate a state-resolved mean-field framework in which a cohesin is classified by whether its arms are free or blocked by a given obstacle class. The resulting balance equations combine cohesin loading with advection in loop length, stochastic blocking, obstacle release, and cohesin dissociation. This framework allows us to move from the microscopic kinetics of cohesin-roadblock encounters to three experimentally accessible observables: the mean loop length *λ*, the loop-length distribution *N* (*x*), and the correlation between two arm extensions *ρ*. As we show below, these observables play distinct roles. The mean loop length is controlled by a single renormalized obstacle density, whereas the full loop statistics and arm-arm correlation reveal the symmetry and internal coordination state of the extruding motor.

### Roadblock kinetics set the loop length through an effective obstacle density

In the dilute limit, when *γ* and *γ*_*i*_ are both much smaller than unity, encounters with roadblocks are rare and the stationary mean loop length *λ* approaches the intrinsic processivity *l*_*p*_. At finite obstacle density, repeated blocking and release events suppress loop growth. For an arbitrary number of roadblock classes the mean-field theory yields a simple universal result:

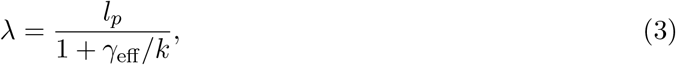

where the effective density *γ*_eff_ is additive in all static roadblocks and other cohesins:

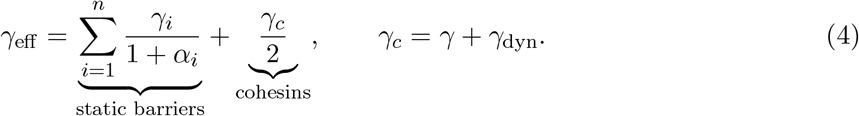

Here *γ*_dyn_ is a self-consistent correction arising from the relative motion of neighboring extruders (see below). Eq. (3) suggests that the mean loop length is controlled not by the bare processivity alone, but by the renormalized obstacle density *γ*_eff_ that incorporates both static roadblocks and neighboring cohesins. Accordingly, in the obstacle-free limit, *γ*_eff_ → 0 and *λ* → *l*_*p*_. For a fixed total extrusion speed *v*, two-sided extrusion (*k* = 2) reduces the speed of each arm to *v/*2, halves the encounter rate, and therefore yields a larger *λ* than one-sided extrusion (*k* = 1), see Fig. 1b.

The first term in Eq. (4) contains the contributions of *n* static roadblock classes. Long-lived roadblocks (*α*_*i*_ ≪ 1) contribute nearly their full density *γ*_*i*_, whereas short-lived roadblocks are suppressed by the factor 1/(1 + *α*_*i*_). Thus, barrier abundance alone does not determine the effect on loop growth; it is the density weighted by barrier persistence that matters. For a single barrier class *b*, we therefore define the effective barrier density 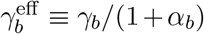, which we use below when comparing the theory with experiment (Table S2).

The cohesin contribution *γ*_*c*_ in Eq.(4) contains the static term *γ* and the dynamic correction *γ*_dyn_, both reduced by the cohesin persistence factor (1 + *α*_*c*_)^−1^ = 1/2. Unlike static roadblocks, this correction originates from the relative motion of neighboring extruders and is therefore most important in the dilute limit, where *γ*_dyn_ ≈ (*k/*2)*γ* for *γ, γ*_*i*_ → 0 (see Supplementary Section 2). Under experimentally relevant conditions, *γ* ≈ 2.7 (Table S2), it remains small, *γ*_dyn_ ≈ 0.4 ≪ *γ*. Most cohesins therefore behave effectively as blocked obstacles with *α*_*c*_ = 1, so the extruder mobility contributes only weakly to *γ*_eff_.

Equation (3) also admits a direct kinetic interpretation for the processivity-normalized mean loop length 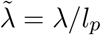 that is convenient for experimental tests (Fig. 1b):

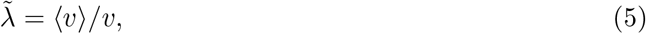

where ⟨*v*⟩ is the average loop-growth speed. In HCT116 cells [6], recovery of a 900±50 kb Hi-C loop occurs within ≈ 40 min after cohesin restoration, therefore ⟨*v*⟩ ≈ 0.38 kb/s; together with single-molecule measurements of the intrinsic cohesin speed *v* ≈ 1 kb/s [11, 12] this yields 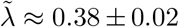 for the normalized loop length. Independently, inserting measured kinetic parameters for CTCF and MCM complexes (Table S2) into Eq.(4) yields *γ*_eff_ ≈ 3.4 ± 0.7 and *λ/l*_*p*_ ≈ 0.37 ± 0.05 for two-sided extrusion, but only about 0.23 ± 0.04 for one-sided extrusion. The observed mean loop length is therefore quantitatively consistent with effectively two-sided, but not strictly one-sided extrusion.

Stochastic simulations of active extrusion with explicit loading, dissociation, steric interactions, and transient barriers confirm the mean-field result (Eq. (3)) across the relevant parameter range (Fig. 1b, Fig. S8). In particular, the simulated mean loop length follows the same universal dependence on *γ*_eff_ for both one-sided and two-sided extrusion, showing that the reduction of loop length by chromatin disorder is well captured by a single coarse-grained control parameter. These results thus provide a unified description of how roadblock density, lifetime, permeability, extruder crowding, and cohesin turnover shape the mean loop scale *in vivo*.

### WAPL depletion provides a perturbational test of the universal loop-length law

WAPL (Wings apart-like protein homolog) depletion simultaneously prolongs cohesin residence on chromatin and increases the abundance of chromatin-bound cohesin [34, 35]. These effects act oppositely on the mean loop length: the longer residence time increases the bare processivity, *l*_*p*_ = *vτ*, whereas the reduced cohesin spacing increases *γ*_eff_ through more frequent cohesin–cohesin encounters. We therefore compared matched wild-type and ΔWAPL measurements with Eq. (3) (Fig. S1).

Across the wild-type datasets, the two-sided theory is consistent with an effective cohesin spacing *d*_WT_ ≈ 174 kb [24]; the ΔWAPL measurements are reproduced when this spacing decreases to *d*_ΔWAPL_ ≈ 104 kb. Although WAPL depletion increases *l*_*p*_ by approximately 14-fold, the accompanying increase in chromatin-bound cohesin density strongly increases *γ*_eff_. These effects largely cancel for one-sided extrusion, which underpredicts the observed loop-length increase, whereas the two-sided theory quantitatively reproduces the perturbation (Fig. S1). WAPL depletion therefore provides an independent perturbational test favouring effectively two-sided extrusion.

### Loop-length distributions distinguish one-sided and two-sided extrusion

The mean loop length already provides quantitative information about extrusion symmetry: when combined with independently constrained obstacle densities and cohesin kinetics, both the steady-state comparison (Fig. 1b) and the WAPL-depletion perturbation (Fig. S1) favor *k* = 2. However, because the mean depends on the combination *γ*_eff_ */k* (Eq. (3)), it cannot by itself uniquely determine the extrusion symmetry *k*. We therefore asked whether the full stationary loop-length distribution *N* (*x*) provides a more discriminating signature. In the weak-disorder limit, *γ*_eff_ ≪ 1, obstacles rarely perturb extrusion, and for either *k* the distribution approaches the obstacle-free exponential set by the cohesin residence time, with mean *l*_*p*_ = *vτ*. Once roadblocks become sufficiently abundant or persistent to constrain loop growth, however, the distributions generated by one-sided and two-sided extrusion become qualitatively distinct.

The stationary distribution follows from the state-resolved kinetic equations for freely extruding, partially blocked, and fully blocked cohesins (Supplementary Section 3). For one-sided extrusion, only the freely extruding state can increase the loop length; blocking of the active arm arrests further growth. For two-sided extrusion, by contrast, one arm can remain mobile after the other is blocked, introducing additional growing states and hence additional characteristic length scales. The exact stationary solution is therefore a single exponential for one-sided extrusion but a finite sum of exponential modes for two-sided extrusion.

This distinction becomes transparent by representing the kinetics as a directed length-state graph (Fig. 2a,b). Each node denotes a cohesin state defined by whether its arm or arms are free or blocked, while directed edges represent allowed transitions between these states upon an increase of loop length. Unlike a conventional kinetic graph organized in time, the resulting length-state graph therefore describes propagation directly in loop-length space. For a single roadblock class *b* together with neighbouring cohesins *c*, one-sided extrusion contains the states {*A*_0_, *A*_*b*_, *A*_*c*_}, whereas two-sided extrusion requires the enlarged state space {*A*_0_, *A*_*b*_, *A*_*c*_, *A*_*bb*_, *A*_*bc*_, *A*_*cc*_}. Crucially, nodes differ in whether loop length can still increase: in one-sided extrusion only *A*_0_ is propagating, whereas in two-sided extrusion the states *A*_*b*_ and *A*_*c*_, with one blocked and one mobile arm, remain propagating. The number of such propagating states determines the number of exponential decay modes in the stationary loop-length distribution; fully blocked states inherit these modes without introducing additional length scales.

**Figure 2:**
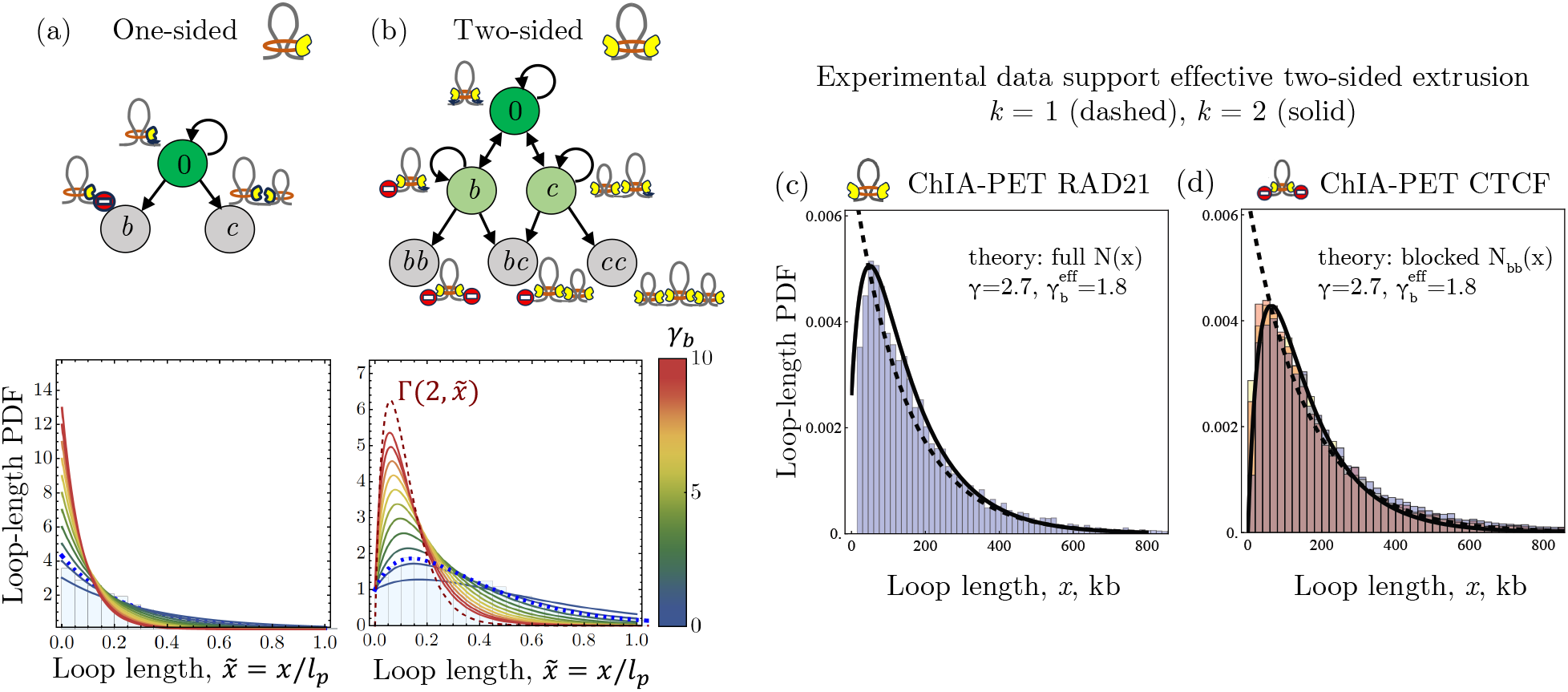
Loop-length distributions distinguish extrusion symmetry and support effective two-sided extrusion. (a,b) Length-state graphs (top) and theoretical loop-length distributions *N* (*x*) (bottom) for one-sided (*k* = 1) and two-sided (*k* = 2) extrusion. States 0, *b*, and *c* denote freely extruding arms and blocking by a chromatin barrier or neighbouring cohesin, respectively. For one-sided extrusion, only the free state propagates in loop-length space, yielding a single exponential distribution. For two-sided extrusion, states with one blocked and one mobile arm remain propagating, generating additional decay modes and a finite-length maximum. Curves are shown for increasing barrier density *γ*_*b*_ at fixed *γ* = 2.7 and *α* = 0; the red dashed curve denotes the limiting shape-2 Gamma distribution. Blue histograms show stochastic simulations at *γ*_*b*_ = 1.8, with dotted curves indicating the analytical predictions. (c) RAD21 ChIA-PET loop-length distribution in GM12878 cells [36], compared with the full theoretical distribution *N* (*x*). (d) CTCF ChIA-PET loop-length distributions in GM12878 (blue), K562 (yellow), and HeLa (pink) cells [37], compared with the doubly blocked distribution *N*_*bb*_(*x*). Dashed and solid black curves show the one-sided and two-sided predictions, respectively, evaluated using independently estimated parameters *γ* = 2.7, 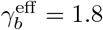, and *l*_*p*_ = 431 kb (Tables S1,S2), without fitting to the displayed distributions. The observed finite-length maxima are captured by the effective two-sided theory.

For one-sided extrusion, the graph therefore contains a single propagating state (Fig. 2a). Once the process leaves this state, further loop growth is impossible, and the stationary distribution is necessarily

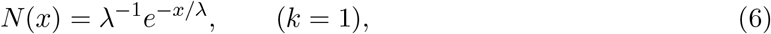

independent of the number of roadblock classes, their densities, or lifetimes.

For two-sided extrusion, blocking one arm does not terminate growth as long as the second arm remains mobile (Fig. 2b). With *n* roadblock classes together with neighbouring cohesins, there are *n* + 2 propagating states, and the stationary distribution is generically

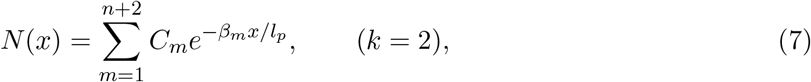

where *β*_*m*_ *>* 0 define the characteristic decay scales and *C*_*m*_ their weights. Unlike the single exponential of Eq. (6), this superposition can develop a maximum at nonzero loop length: loops first extend before accumulating in partially and fully blocked states. In the weak-disorder limit these states are rarely populated, and the distribution continuously returns to the obstacle-free exponential.

The same decay modes appear in the doubly barrier-blocked population (node “bb” in Fig. 2b), whose normalized distribution we denote by *N*_*bb*_(*x*). In the minimal model with one roadblock class and neighbouring cohesins, both *N* (*x*) and *N*_*bb*_(*x*) therefore contain three exponential modes, with different statistical weights (Supplementary Section 3.1.2). Notably, *N*_*bb*_(0) = 0 for two-sided extrusion because both arms must first extend before the loop can become blocked at both ends. Consequently, *N*_*bb*_(*x*) develops a pronounced finite-length maximum over a broad range of barrier parameters (Fig. S12a).

Importantly, these loop statistics diagnose effective rather than instantaneous two-arm motion. Extrusion may proceed simultaneously from both sides or through alternating activity in the two genomic directions. If switching is rare compared with cohesin dissociation, most trajectories remain effectively one-sided; sufficiently frequent switching allows both directions to contribute appreciably within one residence time. Stochastic alternation and blockage-triggered switching generate length-state graphs with multiple propagating states analogous to simultaneous two-arm extrusion (Supplementary Section 3.2.2; Fig. S5). Thus, the multiexponential signature applies more generally to effectively two-sided extrusion.

For *k* = 2, formation of a finite-length maximum requires sufficiently strong kinetic disorder. With long-lived roadblocks (*α* → 0) and negligible cohesin crowding, the threshold is 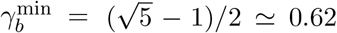, increasing to 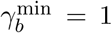 for rapidly turning-over roadblocks (Supplementary Section 3.1; Fig. S2). The independently constrained *in vivo* parameters lie within the regime where a pronounced peak is expected (Table S2). Stochastic simulations reproduce this distinction: one-sided distributions remain exponential, whereas two-sided distributions become multiexponential and develop finite-length maxima as disorder increases (Fig. 2a,b; Figs. S6,S7).

These results provide two complementary experimental tests of effective extrusion symmetry. Cohesin-associated interactions probe the full distribution *N* (*x*), whereas loops anchored by extrusion barriers at both ends probe *N*_*bb*_(*x*). Both can exhibit a finite-length maximum for effectively two-sided extrusion, whereas strictly one-sided extrusion remains monotonic. We therefore next compare these predictions with cohesin- and CTCF-associated chromosome-contact datasets.

### Experimental loop-length statistics support effective two-sided extrusion *in vivo*

We tested these predictions using five published protein-directed chromosome-contact datasets: RAD21 ChIA-PET in GM12878, CTCF ChIA-PET in GM12878, K562 and HeLa-S3, and CTCF MNase HiChIP in K562 [36–38]. RAD21-associated interactions were compared with the full distribution *N* (*x*), whereas CTCF-directed interactions, which preferentially report barrier-anchored loops, were compared with the doubly blocked distribution *N*_*bb*_(*x*).

The curves in Fig. 2c,d are predictions evaluated with a common reference parameter set constrained from published molecular measurements (*γ* = 2.7, 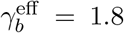, *l*_*p*_ = 431 kb, *α* = 1.2; Tables S1,S2), rather than fits to the displayed distributions. All five datasets exhibit finite-length maxima captured by the two-sided prediction, whereas the one-sided prediction remains monotonic. Across assays and cell types, theory–experiment NMAEs are approximately 9–13% (Table S9), comparable to or smaller than the 16.9% difference between independent GM12878 RAD21 replicates (Fig. S12b).

Loop-length profiles alone do not uniquely identify the microscopic kinetic parameters: NMAE landscapes contain broad low-error valleys (Fig. S12d–f). We therefore use the distributions as a forward consistency test rather than for parameter fitting. The independently constrained reference region lies within the low-error regime for all three cell lines, and varying each parameter over its experimental uncertainty preserves the characteristic finite-length maximum (Fig. S13). Thus, the qualitative inference of effective two-sided extrusion is considerably more robust than inference of individual kinetic parameters.

### Roadblocks and cohesins control coordination between the arms

The preceding results show that effectively two-sided extrusion produces loop-length statistics richer than those of a one-sided motor. The total loop length, however, does not specify how it is divided between the arms: the same *L* = *ℓ* + *r* may arise from nearly symmetric motion, *ℓ* ≃ *r*, or strongly asymmetric extrusion, *ℓ* ≫ *r* or *r* ≫ *ℓ*. We therefore ask whether the two arms advance independently or retain a memory of their common extrusion history. This distinction is biologically relevant because the joint statistics of *ℓ* and *r*, rather than their sum alone, determine two-dimensional patterns around cohesin-loading sites in chromosome-contact maps.

A useful benchmark is provided by statistically independent arms. Since each arm has an exponential extension distribution with mean *λ/*2, their sum follows a Gamma distribution with shape parameter 2, *N* (*x*) = 4*xλ*^−2^ exp(−2*x/λ*). This therefore represents the independent-arm limit. In general, however, *N* (*x*) is not Gamma (Fig. 2b), approaching this form only for sufficiently dense and persistent roadblocks (*γ*_*b*_ → ∞, *α* = 0). Thus, the obstacle landscape controls not only loop length but also arm coordination.

To quantify this coordination, we introduce the joint distribution *A*(*ℓ, r*) and the Pearson correlation coefficient

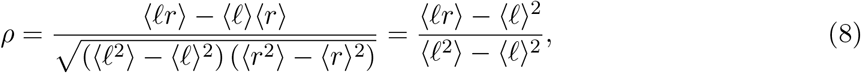

where the second equality follows from left–right symmetry, ⟨*ℓ*⟩ = ⟨*r*⟩ = *λ/*2 and Var(*ℓ*) = Var(*r*). Here *ρ* = 1 denotes perfectly correlated arm extensions, *ρ* = 0 statistically independent arms, and intermediate values partial coordination.

A central result emerges in the absence of static roadblocks, *γ*_*b*_ = 0, when the only obstacles are neighbouring cohesins. Even arbitrarily frequent cohesin–cohesin collisions do not make the arm extensions independent. Instead, in the strong-crowding limit, *γ*_*c*_ → ∞,

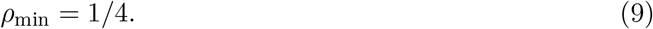

Thus, cohesin crowding strongly desynchronizes the arms but leaves a finite correlation floor.

The origin of this 1/4 floor is the shared cohesin residence time. At high density, the two arms experience statistically independent collision histories once the lifetime *T* of the tagged cohesin is fixed, but both share the same *T*. Long-lived cohesins therefore tend to generate longer extensions on both sides, whereas short-lived cohesins generate shorter ones. This common-lifetime fluctuation produces a residual positive covariance even when the local collision histories are independent. For collisions between identical cohesins, variance decomposition gives *ρ* = 1/4 as *γ*_*c*_ → ∞ (Supplementary Section 4.2).

Static roadblocks can produce stronger desynchronization because one arm may remain arrested for a substantial fraction of the cohesin lifetime. For persistent roadblocks, the high-density limit is

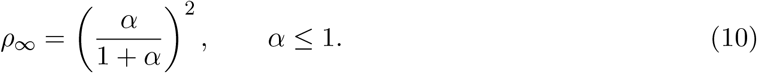

Roadblocks with a cohesin-like lifetime, *α* = 1, recover *ρ* = 1/4, whereas increasingly persistent barriers drive *ρ*_∞_ toward zero. Persistent roadblocks can therefore reduce arm correlation below the crowding floor.

When static roadblocks and neighbouring cohesins coexist, increasing cohesin density can either decrease or increase arm correlation, depending on the roadblock regime (Fig. 3a,b). For *α <* 1, the crossover occurs at

**Figure 3:**
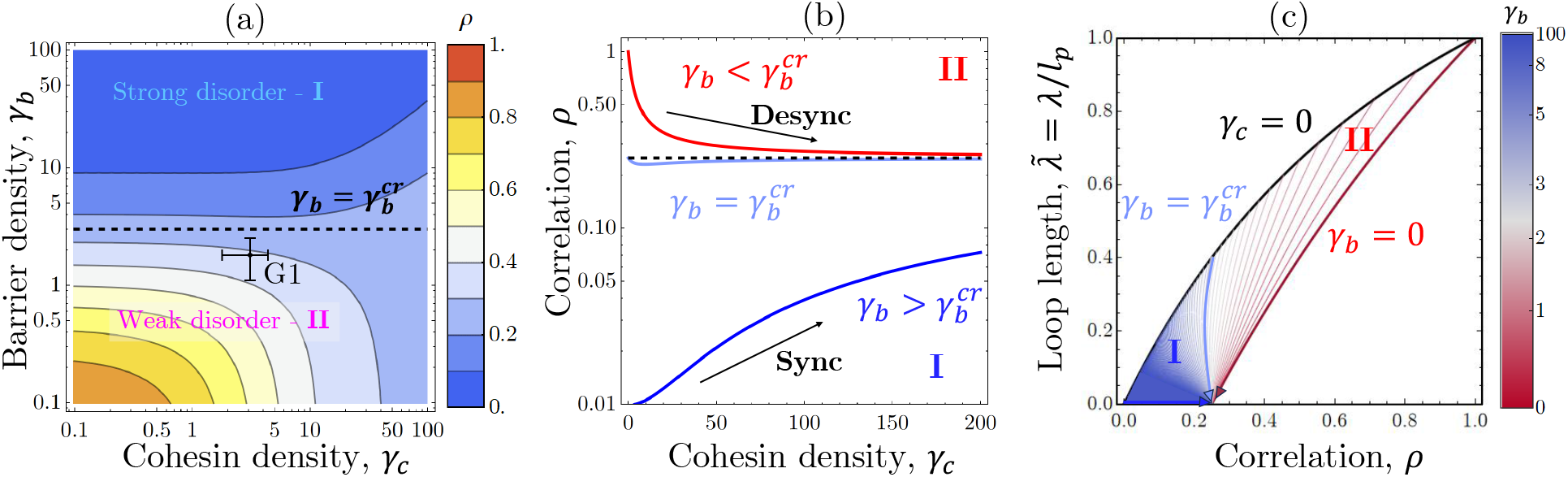
Roadblocks and cohesin crowding control coordination between cohesin arms. The two obstacle classes are neighboring cohesins, described by *γ*_*c*_ = *γ* + *γ*_dyn_, and static roadblocks, with density *γ*_*b*_ and persistence *α*; in this figure *α* = 0. (a) Pearson correlation coefficient *ρ* between the arm extensions in the (*γ*_*c*_, *γ*_*b*_) plane. The dashed line marks the critical roadblock density, 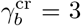, separating the strong-roadblock regime I from the weak-roadblock regime II. The experimentally inferred G1 state lies in regime II, close to the synchronization–desynchronization boundary. (b) Arm correlation *ρ* as a function of cohesin density *γ*_*c*_ for representative roadblock densities below (*γ*_*b*_ = 0, red), at (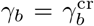, light blue), and above (*γ*_*b*_ = 100, blue) the critical value. Increasing cohesin density desynchronizes the arms in regime II, leaves *ρ* nearly unchanged at criticality, and increases synchronization in regime I. In the absence of static roadblocks, the correlation approaches the universal crowding-induced lower bound *ρ* = 1/4. (c) “Sail plot” showing the normalized mean loop length 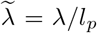 against the arm correlation *ρ*. The black boundary corresponds to roadblocks alone (*γ*_*c*_ = 0), whereas the red boundary corresponds to pure cohesin crowding (*γ*_*b*_ = 0). The interior curves show trajectories at fixed *γ*_*b*_ upon increasing *γ*_*c*_, with color denoting the roadblock density. In regime II, increasing cohesin density shortens the loop and decreases *ρ*; in regime I, it shortens the loop but increases *ρ*. The critical trajectory 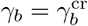 separates these responses.

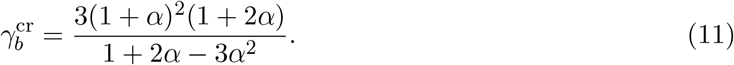

For 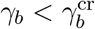 (regime II), cohesin–cohesin encounters dominate and increasing *γ*_*c*_ decreases both the mean loop length and *ρ*, driving the latter toward 1/4. For 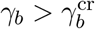 (regime I), persistent roadblocks have already reduced *ρ* below this value; adding more transient cohesin obstacles instead restores coordination and pulls *ρ* back toward 1/4.

The smallest critical density occurs for infinitely persistent barriers, *α* = 0,

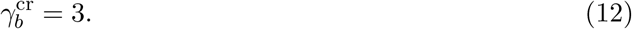

Roadblocks must therefore be spaced, on average, by no more than one-third of the processivity length before their effect on arm coordination exceeds that of cohesin crowding. As roadblock turnover increases, a larger density is required; for *α >* 1, the critical density disappears and the system remains in regime II. Arm coordination is therefore governed not simply by the total obstacle density, but by the relative abundance and persistence of cohesin motors and roadblocks.

The “sail plot” in Fig. 3c summarizes these regimes by placing the normalized mean loop length 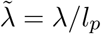 and arm correlation *ρ* on the same diagram. The unobstructed state lies at 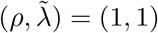. Along the roadblock-only boundary (*γ*_*c*_ = 0), persistent barriers can reduce both quantities toward zero, whereas along the pure-crowding boundary (*γ*_*b*_ = 0), increasing *γ*_*c*_ shortens loops but cannot reduce *ρ* below 1/4. At fixed *γ*_*b*_, increasing crowding therefore drives trajectories toward lower *ρ* in regime II but toward higher *ρ* in regime I, with 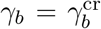 separating the two responses. All crowding-dominated trajectories converge toward the characteristic value *ρ* = 1/4.

Using experimentally constrained densities and residence times for CTCF, MCM, and cohesin (Table S2), the effective single-obstacle model places interphase chromatin in regime II, although relatively close to the transition (Fig. 3a). CTCF is comparatively dynamic (*α*_CTCF_ ≈ 1.2) and sparse 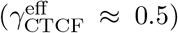, whereas MCM is highly persistent (*α*_MCM_ ≈ 0) but remains below the critical density 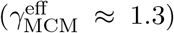. Approximating their combined effect by a persistent (*α* = 0) effective roadblock class gives *γ*_*b*_ ≈ 1.8, still below 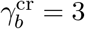 but closer to the transition.

Using the analytical expression for *ρ* (Supplementary Eq. (140)) with the independently constrained interphase parameters (Table S2), the theory predicts *ρ* ≈ 0.32 ± 0.06, indicating substantial but incomplete arm desynchronization. Changes in barrier abundance or residence time can therefore modulate arm coordination *in vivo*.

### Fountain anisotropy reveals the cohesin-crowding correlation floor

The arm–arm correlation *ρ* should leave a measurable signature in chromosome-contact maps. Extrusion fountains are cohesin-dependent features associated with facilitated cohesin loading [29–31]: preferential loading within a localized genomic region followed by effectively two-sided extrusion generates a feature extending approximately perpendicular to the Hi-C diagonal [29, 30]. Similar extension of the two arms produces contacts near *ℓ* ≃ *r*, whereas interruption of either arm creates an imbalance between *ℓ* and *r* and broadens the fountain transversely (Fig. 4a). Fountain anisotropy can therefore report the degree of arm coordination. We asked whether the broad fountains observed *in vivo* are quantitatively consistent with the desynchronization generated by local cohesin crowding.

**Figure 4:**
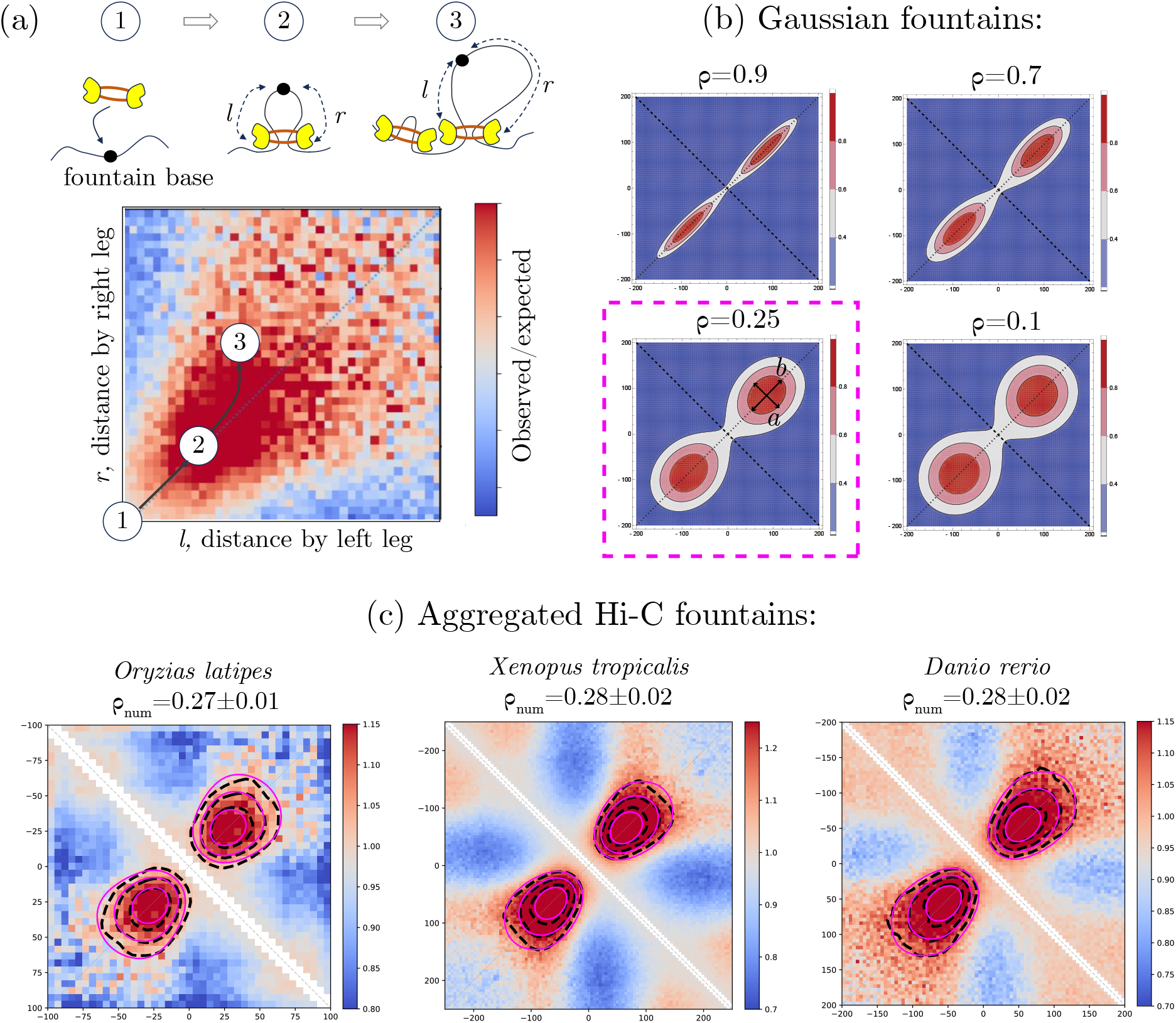
Fountain anisotropy reports cohesin-arm coordination. (a) Schematic fountain formation in (*ℓ, r*) space, where *ℓ* and *r* are the genomic distances extruded by the two cohesin arms from the fountain base. Coordinated arm extension generates contacts along *ℓ* ≃ *r* (state 2), whereas interruption of either arm produces an imbalance between *ℓ* and *r*, broadening the fountain transversely (state 3). (b) Gaussian fountain profiles for representative arm–arm correlations. Strong coordination produces a narrow, elongated fountain, whereas decreasing *ρ* broadens the profile. At the cohesin-crowding limit, *ρ* = 1/4, the fountain remains anisotropic. Gaussian widths parallel and perpendicular to the fountain axis are denoted by *b* and *a*, respectively. (c) Aggregated observed/expected Hi-C fountains from medaka fish (*Oryzias latipes*), western clawed frog (*Xenopus tropicalis*), and zebrafish (*Danio rerio*). Dashed black contours show the experimental signal and magenta contours the numerical kinetic-model fits, with normalized root-mean-square errors of 0.09, 0.11, and 0.09, respectively (Table S7). The fitted arm–arm correlations are consistent with the Gaussian estimates and lie close to the theoretical cohesin-crowding limit *ρ* = 1/4.

A simple model-light estimate of *ρ* follows from the central fountain ellipticity. For two statistically equivalent Gaussian variables,

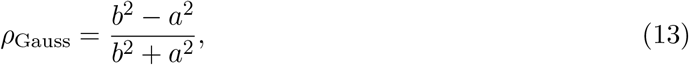

where *b* and *a* are the widths parallel and perpendicular to the fountain axis, respectively (Fig. 4b). Thus, increasingly transverse fountain broadening corresponds to decreasing arm correlation.

We analysed aggregated fountains from medaka fish (*Oryzias latipes*) [39], western clawed frog (*Xenopus tropicalis*) [40], and zebrafish (*Danio rerio*) [30] (Fig. 4c). Fountains were identified genome-wide using the *fontanka* algorithm [30], aligned at their inferred bases, and converted to observed/expected Hi-C signal to remove the dominant genomic-distance dependence. This normalization does not make the signal a literal joint distribution of arm extensions, because three-dimensional polymer fluctuations also contribute to contact formation [24, 25]. Nevertheless, the central fountain contours are approximately elliptical. Gaussian fits within the central 80-kb window reproduce these contours with NRMSEs below 11% and yield *ρ*_Gauss_ ≈ 0.25–0.30 across the three species (Table S7). The estimates remain stable to fitting-window size and bootstrap resampling (Table S6).

For quantitative inference, we fitted the complete fountain profiles directly with the numerical joint arm-extension distribution *A*(*ℓ, r*) from the kinetic model (Fig. 4c and Fig. S9). The model allows cohesin loading to be distributed over a region of width *σ*_*p*_ around the fountain base, within which the local cohesin density is elevated to *γ*_*c*_. We initially fixed the static-roadblock density at the independently estimated reference value *γ*_*b*_ = 1.8 and assumed long-lived roadblocks, *α* = 0 (Table S2); *l*_*p*_ and *σ*_*p*_ set the genomic extent and loading-zone width, whereas *γ*_*c*_ controls the arm correlation.

The numerical model reproduces all three fountain profiles with full-window NRMSEs of 0.09– 0.11 (Table S7). The correlations calculated from the fitted kinetic parameters fall within the narrow range *ρ*_num_ ≈ 0.27–0.28, in agreement with the independent Gaussian estimates. Varying *γ*_*b*_ over its experimentally constrained range changes the inferred *ρ*_num_ by no more than approximately 10% (Table S8), showing that the inferred partially desynchronized state is robust to uncertainty in the static-roadblock contribution. Thus, the model-light fountain ellipticity and the full kinetic-model fits provide mutually consistent estimates of arm coordination.

The fitted correlations lie close to the theoretical cohesin-crowding limit *ρ*_min_ = 1/4 (Eq. (9)). Furthermore, reproducing the observed anisotropy requires high local cohesin densities, *γ*_*c*_ ≈ 10– 14 (Table S7), approximately three-to fivefold above the genome-wide HeLa estimate (Table S2), distributed over loading regions of width 2*σ*_*p*_ ≈ 30–100 kb. The fits also require enhanced bare processivities, *l*_*p*_ ≈ 1.7–3.8 Mb, substantially above the genome-wide estimate (Table S1), suggesting that fountain regions combine enhanced cohesin loading with locally prolonged cohesin residence and/or increased extrusion speed. Together, these results show that local cohesin accumulation is quantitatively sufficient to produce the observed degree of arm desynchronization and fountain geometry.

The correlation near 1/4 is not, however, unique to crowding. In the minimal alternating-arm model of Supplementary Section 4.4, symmetric Poisson switching alone yields *ρ* = 1/4 at *wτ* = 7/6, corresponding to approximately 1.17 switches per cohesin residence time. The recurrent *ρ*_num_ ≈ 0.27–0.28 (Fig. 4c) therefore does not identify crowding as a unique microscopic mechanism, but demonstrates that cohesin–cohesin encounters are quantitatively sufficient to account for the observed fountains. These correlations remain compatible with sufficiently rapid arm switching, whereas slower switching can reduce *ρ* below the crowding floor and may describe more strongly desynchronized regimes in other systems.

### Enhancer-rich regions may form loading platforms that couple fountain formation to promoter capture

What genomic environment could generate the locally crowded extrusion regime inferred from fountain anisotropy? Extrusion fountains have been associated with enhancers across several species [29–32], and enhancer-associated chromatin has been implicated in facilitated cohesin loading. Consistent with this connection, *Danio rerio* fountains [30] containing at least one annotated enhancer within ±50 kb of their base exhibit stronger observed/expected Hi-C signal than fountains lacking nearby enhancers (Fig. 5a,b).

**Figure 5:**
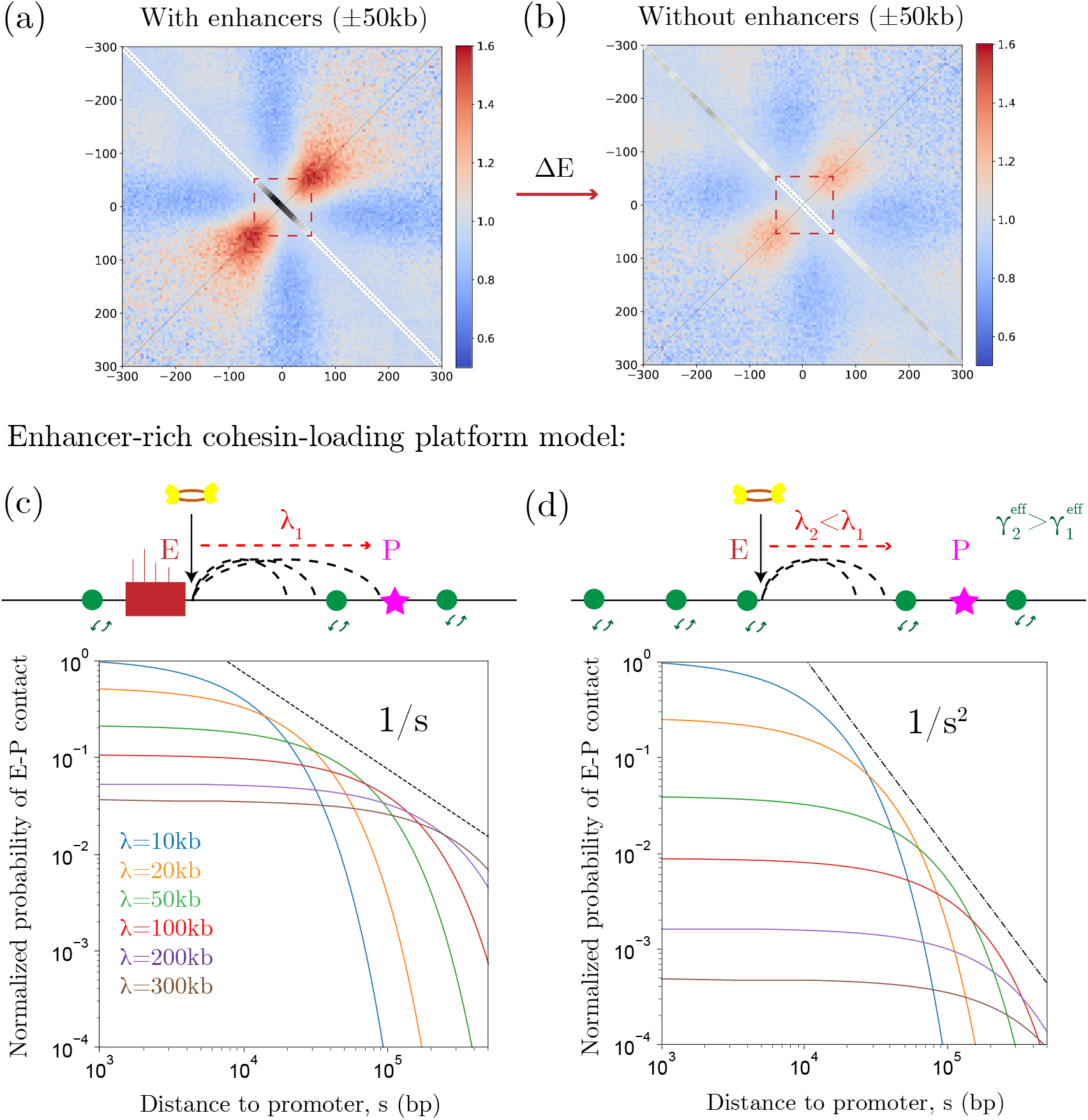
Enhancer-rich loading regions strengthen fountains and may facilitate promoter capture. (a,b) Aggregated *Danio rerio* fountains stratified by the presence of enhancers within ±50 kb of the fountain base. Fountains with nearby enhancers show stronger observed/expected Hi-C signal than those without, consistent with enhanced cohesin loading in enhancer-rich regions. Together with Figs. S10– S11, these data support a distributed enhancer-rich loading region rather than a single localized loading site. (c) Platform-assisted enhancer–promoter search. A cohesin loaded near the edge of an enhancer-rich platform (E) is retained on the platform-facing side while the opposite arm extrudes toward a promoter at genomic distance *s*. Optimizing over the mean extrusion length *λ* gives a promoter-capture envelope scaling as *s*^−1^. (d) For an isolated enhancer, productive capture additionally requires an arresting obstacle on the opposite side. This extra configuration weight yields the steeper optimal envelope *s*^−2^ in the minimal model. Thus, distributed loading and retention within an enhancer-rich platform provide a more favourable distance dependence for promoter search.

Our analysis further suggests that fountains arise from broader enhancer-rich regions rather than individual enhancers. Realigning fountains to the nearest annotated enhancer weakens rather than sharpens the aggregate signal (Fig. S10a,b), indicating genuine offsets between enhancer positions and fountain bases. Fountain bases also commonly contain multiple enhancers within a 100-kb window (Fig. S11a–e), while the mean enhancer density is well described by a Gaussian with 2*σ*_enh_ ≈ 50 kb and returns to background by 4*σ*_enh_ ≈ 100 kb (Fig. 5a,b and Fig. S10c). Finally, mean fountain strength increases with the number of nearby enhancers (Fig. S11f; *R*^2^ = 0.94 for the group means), with the strongest fountains containing, on average, eight to nine enhancers.

Notably, the observed enhancer width, 2*σ*_enh_ ≈ 48 ± 2 kb (Fig. S10c), is comparable to the cohesin-enriched loading-region width, 2*σ*_*p*_ ≈ 74 ±16 kb, inferred independently from the numerical fountain fit in *Danio rerio* (Table S7). This correspondence is consistent with enhancer enrichment marking the spatial extent of enhanced cohesin loading.

Motivated by these observations, we propose an enhancer-rich cohesin-loading platform (Fig. 5c), refining the single-site loading mechanism proposed previously [29, 30]. Repeated loading across multiple enhancers would raise local cohesin density and the frequency of cohesin–cohesin encounters, driving arm coordination toward the crowding-dominated limit *ρ* → 1/4. The resulting cohesin-rich region could also act as a dynamic barrier: for a cohesin loaded near its edge, the platform-facing arm may be repeatedly stalled by neighbouring cohesins while the opposite arm extrudes outward.

This organization may facilitate contacts with surrounding promoters (Fig. 5d). For a promoter at genomic distance *s*, retaining one arm within the platform while the other extrudes outward gives the endpoint density

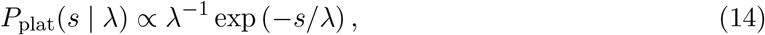

where *λ* is the mean outward extrusion length. For fixed *s*, this is maximal at *λ* = *s*, yielding

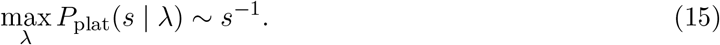

For an isolated enhancer, by contrast, productive enhancer–promoter capture additionally requires an arresting obstacle on the side opposite to the promoter. The weight of such configurations is proportional to *γ*_eff_. In the strong-disorder regime, *λ* ~ *l*_*p*_*/γ*_eff_ (Eq. (3)), so *γ*_eff_ ~ *l*_*p*_*/λ* and

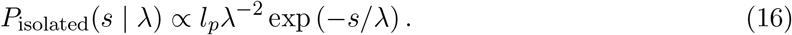

The optimum occurs at *λ* = *s/*2, giving

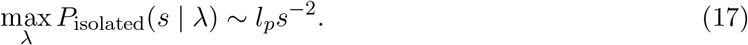

Thus, within this minimal model, distributed loading and retention within an enhancer-rich platform change the optimized promoter-capture envelope from *s*^−2^ for an isolated enhancer, which requires an additional arrest event, to *s*^−1^ for platform-mediated retention. Enhancer-rich loading regions could therefore both generate the locally crowded regime underlying fountains and provide a distributed retention zone favouring outward extrusion toward surrounding promoters.

## Discussion

We developed a kinetic theory connecting loop extrusion on chromatin to three experimentally accessible observables: the mean loop length, the full loop-length distribution, and coordination between two cohesin arms. Diverse sources of kinetic disorder enter the mean loop length through a single effective obstacle density that renormalizes the bare processivity *l*_*p*_ (Eq. (3)). This universality also exposes a limitation of the mean as a mechanistic observable: distinct obstacle landscapes and extrusion symmetries can generate the same *λ* after renormalization by *γ*_eff_ */k*. Resolving the underlying extrusion dynamics therefore requires information beyond the mean.

The full loop-length distribution breaks this degeneracy. One-sided extrusion yields a single exponential distribution regardless of the number, density, or lifetime of transient roadblocks (Eq. (6)). Effectively two-sided extrusion, by contrast, allows loop growth to continue when one arm is blocked, creating multiple growing states and hence a finite superposition of exponential modes that can develop a maximum at nonzero loop length (Eq. (7)). Peaked distributions in RAD21 ChIA-PET and in CTCF-directed ChIA-PET and MNase HiChIP are consistent with the corresponding two-sided predictions across three cell types (Fig. 2 and Fig. S12). Together, these complementary observations support effective two-sided loop extrusion in living cells.

This conclusion concerns the time-integrated symmetry of extrusion and does not require simultaneous motion of both arms. Early single-molecule experiments reported symmetric or bidirectional cohesin extrusion *in vitro* [11, 12], *whereas more recent single-molecule and in vivo* studies indicate that extrusion can be instantaneously asymmetric and switch direction during a cohesin residence time, potentially in connection with NIPBL turnover or encounters with CTCF [14, 15]. Regulator-resolved models likewise predict intermittent extrusion controlled by cohesin-associated factors [16]. Our theory coarse-grains these microscopic states into effective kinetic rates and therefore probes the accumulated contribution of the two genomic directions over a cohesin residence time. Rapid directional switching can consequently produce effectively two-sided loop growth even when extrusion is asymmetric at any given instant.

For effectively two-sided extrusion, the correlation *ρ* between accumulated arm extensions provides an additional dynamical readout of the obstacle landscape. Even when the local blocking histories of the two arms become statistically independent, they remain coupled through the shared cohesin residence time. This produces the particularly simple result that pure cohesin crowding drives *ρ* toward, but not below, the characteristic value 1/4. More persistent chromatin roadblocks can reduce the correlation further by arresting one arm for a substantial fraction of the cohesin lifetime. Arm coordination is therefore governed not simply by obstacle abundance, but by the relative abundance and persistence of different obstacle classes.

The value *ρ* = 1/4 is not unique to crowding. In the minimal alternating-arm model of Supplementary Section 4.4, symmetric Poisson switching alone yields *ρ* = 1/4 at *wτ* = 7/6, corresponding to approximately 1.17 switches per cohesin residence time. Slower switching strengthens competition between the mutually exclusive active states and can reduce the correlation below 1/4, and even below zero. Cohesin crowding therefore provides a natural environment-driven mechanism for partial desynchronization that does not require intrinsic arm switching, while remaining compatible with sufficiently rapid switching dynamics.

Fountain geometry makes this otherwise hidden arm correlation accessible in chromosome-contact maps. Full kinetic-model fits to extrusion fountains from medaka fish, western clawed frog, and zebrafish yield *ρ*_num_ ≈ 0.27–0.28, close to the crowding limit *ρ* = 1/4 (Fig. 4c), while independent Gaussian estimates from fountain ellipticity give consistent values. The same fits require strong local cohesin accumulation, showing that cohesin–cohesin encounters are quantitatively sufficient to account for both the observed degree of arm desynchronization and the resulting fountain geometry. The inferred correlation does not uniquely identify crowding as the microscopic mechanism, but establishes a quantitative connection between fountain anisotropy and cohesin-arm coordination.

Our enhancer analysis in *Danio rerio* suggests a genomic environment capable of generating this locally crowded state. Fountain bases occupy broad enhancer-rich regions rather than generally coinciding with individual enhancers, and fountain strength increases with the number of surrounding enhancers (Fig. S11). The enhancer-enrichment width, 2*σ*_enh_ ≈ 50 kb, is comparable to the cohesin-enriched loading width inferred independently from the *Danio rerio* fountain, 2*σ*_*p*_ ≈ 74 ± 16 kb, while the inferred loading regions across the three species span approximately 30–100 kb. Notably, the broad spatial extent of the loading zone is also qualitatively reflected in the broad bases of the fountains across species. (Fig. 4c). Active enhancers may recruit the extrusion machinery because enhancer-bound transcription factors interact with the cohesin-loading complex NIPBL–MAU2 [41–44]. Such enrichment could affect not only cohesin loading but also extrusion kinetics. PDS5 can terminate loop extrusion by promoting NIPBL dissociation from cohesin [45], while quantitative cofactor perturbations show that the balance between NIPBL and PDS5 dosage tunes the extrusion speed [46]. An NIPBL-rich environment could therefore increase *l*_*p*_ = *vτ* through altered cohesin residence time, extrusion speed, or both. Consistent with this possibility, the fountain fits require *l*_*p*_ ≈ 1.8–3.5 Mb, substantially above the genome-wide estimate of approximately 0.4 Mb (Tables S1,S7). We therefore propose that fountain bases correspond to distributed, enhancer-rich cohesin-loading platforms rather than uniquely positioned loading sites (Fig. 5c).

This distributed-platform picture also addresses two limitations of focal enhancer loading highlighted by recent work of Anderson *et al*. [47]. First, strong cohesin loading at a single enhancer was found to repress transcription of its distal target promoter, arguing against individual enhancers acting as strong focal loading sites. Second, loading at an isolated enhancer does not by itself retain the extruder: to support long-range contacts, one side of the complex must remain near the enhancer while the other extrudes toward the target (Fig. 5d). An enhancer-rich platform offers a way around both constraints. Distributing loading across a broader region avoids requiring strong cohesin recruitment at any single enhancer, while repeated cohesin–cohesin encounters within the platform provide a collective mechanism for retaining the platform-facing arm near an enhancer.

This collective retention can bias cohesins loaded near the platform edge toward outward extrusion: the inward-facing arm is repeatedly hindered by neighbouring cohesins, while the opposite arm continues to extrude toward surrounding promoters (Fig. 5c). Unlike a single hard barrier, the constraint is soft and renewable, because bypass or release of one neighbouring cohesin need not eliminate retention if another complex deeper within the platform provides a subsequent blocking event. In the minimal model, retaining one arm within the platform gives an optimized promoter-capture envelope proportional to *s*^−1^, whereas an isolated enhancer in Fig. 5d requires an additional arrest event and therefore yields the steeper *s*^−2^ dependence. These scalings are not complete chromosome-contact laws, which will additionally depend on three-dimensional chromatin conformation, capture kinetics, and chromatin dynamics [48–52]. Rather, they show how distributed loading and retention can increase the relative weight of regulatory contacts at larger genomic separations. Analogous principles arise in protein–DNA target search, where auxiliary binding sites and DNA looping can create “funnels” that promote rebinding to regulatory targets [53], and genomic organization itself may be shaped by selection for efficient regulatory search [54]. In this sense, enhancer-rich loading platforms could couple local cohesin retention to outward extrusion and thereby increase encounters with surrounding promoters.

Together, our results show how cohesin dynamics and roadblock kinetics shape loop extrusion across scales. The universal mean-loop law captures how chromatin disorder renormalizes intrinsic motor processivity, loop-length statistics report effective extrusion symmetry, and fountain geometry reveals a partially desynchronized arm state consistent with strong local cohesin crowding. The proposed enhancer-rich loading platform links this physical regime to regulatory genome organization and suggests how distributed cohesin loading may facilitate outward promoter search.

## Methods

### Stochastic simulations of loop extrusion

To validate the mean-field assumptions of the theory, we implemented a stochastic time-resolved simulator of loop extrusion on a one-dimensional chromatin substrate.

The simulation was parameterized by the extrusion mode *k*, the cohesin density *γ*, the roadblock density *γ*_*b*_, and the relative roadblock turnover rate *α* = *τ/τ*_*b*_. We considered both one-sided (*k* = 1) and symmetric two-sided (*k* = 2) extrusion and independently varied *γ, γ*_*b*_, and *α*. The intrinsic extrusion parameters were kept fixed at *v* = 1 and *τ* = 1, corresponding to processivity *l*_*p*_ = *vτ* = 1. Two additional quantities controlled the size of the simulation: the total number of cohesins, 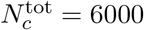, and the target mean number of cohesins simultaneously bound to chromatin, *n*_*c*_ = 200. The total number of cohesins, *N*_*c*_ was kept fixed for numerical convenience, because the loading times, residence times, and loading positions of all cohesins were generated before the start of each simulation. By contrast, *n*_*c*_ sets the characteristic number of simultaneously present and potentially interacting cohesins and therefore controls the instantaneous system size. These two quantities were used to determine the total simulation time and size of chromatin fragment,

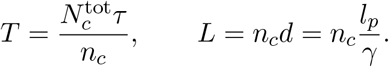

The time step was fixed at Δ*t* = 0.005.

Loading times 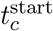 were sampled independently and uniformly over the total simulation time *T*. Cohesin residence times 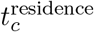 were sampled from an exponential distribution with mean *τ*. The cohesin loading coordinates *s*_*c*_ were sampled uniformly over an interval of length *L*.

Transient static roadblocks were initialized in the same manner. Their mean genomic spacing and residence time were defined as

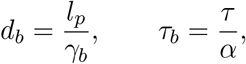

respectively. Roadblock binding times 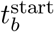 were sampled uniformly over the simulation interval, their residence times 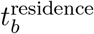 from an exponential distribution with mean *τ*_*b*_, and their genomic coordinates *s*_*b*_ uniformly over [0, *L*]. Roadblocks remained immobile at their sampled genomic positions throughout their residence times.

At each simulation step, we first identified all cohesins and roadblocks that were active at the current time, i.e. objects satisfying 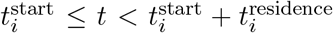. Active cohesins and roadblocks were then combined into a single list. For each cohesin, the current positions of the right and left loop edges were calculated as

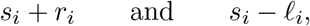

respectively, where *r*_*i*_ and *ℓ*_*i*_ are the accumulated extensions of its two arms at time *t*.

Each active arm then attempted to advance by

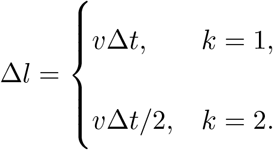

For bidirectional extrusion both arms were active, for one-sided extrusion active arm was chosen randomly for each cohesin before the start of simulation.

For a right-moving arm, objects located to the right of the cohesin were examined sequentially; analogously, objects to the left were examined for a left-moving arm. The attempted displacement was rejected if the moving loop edge encountered either edge of another active cohesin loop or the genomic position of an active roadblock. Collision detection was performed within a numerical tolerance of 2Δ*l* around the current position of the moving loop edge. Otherwise, the corresponding arm length was increased by Δ*l*. Thus, a blocked arm remained stationary while the blocking object was present and could resume extrusion at subsequent time steps after the constraint disappeared.

To collect stationary loop statistics, sampling was started at *t* = *T/*2. At each subsequent time step, the left and right arm extensions, *ℓ*_*i*_ and *r*_*i*_, of all active cohesins were recorded, together with the corresponding total loop lengths, *x*_*i*_ = *ℓ*_*i*_ + *r*_*i*_. The arm lengths and total loop lengths were accumulated over the remainder of the simulation. At the end of each run, these pooled values were used to construct the loop-length distributions and to calculate the mean loop length *λ*.

For each combination of model parameters, the simulation was repeated five independent times.

### Numerical calculation of fountain profiles

To interpret the shape of extrusion fountains, we used a minimal numerical model of two-sided loop extrusion initiated from a localized loading region. The model was formulated in two-arm coordinates, where the left and right extrusion arms are described by their dimensionless extensions, 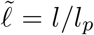 and 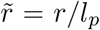. The state of an extrusion complex was represented by 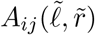, where the two indices denote the states of the left and right arms, respectively, *i, j* in {0, *b, c*}. Here, 0 denotes an actively extruding arm, *c* denotes an arm blocked by another cohesin, and *b* denotes an arm blocked by a static genomic barrier. We considered the minimal case with one static barrier class *b* and other cohesins *c*.

The stationary time-integrated probabilities satisfy the first-order balance equations (Supplementary Section 4, Eq. (131-135)).

The stationary system was solved in the first quadrant, 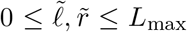, using a uniform grid 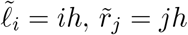, with *h* = *L*_max_*/N* and *N* = 600. The domain size was chosen as *L*_max_ = 2.

Cohesin loading at zero initial arm extension was represented by a narrow Gaussian regularization of a point source

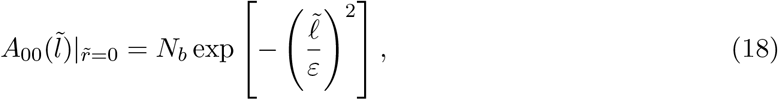

with *ε* = 10^−3^ and *A*_00_(0, 0) = *N*_*b*_. Here, *N*_*b*_ is the dimensionless amplitude of the cohesin-loading source. Because the stationary equations are linear and the resulting model contact maps were subsequently normalized, *N*_*b*_ affects only the overall signal amplitude and was set to *N*_*b*_(0) = 1 without loss of generality.

The discretization followed the characteristic directions of the transport terms. A first-order upwind finite-difference marching scheme was used: *A*_00_, whose transport operator contains derivatives with respect to both arm coordinates, was propagated along diagonal grid directions; the one-sided blocked states *A*_0*c*_ and *A*_0*b*_ were propagated along the 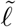 coordinate, whereas *A*_*c*0_ and *A*_*b*0_ were propagated along the 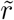 coordinate. At each grid point, the doubly blocked states were calculated locally from the algebraic closure relations (Supplementary Section 4, Eq. (136-139)).

To account for spatially distributed cohesin loading, we replaced the point-like loading site at the fountain base by a loading platform of the width *σ*_*p*_. Specifically, we assumed that cohesin can load at a position 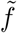 around the fountain base, with probability density 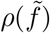. For a contact between loci located at 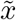 and 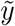 relative to the fountain base, a cohesin loaded at 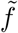 must extrude left and right arms of lengths 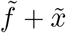 and 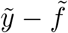, respectively. Therefore, its contribution to this contact is given by 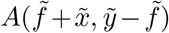, where 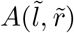 is the point-source contact kernel for a loop with arm lengths 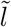 and 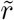.

Summing over all possible loading positions between the two interacting loci gives the fountain profile

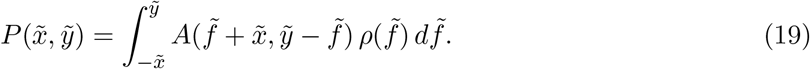

The integration limits follow from the requirement that the loading position lies between the two contacting loci, so that both loop-arm lengths are non-negative. Thus, the convolution represents a superposition of point-source fountains whose bases are distributed around the nominal fountain base.

We used a normalized Gaussian loading density centered at the fountain base,

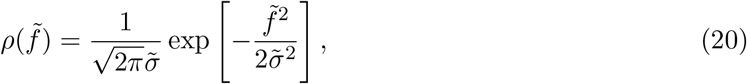

where 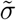 is the dimensionless width of the loading zone.

The numerical contact map was first generated in dimensionless extrusion coordinates and then rescaled to genomic units using *l*_*p*_, i.e. 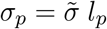.

### Aggregate fountain fitting

We analyzed loop-extrusion fountain maps using the Hi-C data and fountain annotations reported by Galitsyna et. al. [30]. The analysis was performed for three vertebrate species: *Oryzias latipes, Xenopus tropicalis*, and *Danio rerio*. Hi-C signals around fountain bases were extracted at 5-kb resolution using species-specific flanking windows of 200 kb for *Danio rerio*, 250 kb for *Xenopus tropicalis*, and 100 kb for *Oryzias latipes*, then converted to chromosome-specific observed/expected values, and averaged pixel-wise. The aggregate maps included 1885 fountains for *Oryzias latipes*, 1733 fountains for *Xenopus tropicalis*, and 1460 fountains for *Danio rerio*. The same aggregate maps were used for both the empirical Gaussian fit and the numerical extrusion-model fit.

The fit quality was quantified by the normalized root-mean-square error,

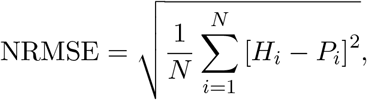

where *H*_*i*_ and *P*_*i*_ are the normalized Hi-C and model signals at pixel *i*, respectively, and the sum runs over the selected fitting window. The near-peak error, denoted NRMSE^80kb^, was computed in the central near-peak window used for fitting. The full-window error, NRMSE^full^, was computed over the complete fitting window and was used as a diagnostic of how well the model reproduced the global fountain profile.

As an empirical description of fountain geometry, we fitted each aggregate map with an anisotropic two-dimensional Gaussian,

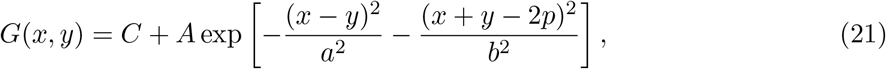

where *A* is the amplitude, *C* is the local background, *p* is the displacement of the fountain maximum from the base, and *a* and *b* are the widths along the minor and major axes of the fountain. Here *x* and *y* are the genomic coordinates of the interacting loci relative to the fountain base. The primary diagonal (*y* = *x*) is parallel to the fountain, whereas the secondary diagonal (*y* = −*x*) is perpendicular to it. Accordingly, *a* and *b* are the transverse and longitudinal widths. The Gaussian maximum is located at (*x, y*) = (*p, p*).

The fit was performed in two steps. First, a full window around the expected fountain peak was used to obtain an initial estimate of the peak position and axis widths. Second, the model was refitted in a near-peak window to reduce the influence of distal asymmetric tails and background signal. From the fitted Gaussian widths, we computed the empirical correlation coefficient using Eq.(13). This approach results in a model-light estimate of the fountain asymmetry and was used as a reference for comparison with the full numerical model.

The same aggregate maps were then fitted with the numerical extrusion model described above. For each candidate parameter set, we solved the two-arm extrusion equations numerically, convolved the resulting point-source kernel with a Gaussian loading-position distribution, rescaled the dimensionless model coordinates to genomic coordinates using processivity *l*_*p*_, and interpolated the resulting model map onto the Hi-C aggregate grid. As in the Gaussian fitting procedure, the resulting model profile *P* (*x, y*) was compared with the experimental map after the rescaling with offset, *A* + *C* · *P* (*x, y*), where the offset *A* and amplitude *C* were optimized by least-squares minimization for each candidate parameter set. The fitted parameters were *l*_*p*_, the dimensionless loading-zone width 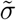, and the cohesin density *γ*_*c*_. The barrier density *γ*_*b*_ was fixed to 1.8, based on the HeLa estimation (Table S2), and all fits shown here were performed at *α* = 0.

Model parameters were optimized using a bounded coordinate-descent procedure. The search was initialized at *l*_*p*_ = 500kb, 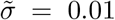, and *γ*_*c*_ = 5, with parameter bounds 100kb ≤ *l*_*p*_ ≤ 10000kb, 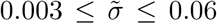, and 0 ≤ *γ*_*c*_ ≤ 40. The initial coordinate steps were 300 kb, 0.001, and 1 for *l*_*p*_, 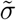, and *γ*_*c*_, respectively. At each iteration, the current parameter set and the two neighboring points obtained by adding or subtracting the corresponding step along each parameter axis were evaluated, while the remaining parameters were kept fixed. Candidate values outside the prescribed bounds were clipped to the nearest boundary.

The search was initialized at *l*_*p*_ = 500kb, 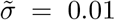, and *γ*_*c*_ = 5, with parameter bounds 100kb ≤ *l*_*p*_ ≤ 10000kb, 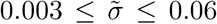, and 0 ≤ *γ*_*c*_ ≤ 40. The initial coordinate steps were 300 kb, 0.001, and 1 for *l*_*p*_, 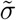, and *γ*_*c*_, respectively. At each iteration, the current parameter set and the two neighboring points obtained by adding or subtracting the corresponding step along each parameter axis were evaluated, while the remaining parameters were kept fixed. Candidate values outside the prescribed bounds were clipped to the nearest boundary. The near-peak error, denoted NRMSE^80 kb^, was computed in the central 80-kb near-peak window used for the fitting. The full-window error, NRMSE^full^, was computed over the complete fitting window and was used as a diagnostic of how well the model reproduced the global fountain profile. If none of the coordinate moves produced sufficient improvement, all step sizes were multiplied by 0.5. Optimization was terminated when the steps reached 10^−4^ for 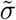, and 0.1 for *γ*_*c*_. The same initialization, bounds, and convergence criteria were used for all aggregate maps.

### Enhancer annotations and fountain-associated enhancer analysis

Active enhancer annotations for the *Danio rerio* Dome stage were obtained from the PADRE/ChromHMM annotation of [55]. Only regions assigned to the `5_EnhA1` state, classified in the original annotation as active enhancer chromatin, were retained. Fountain-base coordinates and fountain scores were obtained from [30].

For each fountain, we counted the number of enhancers located within 50 kb of the inferred fountain base. Fountains were divided according to the presence or absence of at least one nearby enhancer, and their Hi-C maps were aligned to the fountain bases and averaged (Fig. 5a,b) in same window, as described in Sec. *Aggregate fountain fitting*.

To determine whether individual enhancers precisely mark fountain bases, the same enhancer-associated fountains were realigned to the midpoint of the nearest enhancer interval and reaggregated (Fig. S10a,b). The distribution of enhancer positions relative to fountain bases was approximately Gaussian, with a width of *σ*_enh_ = 24 ± 1 kb (2*σ*_enh_ ≈ 50 kb), indicating that enhancer enrichment extends over a finite region around the fountain base (Fig. S10c). Finally, fountains were grouped according to the number of enhancer intervals located within 50 kb of the base. Mean fountain scores and their standard deviations were calculated for each group, and linear regression was performed using the group means (Fig. S11).

### Loop-length distributions from experimental data

Five experimental datasets were used to construct loop-length distributions: three CTCF ChIA-PET datasets, one RAD21 ChIA-PET dataset, and one CTCF MNase HiChIP dataset.

#### ChIA-PET datasets and preprocessing

CTCF-associated chromatin interactions were analysed in K562, GM12878, and HeLa-S3 cells [37], and RAD21-associated interactions were analysed in GM12878 cells (dataset contained 2 replicates) [36].

For all datasets, processed interactions were converted to a common representation containing the genomic coordinates of the two anchors (the start and end of corresponding fragment for each anchor) and the number of supporting paired-end tags (PET count). Only intrachromosomal interactions were retained. The genomic loop length was defined as the distance between centres of anchors.

To construct an empirical distribution of cohesin-associated loop lengths, each interaction was weighted by its PET count, so that the contact length supported by *n* PETs contributed *n* observations to the distribution. Where multiple replicates were available, the corresponding PET-weighted distributions were pooled.

After this preprocessing, the resulting ensembles contained 90 824 observations for K562 CTCF, 1 712 596 for GM12878 CTCF, 299 579 for HeLa CTCF, and 155 049 total in two replicas for GM12878 RAD21.

#### MNase HiChIP dataset and preprocessing

The independent K562 CTCF dataset was obtained from MNase CTCF HiChIP [38]. Four libraries were generated from K562 cells by MNase digestion, CTCF immunoprecipitation, proximity ligation, and paired-end sequencing. We used the fragment-level interaction dataset corresponding to CTCF-associated fragments with an *extended fragment end*. In the original analysis, Sept et al. identified a downstream extension of short TF-protected fragments (fragment length *<* 120 bp), extending approximately 15 bp beyond the CTCF footprint, as a signature of cohesin occupancy at CTCF-bound sites; this extension was strongly attenuated upon acute RAD21 depletion [38]. For the identification of CTCF/cohesin-associated fragments, we followed the original fragment-selection procedure implemented by the authors and used their analysis code available from the GitHub repository (https://github.com/aryeelab/cohesin_extrusion_reproducibility).

Sept et al. reported a bimodal interaction-length distribution for short CTCF-associated fragments, comprising proximal and long-range contact populations, and showed that fragments with an extended end (cohesin-associated) are preferentially enriched in the long-range component (Fig. 4H in [38]). The original study operationally defined long-range interactions as those exceeding 10 kb, however, a pronounced increase in the proximal-contact population became apparent below approximately 30 kb. To restrict the analysis to the long-range cohesin-associated component, we therefore excluded interactions shorter than 30 kb before normalization. The final distribution contained 150 214 interactions.

### Comparison with theoretical loop-length distributions

Experimental loop-length distributions were compared with the analytical distributions derived for the two-sided loop-extrusion model (Fig. S12a). The model describes several loop states depending on the interaction of extruding cohesins with barriers. CTCF datasets were compared with the doubly barrier-blocked distribution, *A*_*bb*_, whereas the RAD21 distribution was compared with the total cohesin-associated loop-length distribution, *A* = *A*_00_ + *A*_*b*_ + *A*_*c*_ + *A*_*bb*_ + *A*_*bc*_ + *A*_*cc*_.

The theoretical distributions depend on the dimensionless cohesin density *γ*_*c*_, the barrier density *γ*_*b*_, and the parameter *α*. Exploration of the full parameter space revealed a strong parameter degeneracy: substantially different parameter combinations can generate very similar loop-length distributions (Fig. S12d–f).

To reduce this degeneracy, *α* was fixed at *α* = 1.2, corresponding to the HeLa estimates for CTCF (Table S2), and the dependence of the agreement between experiment and theory on *γ*_*c*_ and 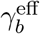 was examined. The parameter grid covered *γ*_*c*_ ≃ 0.1–9.9 and 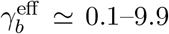, with a step of 0.2. For each point of the parameter grid, the theoretical and experimental loop-length distribution were binned using 10-kb binning and then compared with each other using the normalized mean absolute error (NMAE),

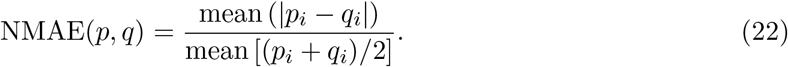

Here, *p*_*i*_ and *q*_*i*_ denote the normalized values of the experimental and theoretical loop-length distributions, respectively, in the *i*-th bin of the loop-length grid, and the mean is taken over all bins included in the comparison. The use of an absolute rather than squared error metric reduces the influence of isolated high-amplitude fluctuations in the experimental histograms.

For cell lines represented by two independent experimental measurements, a single cell-line score was defined as the arithmetic mean of the corresponding NMAE values. For GM12878,

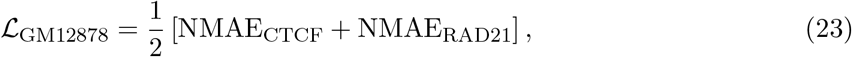

where the CTCF distribution was compared with *A*_*bb*_ and the RAD21 distribution with *A*. For K562,

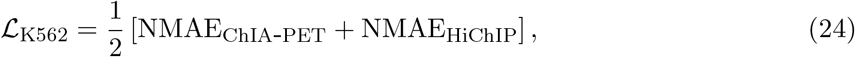

with both CTCF-centred measurements compared with *A*_*bb*_. For HeLa, only the CTCF ChIA-PET distribution was used.

The resulting NMAE landscapes are shown in Fig. S12(d–f). Because of the broad low-error regions produced by the parameter degeneracy, these landscapes were not interpreted as providing unique estimates of *γ*_*c*_ and 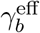. Instead, they were used to test the consistency of the experimental loop-length distributions with parameters estimated independently for HeLa cells in Tables S1,S2. The independently estimated parameter values and their uncertainty region are indicated in Fig. S12(d–f) by a cross and a pink ellipse, respectively.

## Data availability

No new experimental data were generated in this study. Experimental measurements used for model parameterization were compiled from previously published studies. The HeLa WT parameter set, derived from FRAP, fluorescence microscopy, mass spectrometry, and Hi-C measurements, was compiled from [6, 12, 24, 34, 56–61]. Parameters describing MCM abundance, spacing, and barrier activity were obtained from [10, 60]. Measurements for WAPL-depleted cells, including cohesin residence times and loop-size estimates together with their matched WT controls, were compiled from [31, 34, 62–66]. More detailed information on the used parameters is provided in Supplementary Tables S1 and S3.

Five publicly available chromatin-interaction datasets were used to construct the experimental loop-length distributions: three CTCF ChIA-PET datasets, one RAD21 ChIA-PET dataset, and one CTCF MNase HiChIP dataset. CTCF-associated interactions were analysed in K562 cells (GSM970216; GSE39495), GM12878 cells (GSM1872886; GSE72816), and HeLa cells (GSM1872888; GSE72816) [37]. RAD21-associated interactions were analysed in GM12878 cells using the two-replicate dataset GSM1436265 (GSE59395) [36]. K562 CTCF MNase HiChIP data were obtained from GEO accession GSE285087 [38]. The corresponding preprocessing and loop-selection procedures are described in the Methods and Supplementary Information.

The zebrafish (*Danio rerio*) Hi-C data and extrusion-fountain annotations from Galitsyna et al. were obtained from GEO accession GSE195609 [30]. Zebrafish enhancer annotations used in the analysis, including PADREs active at the Dome developmental stage, were obtained from the DANIO-CODE Data Coordination Center (https://danio-code.zfin.org) [55]. For the analyses of *Oryzias latipes* and *Xenopus tropicalis*, we used Hi-C data originally generated experimentally by Nakamura et al. [39] and Niu et al. [40], respectively, and subsequently re-analysed for extrusion-fountain detection by Galitsyna et al. [30]. The resulting fountain annotations and intermediate processed datasets for these species are available in the Galitsyna et al. supplementary datasets and OSF repository (https://osf.io/mt4vf).

Processed data generated in this study, including the loop-length distributions, observed/expected fountain aggregates, Gaussian and kinetic-model fits, inferred arm–arm correlations, fitted model parameters, enhancer-density profiles, and numerical simulation results underlying the figures and tables, are provided in the Source Data file. Source data are provided with this paper.

## Code availability

Custom Python code used for stochastic loop-extrusion simulations, numerical solution of the two-arm kinetic model, construction of aggregate Hi-C “fountains”, Gaussian and analytical model fitting, inference of arm–arm correlations and enhancer analysis is publicly available at https://github.com/melamory1blimm/Loop-extrusion and has been archived in Zenodo at https://doi.org/10.5281/zenodo.22028319.

## Author contributions

K.P. designed the research; A.C. and K.P. performed the analysis; A.C., M.G., R.M. and K.P. wrote the paper. All authors have reviewed and approved the manuscript.

## Acknowledgements

We gratefully acknowledge M. Kardar, L. Mirny, and P. Virnau for illuminating discussions and A. Cherstvy and A. Goychuk for valuable comments on the original form of the manuscript. The work of A.C. and K.P. was supported by the Russian Science Foundation (Grant No. 25-13-00277). R.M. acknowledges support from DFG (CRC Data assimilation, Grant 318763901). K.P. also acknowledges support from the Alexander von Humboldt Foundation.

## References

[1] J. Dekker and L. Mirny. The chromosome folding problem and how cells solve it. Cell, 187 (23):6424–6450, 2024.

[2] P. Fraser and W. Bickmore. Nuclear organization of the genome and the potential for gene regulation. Nature, 447(7143):413–417, 2007.

[3] A. L. Sanborn, S. S. Rao, S. C. Huang, N. C. Durand, M. H. Huntley, A. I. Jewett, et al. Chromatin extrusion explains key features of loop and domain formation in wild-type and engineered genomes. Proceedings of the National Academy of Sciences of the United States of America, 112(47):E6456–E6465, 2015.

[4] G. Fudenberg, M. Imakaev, C. Lu, A. Goloborodko, N. Abdennur, and L.A. Mirny. Formation of chromosomal domains by loop extrusion. Cell Reports, 15(9):2038–2049, 2016.

[5] C. A. Brackley, J. Johnson, D. Michieletto, A. N. Morozov, M. Nicodemi, P. R. Cook, and D. Marenduzzo. Nonequilibrium chromosome looping via molecular slip links. Physical Review Letters, 119(13):138101, 2017.

[6] S. S. Rao, S. C. Huang, B. G. St Hilaire, J. M. Engreitz, E. M. Perez, et al. Cohesin loss eliminates all loop domains. Cell, 171(2):305–320, 2017.

[7] E. P. Nora et al. Molecular basis of CTCF binding polarity in genome folding. Nature Communications, 11(1):5612, 2020.

[8] E. de Wit et al. CTCF binding polarity determines chromatin looping. Molecular Cell, 60(4): 676–684, 2015.

[9] H. B. Brandão, P. Paul, A. A. van den Berg, D. Z. Rudner, X. Wang, and L. A. Mirny. RNA polymerases as moving barriers to condensin loop extrusion. Proceedings of the National Academy of Sciences of the United States of America, 116(41):20489–20499, 2019.

[10] B. J. Dequeker et al. MCM complexes are barriers that restrict cohesin-mediated loop extrusion. Nature, 606(7912):197–203, 2022.

[11] Y. Kim, Z. Shi, H. Zhang, I. J. Finkelstein, and H. Yu. Human cohesin compacts DNA by loop extrusion. Science, 366(6471):1345–1349, 2019.

[12] I. F. Davidson, B. Bauer, D. Goetz, W. Tang, G. Wutz, and J. M. Peters. DNA loop extrusion by human cohesin. Science, 366(6471):1338–1345, 2019.

[13] M. Ganji et al. Real-time imaging of dna loop extrusion by condensin. Science, 360(6384): 102–105, 2018.

[14] R. Barth et al. SMC motor proteins extrude DNA asymmetrically and can switch directions. Cell, 188(3):749–763, 2025.

[15] P. Wang et al. Cohesin extrudes chromatin loop unidirectionally through two modes of mechanisms in human cells. bioRxiv, doi: 10.64898/2026.05.01.722300, 2026.

[16] M. M. Tortora and G. Fudenberg. The physical chemistry of interphase loop extrusion. Cell Genomics, 6(3):101098, 2026.

[17] S. K. Nomidis, E. Carlon, S. Gruber, and J. F. Marko. DNA tension-modulated translocation and loop extrusion by SMC complexes revealed by molecular dynamics simulations. Nucleic Acids Research, 50(9):4974–4987, 2022.

[18] R. Takaki, A. Dey, G. Shi, and D. Thirumalai. Theory and simulations of condensin mediated loop extrusion in DNA. Nature Communications, 12(1):5865, 2021.

[19] H. Salari and D. Jost. Active loop extrusion modulates the mechanical response of chromatin under tension. Physical Review Research, 8(1):013188, 2026.

[20] Edward J Banigan, Aafke A van den Berg, Hugo B Brandão, John F Marko, and Leonid A Mirny. Chromosome organization by one-sided and two-sided loop extrusion. eLife, 9:e53558, 2020.

[21] B. Chan and M. Rubinstein. Theory of chromatin organization maintained by active loop extrusion. Proceedings of the National Academy of Sciences of the United States of America, 120(23):e2222078120, 2023.

[22] A. J. Pinto, B. Pradhan, D. Tetiker, M. P. Schmitt, E. Kim, and P. Virnau. SMC motor proteins operate at the near-minimal forces for DNA loop extrusion. bioRxiv: 10.64898/2026.03.05.709531, 2026.

[23] B. Chan and M. Rubinstein. Activity-driven chromatin organization during interphase: Compaction, segregation, and entanglement suppression. Proceedings of the National Academy of Sciences of the United States of America, 121(21):e2401494121, 2024.

[24] K. Polovnikov and D. Starkov. A universal polymer signature in Hi-C resolves cohesin loop density and supports monomeric extrusion. Proceedings of the National Academy of Sciences of the United States of America, 123(16):e2534385123, 2026.

[25] K. E. Polovnikov, H. B. Brandão, S. Belan, B. Slavov, M. Imakaev, and L. A. Mirny. Crumpled polymer with loops recapitulates key features of chromosome organization. Physical Review X, 13(4):041029, 2023.

[26] K. E. Polovnikov and B. Slavov. Topological and nontopological mechanisms of loop formation in chromosomes: Effects on the contact probability. Physical Review E, 107(5):054135, 2023.

[27] B. Slavov and K. Polovnikov. Intrachain distances in a crumpled polymer with random loops. JETP Letters, 118(3):208–214, 2023.

[28] S. Belan and V. Parfenyev. Footprints of loop extrusion in statistics of intra-chromosomal distances: An analytically solvable model. The Journal of Chemical Physics, 160(12):124901, 2024.

[29] Y. Guo et al. Chromatin jets define the properties of cohesin-driven in vivo loop extrusion. Molecular Cell, 82(20):3769–3780, 2022.

[30] A. Galitsyna et al. Extrusion fountains are hallmarks of chromosome organization emerging upon zygotic genome activation. Nature Communications, 17(1):2787, 2026.

[31] N. Q. Liu et al. Extrusion fountains are restricted by WAPL-dependent cohesin release and CTCF barriers. Nucleic Acids Research, 53(12):gkaf549, 2025.

[32] J. Kim, H. Wang, and S. Ercan. Cohesin organizes 3D DNA contacts surrounding active enhancers in C. elegans. Genome Research, 35(5):1108–1123, 2025.

[33] G. Fudenberg and M. Imakaev. FISH-ing for captured contacts: towards reconciling FISH and 3C. Nature Methods, 14(7):673–678, 2017.

[34] G. Wutz et al. Topologically associating domains and chromatin loops depend on cohesin and are regulated by CTCF, WAPL, and PDS5 proteins. The EMBO Journal, 36(24):3573–3599, 2017.

[35] A. Tedeschi et al. WAPL is an essential regulator of chromatin structure and chromosome segregation. Nature, 501(564–568):204904, 2013.

[36] N. Heidari et al. Genome-wide map of regulatory interactions in the human genome. Genome Research, 24(12):1905–1917, 2014.

[37] Z. Tang et al. CTCF-mediated human 3D genome architecture reveals chromatin topology for transcription. Cell, 163(7):1611–1627, 2015.

[38] Corriene E Sept, Y Esther Tak, Viraat Goel, Mital S Bhakta, Christian G Cerda-Smith, Haley M Hutchinson, Marco Blanchette, Christine E Eyler, Sarah E Johnstone, J Keith Joung, et al. High-resolution CTCF footprinting reveals impact of chromatin state on cohesin extrusion. Nature Communications, 16(1):4506, 2025.

[39] R. Nakamura et al. CTCF looping is established during gastrulation in medaka embryos. Genome Research, 31(6):968–980, 2021.

[40] L. Niu et al. Three-dimensional folding dynamics of the Xenopus tropicalis genome. Nature Genetics, 53(7):1075–1087, 2021.

[41] M. H. Kagey et al. Mediator and cohesin connect gene expression and chromatin architecture. Nature, 467(7314):430–435, 2010.

[42] Y. Zhu, M. Denholtz, H. Lu, and C. Murre. Calcium signaling instructs NIPBL recruitment at active enhancers and promoters via distinct mechanisms to reconstruct genome compartmentalization. Genes & development, 35:1–17, 2020.

[43] L. Rinaldi et al. The glucocorticoid receptor associates with the cohesin loader NIPBL to promote long-range gene regulation. Science Advances, 8(13):eabj8360, 2022.

[44] N. J. Rinzema et al. Building regulatory landscapes reveals that an enhancer can recruit cohesin to create contact domains, engage CTCF sites and activate distant genes. Nature Structural & Molecular Biology, 29(6):563–574, 2022.

[45] G. Wutz et al. PDS5 proteins control genome architecture by limiting the lifetime of cohesin-NIPBL complexes. Molecular Cell, 86(9):1614–1634, 2026.

[46] R. Shah et al. Cohesin cofactor dosage sets the rate of loop extrusion, rendering genome folding tunable yet vulnerable to genetic disruption. Molecular Cell, 86(9):1635–1652, 2026.

[47] E. C. Anderson et al. The role of cohesin loading at enhancers in the flux of loop extrusion and long-range transcriptional control. bioRxiv, 2026.

[48] K. Polovnikov and M. Kardar. Universal center-of-mass scaling shapes segment fluctuations and chromatin dynamics. Physical Review Research, 7(4):043104, 2025.

[49] K.E Polovnikov, M. Gherardi, M. Cosentino-Lagomarsino, and M.V. Tamm. Fractal folding and medium viscoelasticity contribute jointly to chromosome dynamics. Physical Review Letters, 120(8):088101, 2018.

[50] D. Grebenkov, R. Metzler, and G. Oshanin. Target search problems. Springer Nature Switzerland, 2024.

[51] L. Hedström, R. Metzler, and L. Lizana. Enhancer-insulator pairing reveals heterogeneous dynamics in long-distance 3D gene regulation. PRX Life, 2(3):033008, 2024.

[52] M. V. Tamm and K. Polovnikov. Dynamics of Polymers: Classic Results and Recent Developments. Order, Disorder and Criticality: Advanced Problems of Phase Transition Theory. World Scientific, 2018.

[53] M. Bauer, E. S. Rasmussen, M. A. Lomholt, and R. Metzler. Real sequence effects on the search dynamics of transcription factors on DNA. Scientific Reports, 5(1):10072, 2015.

[54] G. Kolesov, Z. Wunderlich, O. N. Laikova, M. S. Gelfand, and L. A. Mirny. How gene order is influenced by the biophysics of transcription regulation. Proceedings of the National Academy of Sciences, 104(35):13948–13953, 2007.

[55] Damir Baranasic, Matthias Hörtenhuber, Piotr J Balwierz, Tobias Zehnder, Abdul Kadir Mukarram, Chirag Nepal, Csilla Várnai, Yavor Hadzhiev, Ada Jimenez-Gonzalez, Nan Li, et al. Multiomic atlas with functional stratification and developmental dynamics of zebrafish cis-regulatory elements. Nature Genetics, 54(7):1037–1050, 2022.

[56] A. Brunner, N. R. Morero, W. Zhang, M. J. Hossain, M. Lampe, H. Pflaumer, et al. Quantitative imaging of loop extruders rebuilding interphase genome architecture after mitosis. Journal of Cell Biology, 224(3):e202405169, 2025.

[57] G. Wutz et al. ESCO1 and CTCF enable formation of long chromatin loops by protecting cohesinSTAG1 from WAPL. eLife, 9:e52091, 2020.

[58] J. Holzmann, A. Z. Politi, K. Nagasaka, M. Hantsche-Grininger, N. Walther, B. Koch, et al. Absolute quantification of cohesin, CTCF and their regulators in human cells. eLife, 8:e46269, 2019.

[59] Iain F Davidson, Roman Barth, Maciej Zaczek, Jaco van der Torre, Wen Tang, Kota Nagasaka, Richard Janissen, Jacob Kerssemakers, Gordana Wutz, Cees Dekker, et al. CTCF is a DNA-tension-dependent barrier to cohesin-mediated loop extrusion. Nature, 616(7958):822–827, 2023.

[60] Nils A Kulak, Garwin Pichler, Igor Paron, Nagarjuna Nagaraj, and Matthias Mann. Minimal, encapsulated proteomic-sample processing applied to copy-number estimation in eukaryotic cells. Nature Methods, 11(3):319–324, 2014.

[61] DA Jackson, P Dickinson, and PR Cook. The size of chromatin loops in HeLa cells. The EMBO Journal, 9(2):567–571, 1990.

[62] Susannah Rankin. A WAPL a day keeps the sisters apart: WAPL and cohesin dynamics. Developmental Cell, 11(6):754–755, 2006.

[63] Antonio Tedeschi, Gordana Wutz, Sébastien Huet, Markus Jaritz, Annelie Wuensche, Erika Schirghuber, Iain Finley Davidson, Wen Tang, David A Cisneros, Venugopal Bhaskara, et al. WAPL is an essential regulator of chromatin structure and chromosome segregation. Nature, 501(7468):564–568, 2013.

[64] J.H. Haarhuis, Robin van der Weide, Vincent A Blomen, J Omar Yáñez-Cuna, Mario Amendola, Marjon S van Ruiten, Peter HL Krijger, Hans Teunissen, René H Medema, Bas van Steensel, et al. The cohesin release factor WAPL restricts chromatin loop extension. Cell, 169 (4):693–707, 2017.

[65] T. H. S. Hsieh, C. Cattoglio, E. Slobodyanyuk, A. S. Hansen, X. Darzacq, and R. Tjian. Enhancer-promoter interactions and transcription are maintained upon acute loss of CTCF, cohesin, WAPL, and YY1. Nature Genetics, 54(12):1919–1932, 2022.

[66] J. Gassler, H. B. Brandão, M. Imakaev, I. M. Flyamer, S. Ladstätter, W. A. Bickmore, et al. A mechanism of cohesin-dependent loop extrusion organizes zygotic genome architecture. The EMBO Journal, 36(24):3600–3618, 2017.

