## Supplementary Materials for "Loop statistics and fountain geometry reveal effective two-sided cohesin extrusion and crowding-induced arm desynchronization"

**8 Estimation of loop-length distribution parameters** **59**

### 1. Model

#### 1.1 Main definitions

We consider a single loop-extruding cohesin as a tagged particle moving in a steady-state background of chromatin-bound obstacles and other cohesins. It loads onto chromatin at time $t = 0$ and enlarges a loop of size $x$ until stochastic dissociation with mean residence time $\tau$. During extrusion, both cohesins and obstacles stochastically bind to and unbind from chromatin, so that their mean densities remain constant in the steady state.

We denote by $v$ the total loop-growth rate. For one-sided extrusion, $v$ is the speed of the single moving arm; for symmetric two-sided extrusion, each arm advances at $v/2$, so that the total loop still grows at rate $v$. Throughout this work we focus on two limiting mechanisms: strictly one-sided extrusion, in which only one cohesin arm is mobile, and symmetric two-sided extrusion, in which both arms move simultaneously. Alternative scenarios, such as stochastic switching of the active arm or switching induced by encounters with obstacles, are discussed separately in Sec. 3.2.2.

Two basic length scales characterize the problem. The first is the intrinsic processivity,

$$l_p = v\tau,$$

i.e., the mean loop size generated in the absence of obstacles. The second is the mean extruder spacing $d$, defined as the average distance between neighboring cohesins along chromatin in the steady state.

Loop growth is interrupted by blocking events caused by a mixture of $n$ obstacle classes, $i = 1, \ldots, n$ (e.g., CTCF or transcriptional complexes), and by other extruders, treated as a special class $i = c$. In the steady state, each obstacle class $i$ is characterized by a mean spacing $d_i$ and a mean lifetime $\tau_i$. For the extruder class, we set

$$d_c \equiv d, \qquad \tau_c \equiv \tau.$$

Unless stated otherwise, cohesins are treated as opaque obstacles, i.e., mutually blocking upon encounter. In toy models or limiting cases where cohesins are allowed to pass through one another, this is stated explicitly.

We note that static barriers need not be perfectly opaque in vivo: upon encounter, a barrier may stall cohesin only with probability $p$. Such partial permeability can be absorbed into the effective spacing between blocking events, which is increased by a factor $p^{-1}$ relative to the raw genomic spacing between barrier sites. In the theoretical formulation below, we therefore treat static barriers as effectively impermeable, with $d_i$ understood as the *effective* spacing between successful blocking events. When comparing to experiment, this effective spacing is obtained by dividing the measured raw separation between barriers by the independently measured blocking probability $p$ (Table S1).

With this convention, all definitions of $d_i$, $\gamma_i$, and the corresponding mean-field equations remain unchanged.

#### 1.2 Bound states of the system

We describe the system in terms of state-resolved probability densities $A_q(x,t)$, where the set of states depends on whether extrusion is one-sided or two-sided.

**One-sided extrusion.** In this case, blocking can occur only on the moving side, when the advancing cohesin arm encounters a pre-existing obstacle. We neglect the complementary process in which an obstacle binds directly onto the cohesin, since this requires a localized coincidence in both space and time and is therefore higher order in obstacle density. The state space of chromatin-bound cohesins then consists of one freely extruding state and $n+1$ blocked states,

$$q \in \{\,0,\ i\,\},$$

where 0 denotes the mobile state and $i$ labels the blocking class (the $n$ static obstacle types plus cohesin, treated as an additional class $c$). The total number of states is therefore

$$\Gamma_{\text{uni}} = 1 + (n+1) = n+2.$$

**Two-sided extrusion.** Here the two arms may be blocked independently. If one is interested only in the total loop length, it is sufficient to distinguish whether zero, one, or two arms are blocked, without resolving left and right separately. In that reduced description, the state space is

$$q \in \{\,00,\ i,\ ij\,\},$$

where $00 \equiv 0$ denotes the fully mobile state, $i$ a state in which one of the two arms is blocked by obstacle class $i$, and $ij$ a state in which the two arms are blocked by obstacle classes $i$ and $j$. Since the pair $ij$ is unordered in this representation, the total number of states is

$$\Gamma_{\text{bi}}^{\text{red}} = 1 + (n+1) + \frac{(n+1)(n+2)}{2} = \frac{(n+2)(n+3)}{2}.$$

When the distinction between the left and right arms is required, for example to simplify intermediate calculations or to analyze arm–arm correlations, we use the fully resolved notation

$$q \in \{\,00,\ i0,\ 0i,\ ij\,\},$$

where the first (second) index refers to the left (right) arm; 0 denotes a free arm and $i, j$ label the blocking class. Thus, $A_{00}$ is the fully mobile state, $A_{i0}$ and $A_{0j}$ are partially blocked states, and $A_{ij}$ is the fully blocked state. In this representation the total number of states is

$$\Gamma_{\text{bi}}^{\text{full}} = 1 + 2(n+1) + (n+1)^2 = (n+2)^2.$$

##### 1.3 Transitions between states

We denote by $\Omega_{p\to q}$ the Poisson transition rate from state $p$ to state $q$. Transitions are generated by two physical mechanisms.

First, a mobile cohesin arm may encounter an obstacle and become blocked. Such *encounter transitions* convert a free state into the corresponding blocked state. For a static obstacle class $i \neq c$, characterized by mean spacing $d_i$, the encounter rate of a cohesin arm moving with speed $v/k$ is

$$\Omega^{\mathrm{enc}}_{\cdot\to i} = \frac{v}{kd_i}, \tag{1}$$

where $k = 1$ for one-sided extrusion and $k = 2$ for symmetric two-sided extrusion. For the extruder class $i = c$, encounters occur between a moving arm and other cohesins already present on chromatin. In mean field, this gives the effective encounter rate

$$\Omega^{\mathrm{enc}}_{\cdot\to c} = \frac{v}{kd}\left(1 + \frac{kJ}{2}\right), \tag{2}$$

where $d$ is the mean spacing between cohesins and $J \in [0, 1]$ is the steady-state fraction of mobile cohesins unblocked at a given arm. The factor $1 + kJ/2$ accounts for relative motion between the tagged arm and surrounding extruders. In the stationary state, $J$ is determined self-consistently (see Sec. 2).

Second, a blocked arm may be released when the blocking obstacle dissociates. Such *release transitions* occur at rate

$$\Omega^{\mathrm{rel}}_{i\to\cdot} = \frac{1}{\tau_i}, \tag{3}$$

where $\tau_i$ is the lifetime of obstacle class $i$. Thus, whenever an obstacle of class $i$ detaches from a blocked arm, the system returns to the corresponding state in which that arm is mobile.

More explicitly, in one-sided extrusion, encounter transitions are of the form

$$A_0 \to A_i,$$

while obstacle unbinding generates the reverse process

$$A_i \to A_0.$$

For two-sided extrusion, transitions depend on which arm is affected. In the fully resolved notation, encounter transitions are

$$A_{00} \to A_{i0}, \qquad A_{00} \to A_{0i},$$

when the left or right arm becomes blocked, respectively, and

$$A_{i0} \to A_{ij}, \qquad A_{0i} \to A_{ji},$$

when the second arm is subsequently blocked. Conversely, release transitions are

$$A_{i0} \to A_{00}, \qquad A_{0i} \to A_{00},$$

for partial unblocking, and

$$A_{ij} \to A_{i0}, \qquad A_{ij} \to A_{0j},$$

when one of the two blocking obstacles dissociates. Each blocked index therefore contributes its own release channel with rate $1/\tau_i$.

#### 1.4 Cohesin turnover and balance equations

To account for cohesin dissociation and loading, in addition to the chromatin-bound states $A_q$, we introduce a "virtual" pool $A_s$ of soluble cohesins. This pool is fed by dissociation of chromatin-bound cohesins at rate $1/\tau$ and depleted by loading onto chromatin at rate $w$. Loading corresponds to the transition $A_s \to A_{00}$, since cohesin is assumed to bind chromatin in a free state with both arms initially mobile. Because loading starts from zero loop length, the source term in the $A_{00}$ equation is proportional to $\delta(x)$.

For the state populations

$$P_q(t) = \int_0^\infty A_q(x,t)\, dx,$$

the balance equation reads

$$\frac{dP_q}{dt} = \sum_r \left(\Omega_{r\to q} P_r - \Omega_{q\to r} P_q\right), \tag{4}$$

where the sum runs over all states, including the soluble pool. Thus each population $q$ changes due to influx from other states and outflux from state $q$.

#### 1.5 Dimensionless parameters

To compare different extrusion regimes in a unified way, it is convenient to express the model in terms of a small set of dimensionless parameters. The first is the dimensionless density of cohesins,

$$\gamma = \frac{l_p}{d}, \tag{5}$$

and, more generally, the dimensionless density of obstacle class $i$,

$$\gamma_i = \frac{l_p}{d_i}, \tag{6}$$

where $l_p = v\tau$ is the bare processivity length of an unobstructed cohesin. Thus, $\gamma_i$ measures the expected number of encounters with obstacles of class $i$ over one unobstructed processivity length. Small values $\gamma_i \ll 1$ correspond to sparse obstacles, while $\gamma_i \gg 1$ describes a dense barrier landscape.

For the cohesin obstacle class $i = c$, the situation differs from that of static barriers because the effective collision rate depends not only on the mean spacing between cohesins, but also on their relative motion. We therefore define

$$\gamma_c = \gamma \left(1 + \frac{kJ}{2}\right) = \gamma + \gamma_{\text{dyn}}, \tag{7}$$

where $J \in [0, 1]$ is the steady-sate fraction of mobile cohesins unblocked at a given arm. The first term, $\gamma$, is the static contribution associated with the mean spacing between cohesins, while

$$\gamma_{\text{dyn}} = \frac{k\gamma J}{2} \tag{8}$$

is the additional dynamical correction arising from relative motion of extruders. Thus, unlike external barriers, cohesins act as obstacles in a self-consistent way: their effective density depends on the collective state of the extrusion process itself.

The second important parameter is the obstacle persistence,

$$\alpha_i = \frac{\tau}{\tau_i}, \tag{9}$$

defined as the ratio of the cohesin residence time $\tau$ to the lifetime $\tau_i$ of obstacle class $i$. This parameter controls how likely a blocked cohesin is to be released before dissociation. In particular, $\alpha_i \gg 1$ corresponds to short-lived obstacles, which typically unbind rapidly and therefore act as weak barriers, whereas $\alpha_i \ll 1$ corresponds to long-lived obstacles, which can arrest extrusion for a substantial fraction of the cohesin lifetime ($\alpha_c = 1$).

Finally, extrusion symmetry is characterized by the discrete parameter $k$: $k = 1$ for one-sided extrusion and $k = 2$ for symmetric two-sided extrusion. This parameter enters both the encounter rates and the relation between single-arm motion and total loop growth.

### 2. Mean loop length

The key observable characteristic of an extruded loop is its length. In this section, we derive expressions for the mean loop length for unidirectional and bidirectional extrusion and compare the resulting formulas.

#### 2.1 Unidirectional extrusion

**Time-dependent equations** For unidirectional extrusion, the possible bound states consist of one freely extruding state, $A_0$, and $n+1$ blocked states, $A_i$, corresponding to stalling at obstacles of type $i$ (the $n$ barrier types plus $n+1$-th for other cohesins).

To fully specify the dynamics of these states, one must account for both their evolution in time and their transport in loop-length space. For the length-resolved densities $A_q(x,t)$, loop growth enters through convection with instantaneous speed $v$ in the mobile state 0, and with zero speed in the blocked states $i$.

In dimensionless variables $\tilde{x} = x/l_p$ and $\tilde{t} = t/\tau$, the PDE system reads

$$\frac{\partial A_0}{\partial \tilde{t}} + \frac{\partial A_0}{\partial \tilde{x}} = \tilde{w}\, A_s\, \delta(\tilde{x}) - A_0\left(1 + \sum_{i=1}^{n+1} \gamma_i\right) + \sum_{i=1}^{n+1} \alpha_i A_i, \tag{10}$$

$$\frac{\partial A_i}{\partial \tilde{t}} = \gamma_i A_0 - (1+\alpha_i)\, A_i, \tag{11}$$

$$\frac{dA_s}{d\tilde{t}} = \int_0^\infty \left[A_0 + \sum_{i=1}^{n+1} A_i\right] d\tilde{x} \;-\; \tilde{w}\, A_s. \tag{12}$$

The first equation describes the evolution of freely extruding cohesins. The convective term accounts for loop growth, the source term $\tilde{w} A_s \delta(\tilde{x})$ describes loading from the soluble pool into the zero-length state, and the remaining terms represent loss due to dissociation ($-A_0$) or blocking by the obstacle of type $i$ ($-\gamma_i A_0$) and gain due to release from blocked states $i$ ($\alpha_i A_i$). The second line in fact stands for $n+1$ separate equations, one for each obstacle type $i$, including cohesins. Each such equation describes cohesins stalled at an obstacle of type $i$: the corresponding blocked state is populated by transitions from the mobile state (the term $\gamma_i A_0$) and depleted by dissociation ($-A_i$) and obstacle release ($-\alpha_i A_i$). Finally, the equation for $A_s$ expresses balance in the soluble pool, which is fed by dissociation of chromatin-bound cohesins ($\int_0^\infty [A_0 + \sum_i A_i]\, d\tilde{x}$) and depleted by loading onto chromatin ($-\tilde{w}\, A_s$).

**Stationary state** To obtain loop–length statistics we consider the *stationary* regime,

$$\frac{\partial A_0}{\partial \tilde{t}} = \frac{\partial A_i}{\partial \tilde{t}} = \frac{\partial A_s}{\partial \tilde{t}} = 0,$$

and define the stationary occupancies

$$N_b \;=\; \int_0^\infty \left[A_0 + \sum_{i=1}^{n+1} A_i\right] d\tilde{x}, \tag{13}$$

$$N_s \;=\; A_s, \tag{14}$$

where $N_b$ and $N_s$ are the mean numbers of chromatin–bound and soluble cohesins, respectively. Stationarity of $A_s$ then gives

$$0 \;=\; N_b - \tilde{w}\, N_s \quad\Rightarrow\quad \tilde{w} \;=\; \frac{N_b}{N_s}\,. \tag{15}$$

Finally the system of ODEs for stationary state reads

$$\frac{dA_0}{d\tilde{x}} = N_b\, \delta(\tilde{x}) \;-\; A_0\left(1 + \sum_{i=1}^{n+1} \gamma_i\right) \;+\; \sum_i \alpha_i\, A_i, \tag{16}$$

$$0 = \gamma_i\, A_0 \;-\; (1 + \alpha_i)\, A_i, \tag{17}$$

**Solution and mean length** Equation (17) is purely algebraic and yields the blocked components in terms of $A_0$:

$$A_i(\tilde{x}) \;=\; \frac{\gamma_i}{1 + \alpha_i}\, A_0(\tilde{x}). \tag{18}$$

Substituting (18) into (16) gives a single first–order ODE for the free state:

$$\frac{dA_0}{d\tilde{x}} = N_b\, \delta(\tilde{x}) - A_0\left(1 + \sum_{i=1}^{n+1} \frac{\gamma_i}{1 + \alpha_i}\right). \tag{19}$$

This has the explicit solution

$$A_0(\tilde{x}) = N_b\, \exp\left[-\left(1 + \sum_{i=1}^{n+1} \tfrac{\gamma_i}{1+\alpha_i}\right) \tilde{x}\right], \tag{20}$$

$$A_i(\tilde{x}) = \frac{\gamma_i}{1 + \alpha_i}\, A_0(\tilde{x}), \tag{21}$$

which is normalized by construction via the jump condition implied by the $\delta$–source. Equations (20)–(21) make transparent the role of obstacle persistence: short–lived barriers ($\alpha_i \to \infty$) contribute negligibly to the blocking, whereas long–lived barriers ($\alpha_i \ll 1$) contribute in proportion to their encounter density $\gamma_i$.

The sum $\sum_i \gamma_i/(1+\alpha_i)$ aggregates all possible classes (including long–lived non–cohesin obstacles and cohesins). However for the cohesin class $c$ corresponding factor $\gamma_c$ contains dynamic correction $J$, which should be expressed self-consistently.

In steady state, the fraction $J \in [0, 1]$ of time that an extrusion arm is mobile can be defined directly from the stationary state densities within a mean–field (population–averaged) framework.

In the one–sided case only one arm exists and it is mobile if the cohesin is in the free state $A_0$. Therefore the mobile–arm fraction equals the fraction of free states $A_0$:

$$J_{k=1} \;=\; \frac{1}{N_b} \int_0^{\infty} A_0(\tilde{x})\, d\tilde{x}. \tag{22}$$

Here it is convenient to use the notation for the sum in expression (20)

$$\gamma_{\text{eff}} = \sum_{i=1}^{n+1} \frac{\gamma_i}{1+\alpha_i} \tag{23}$$

Writing the total cohesin density as $\gamma$ and the dynamic correction as $\gamma_{\text{dyn}}$, the effective density $\gamma_{\text{eff}}$ becomes

$$\gamma_{\text{eff}} \;=\; \frac{1}{2}\Big(\gamma + \gamma_{\text{dyn}}\Big) \;+\; \gamma_b, \qquad \gamma_b^{\text{eff}} \equiv \sum_{i\neq c}^{n} \frac{\gamma_i}{1+\alpha_i}, \tag{24}$$

and, for unidirectional extrusion, $\gamma_{\text{dyn}} = \frac{\gamma J}{2}$ (by definition from (7) with $k=1$). Using the mean–field expression $J = \Big[\int_0^\infty A_0(\tilde{x})\,d\tilde{x}\Big]\Big/N_b$ together with (20), one obtains a closed equation for $J$,

$$J \;=\; \frac{1}{1+\frac{1}{2}\gamma\big(1+\frac{J}{2}\big)+\gamma_b^{\text{eff}}}, \tag{25}$$

which yields a quadratic equation on $J$ with the following positive solution

$$J \;=\; \frac{\sqrt{\Big(\gamma + 2(\gamma_b^{\text{eff}}+1)\Big)^2 + 4\gamma} - \Big(\gamma + 2(\gamma_b^{\text{eff}}+1)\Big)}{\gamma}\,. \tag{26}$$

Equations (24)–(26) determine $\gamma_{\text{dyn}}$ self–consistently and thereby close the one–sided theory.

The total state–summed loop–length PDF is exponential with the renormalized rate $1+\gamma_{\text{eff}}$:

$$N(\tilde{x}) \;=\; N_b\,(1+\gamma_{\text{eff}})\,\exp\!\Big[-(1+\gamma_{\text{eff}})\,\tilde{x}\Big], \tag{27}$$

so the dimensionless mean loop length is

$$\tilde{\lambda} = \frac{\lambda}{l_p} \;=\; \int_0^\infty \tilde{x}\,N(\tilde{x})\,d\tilde{x} \;=\; \frac{1}{1+\gamma_{\text{eff}}}. \tag{28}$$

This result has a direct kinetic interpretation: the effective hazard per unit length is the sum of the intrinsic dissociation rate (set to unity in our units) and the encounter rate with the effective population of blockers $\gamma_{\text{eff}}$. Consequently, the mean loop size equals the obstacle–free processivity reduced by the factor $1+\gamma_{\text{eff}}$; short–lived or sparse obstacles contribute little, while long–lived or dense obstacles (including the dynamic cohesin term) suppress the attainable loop size more strongly.

#### 2.2 Bidirectional extrusion

For unidirectional extrusion, we obtained a simple and universal expression for the mean loop length. The symmetric bidirectional case can be reduced to the same problem by considering the two cohesin legs separately. Indeed, the total loop length is the sum of the lengths extruded by the left and right arms, and by linearity of expectation its mean is the sum of the corresponding one-arm means. Since the two arms are statistically identical in the symmetric regime, this gives

$$\tilde{\lambda} \;=\; 2\,\mathbb{E}[\tilde{\ell}_{\text{one arm}}].$$

To distinguish the motion of the two cohesin legs, we use the two-coordinate formulation, with arm-resolved variables $(\tilde{\ell}, \tilde{r})$ for the lengths extruded by the left and right legs, respectively.

To fully specify the dynamics of the state-resolved densities, one must account for both their evolution in time and their transport in the two-dimensional loop-length space. In this representation, loop growth enters through convection along the corresponding coordinate directions: in the fully mobile state 00, both legs move, so the density is convected in both $\tilde{\ell}$ and $\tilde{r}$ directions with speed $v/2$ per leg (equivalently, with total loop-growth speed $v$); in the partially blocked states $0i$ and $i0$, only one leg remains mobile, so convection occurs only along the coordinate of the free leg, with speed $v/2$; and in the fully blocked states $ij$, both legs are stalled and no convective term is present.

The corresponding master-equations are written as follows

$$\frac{\partial A_{00}}{\partial \tilde{t}} + \frac{1}{2}\frac{\partial A_{00}}{\partial \tilde{\ell}} + \frac{1}{2}\frac{\partial A_{00}}{\partial \tilde{r}} = \tilde{w}\,A_s\,\delta(\tilde{l}, \tilde{r}) - \,A_{00}\left(1 + \sum_{i=1}^{n+1} \gamma_i\right) + \sum_{i=1}^{n+1} \alpha_i \Big(A_{i0} + A_{0i}\Big), \tag{29}$$

$$\frac{\partial A_{0i}}{\partial \tilde{t}} + \frac{1}{2}\frac{\partial A_{0i}}{\partial \tilde{\ell}} = \frac{\gamma_i}{2}\,A_{00} - \left(1 + \alpha_i + \frac{1}{2}\sum_{j=1}^{n+1} \gamma_j\right) A_{0i} + \sum_{j=1}^{n+1} \alpha_j A_{ji}, \tag{30}$$

$$\frac{\partial A_{i0}}{\partial \tilde{t}} + \frac{1}{2}\frac{\partial A_{i0}}{\partial \tilde{r}} = \frac{\gamma_i}{2}\,A_{00} - \left(1 + \alpha_i + \frac{1}{2}\sum_{j=1}^{n+1} \gamma_j\right) A_{i0} + \sum_{j=1}^{n+1} \alpha_j A_{ij}, \tag{31}$$

$$\frac{\partial A_{ij}}{\partial \tilde{t}} = \frac{\gamma_j}{2}\,A_{i0} + \frac{\gamma_i}{2}\,A_{0j} - (1 + \alpha_i + \alpha_j)\,A_{ij}. \tag{32}$$

The qualitative interpretation of terms is the same as for the corresponding ones for unidirectional extrusion.

**Stationary state** The stationary-state limit is obtained in complete analogy with the unidirectional case. The corresponding stationary equations read

$$\frac{1}{2}\frac{\partial A_{00}}{\partial \tilde{\ell}} + \frac{1}{2}\frac{\partial A_{00}}{\partial \tilde{r}} = N_b\, \delta(\tilde{l}, \tilde{r}) - A_{00}\left(1 + \sum_{i=1}^{n+1} \gamma_i\right) + \sum_{i=1}^{n+1} \alpha_i \Big(A_{i0} + A_{0i}\Big), \tag{33}$$

$$\frac{1}{2}\frac{\partial A_{0i}}{\partial \tilde{\ell}} = \frac{\gamma_i}{2} A_{00} - \left(1 + \alpha_i + \frac{1}{2}\sum_{j=1}^{n+1} \gamma_j\right) A_{0i} + \sum_{j=1}^{n+1} \alpha_j A_{ji}, \tag{34}$$

$$\frac{1}{2}\frac{\partial A_{i0}}{\partial \tilde{r}} = \frac{\gamma_i}{2} A_{00} - \left(1 + \alpha_i + \frac{1}{2}\sum_{j=1}^{n+1} \gamma_j\right) A_{i0} + \sum_{j=1}^{n+1} \alpha_j A_{ij}, \tag{35}$$

$$0 = \frac{\gamma_j}{2} A_{i0} + \frac{\gamma_i}{2} A_{0j} - (1 + \alpha_i + \alpha_j)\, A_{ij}. \tag{36}$$

**Solution and mean length** To reduce this system to an effective one-dimensional description, we integrate over the coordinate of the *other* arm, say $\tilde{r} \in [0, \infty)$. This projects the two-dimensional dynamics onto the left-arm coordinate $\tilde{\ell}$ and yields equations for states classified only by the status of the left arm.

We therefore introduce

$$A_{0.}(\tilde{\ell}) = \int_0^\infty \left[A_{00}(\tilde{\ell}, \tilde{r}) + \sum_{i=1}^{n+1} A_{0i}(\tilde{\ell}, \tilde{r})\right] d\tilde{r}, \tag{37}$$

$$A_{i.}(\tilde{\ell}) = \int_0^\infty \left[A_{i0}(\tilde{\ell}, \tilde{r}) + \sum_{j=1}^{n+1} A_{ij}(\tilde{\ell}, \tilde{r})\right] d\tilde{r}, \tag{38}$$

so that $A_{0.}$ collects all configurations with a free left arm, while $A_{i.}$ collects all configurations with the left arm blocked by an obstacle of class $i$, irrespective of the state of the right arm.

*Equation for $A_{0.}$.* To derive the balance equation for $A_{0.}$, we integrate Eq. (33) over $\tilde{r}$ and add the sum over $i$ of Eq. (34), also integrated over $\tilde{r}$. The left-hand side becomes

$$\frac{1}{2}\frac{d}{d\tilde{\ell}} \int_0^\infty \left(A_{00} + \sum_i A_{0i}\right) d\tilde{r} + \frac{1}{2}\int_0^\infty \frac{\partial A_{00}}{\partial \tilde{r}}\, d\tilde{r}.$$

Assuming regular decay as $\tilde{r} \to \infty$, the second term reduces to a delta-function at $\tilde{r} = 0$.

After collecting all terms and using the definitions (37)–(38) and multiplying by 2, one finds

$$\frac{dA_{0.}}{d\tilde{\ell}} = N_b\, \delta(\tilde{\ell}) - 2\left(1 + \sum_{i=1}^{n+1} \frac{\gamma_i}{2}\right) A_{0.} + 2\sum_{i=1}^{n+1} \alpha_i A_{i.}. \tag{39}$$

To obtain the equation for a fixed blocked class $i$, we integrate Eq. (35) over $\tilde{r}$ and add the sum over $j$ of Eq. (36), again integrated over $\tilde{r}$. This combines the state $A_{i0}$ with all states $A_{ij}$, i.e. precisely into $A_{i.}$. The derivative term in Eq. (35) is with respect to $\tilde{r}$, so after integration over $\tilde{r}$ it produces only boundary terms, which vanish for given boundary conditions ($A_{i0}(0,0) = 0$, $A_{ij}(0,0) = 0$). Hence the resulting balance is purely algebraic.

Therefore one obtains

$$0 = \frac{\gamma_i}{2} A_{0.} - (1+\alpha_i) A_{i.},$$

or equivalently

$$A_{i.} = \frac{\gamma_i}{2(1+\alpha_i)} A_{0.}. \tag{40}$$

Eliminating $A_{i.}$ from (39)–(40) we obtain a single first–order ODE for $A_{0.}$, whose solution for $\tilde{\ell} > 0$ is purely exponential,

$$A_{0.}(\tilde{\ell}) \;=\; N_b \exp\left[-2\left(1 + \sum_{i=1}^{n+1} \frac{\gamma_i}{2(1+\alpha_i)}\right)\tilde{\ell}\right]. \tag{41}$$

Thus, the *one–arm* length is exponentially distributed. In particular,

$$\mathbb{E}[\tilde{\ell}_{\text{one arm}}] \;=\; \frac{1}{2\left(1 + \sum_{i=1}^{n+1} \frac{\gamma_i}{2(1+\alpha_i)}\right)} = \frac{1}{2\left(1 + \frac{1}{2}\gamma_{\text{eff}}\right)},$$

where we use the same notation as for the case of unidirectional extrusion considering the effective density $\gamma_{\text{eff}}$ according to (24).

As in the case of unidirectional extrusion, the stationary equations contain a dynamical correction $\gamma_{\text{dyn}}$ determined through $J$, which must be determined self-consistently from the following considerations.

For two-sided extrusion, each cohesin has two arms, left and right. In the symmetric case these are statistically equivalent, so it is sufficient to consider one reference arm, say the left one. This arm contributes to the mobile fraction whenever it is free to move, which occurs either in the fully mobile state $A_{00}$, or in the partially blocked states $A_{0i}$, where the opposite arm is blocked while the reference arm remains mobile. The corresponding mobile-arm fraction is therefore

$$J_{k=2} \;=\; \frac{1}{N_b}\int_0^\infty \left[A_{00}(\tilde{x}) + \sum_{i=1}^{n+1} A_{0i}(\tilde{x})\right] d\tilde{x} = \frac{1}{N_b}\int_0^\infty A_{0.} d\tilde{\ell}. \tag{42}$$

By left–right symmetry, the same expression is obtained if one instead tracks the right arm, i.e. by replacing $\{A_{00}, A_{0i}\}$ with $\{A_{00}, A_{i0}\}$.

This definition closes the mean-field problem exactly as in the one-sided case. In particular, for symmetric two-sided extrusion ($k = 2$), the quantity $J$ satisfies the same fixed-point relation (7), now with the appropriate two-sided state space. Solving this relation yields

$$\gamma_{\text{dyn}} \;=\; \frac{\sqrt{8\gamma + \left[\gamma + 4(1+\gamma_b^{\text{eff}})\right]^2} \;-\; \left[\gamma + 4(1+\gamma_b^{\text{eff}})\right]}{2}. \tag{43}$$

The mean *total* loop length follows as twice the one–arm mean:

$$\tilde{\lambda} \;=\; 2\,\mathbb{E}[\tilde{\ell}_{\text{one arm}}] \;=\; \frac{1}{1+\gamma_{\text{eff}}/2}. \tag{44}$$

where

$$\gamma_{\text{eff}} \;=\; \frac{1}{2}\Big(\gamma+\gamma_{\text{dyn}}\Big) \;+\; \gamma_b^{\text{eff}}, \qquad \gamma_b^{\text{eff}} \equiv \sum_{i\neq c}^{n} \frac{\gamma_i}{1+\alpha_i}, \tag{45}$$

This result has a transparent kinetic interpretation: relative to the obstacle–free case ($\tilde{\lambda} = 1$), two–sided extrusion suffers an effective *per–arm* termination hazard equal to $\gamma_{\text{eff}}/2$, because each arm advances at half the total speed and experiences both static barriers and moving–extruder encounters.

#### 2.3 Discussion of results

##### 2.3.1 Universal formula for mean loop size

The one– Eq. (28) and two–sided Eq. (44) formulas can be summarized as

$$\tilde{\lambda} \;=\; \frac{1}{1+\gamma_{\text{eff}}/k}, \qquad k = \begin{cases} 1, & \text{one–sided,} \\ 2, & \text{two–sided,} \end{cases} \tag{46}$$

with $\gamma_{\text{eff}}$ given by the corresponding ($k$-dependent) mean–field expression. In the absence of obstacles the characteristic loop length equals processivity $\tilde{\lambda} = 1$. Any class of barriers (CTCF, transcriptional machinery, other cohesins) reduces $\tilde{\lambda}$ by introducing pauses and premature terminations of extrusion.

Equation (46) shows that in the presence of barriers, bidirectional ($k = 2$) extrusion yields longer loops than one-sided ($k = 1$). Note, however, that the effective barrier density $\gamma_{\text{eff}}$ itself depends on the extrusion mode $k$ (via $\gamma_{\text{dyn}}$ and the fraction of mobile arms $J$), but this dependence is weak and is largely a corrective effect.

Effective density of barriers combines contributions from three factors:

$$\gamma_{\text{eff}} = \frac{\gamma+\gamma_{\text{dyn}}}{2} + \gamma_{\text{b}}, \tag{47}$$

where $\gamma$ is the contribution from interaction with static cohesins, $\gamma_{\text{b}}$ is the contribution from other proteins ("b" stands for contribution from all barriers), $\gamma^{(k)}_{\text{dyn}}$ corresponds to the interaction with moving extruders. $\gamma$ and $\gamma_b$ are the same for bidirectional and unidirectional extrusion. The effective crowding parameter $\gamma_{\text{eff}}$ can be interpreted as the expected number of obstacles an extruder would encounter during its lifetime in the absence of persistent blocking

Below we will consider the factors contributing to $\gamma_{\text{eff}}$ and their relative importance.

1. Cohesin density:

$$\gamma = \frac{l_p}{d}, \tag{48}$$

represents the intrinsic crowding due to extruders themselves.

2. Static barriers

$$\gamma_{\mathrm{b}} = \sum_{i=1}^{n} \frac{\gamma_i}{1+\alpha_i}. \tag{49}$$

Here, $\gamma_i = l_p/d_i$ is the contribution of obstacles of type $i$ with spacing $d_i$, and $\alpha_i = \tau/\tau_i$ compares cohesin residence time $\tau$ to the lifetime of obstacle $i$ ($\tau_i$). $\gamma_{\mathrm{b}}$ accounts for barriers such as CTCF-bound sites, nucleosomes, stalled transcriptional complexes, or other DNA-bound proteins.

3. Dynamic collisions. Encounters with other moving extruders are captured by

$$\gamma_{\mathrm{dyn}}^{(k)} = \frac{\sqrt{4k\gamma + \left[2(1+2\gamma_{\mathrm{b}}^{\mathrm{eff}})+\gamma\right]^2} - \left[2(1+2\gamma_{\mathrm{b}}^{\mathrm{eff}})+\gamma\right]}{2},$$

where $k = 1$ (unidirectional) or $k = 2$ (bidirectional). Unlike static terms, this contribution is nonlinear because the fraction of mobile extruder arms depends self-consistently on $\tilde{\lambda}$, extruder density, and blocking by both static and dynamic obstacles (in the bidirectional case, an unblocked opposing arm never appears static). The expression has simple limits: in sparse conditions ($\gamma, \gamma_{\mathrm{b}} \to 0$), $\gamma_{\mathrm{dyn}}^{(k)} \approx (k/2)\gamma$ (i.e., $\sim 1.5\times$ more terminations than purely static for $k = 1$, $\sim 2\times$ for $k = 2$); in dense, extruder-dominated regimes ($\gamma \gg \gamma_{\mathrm{b}}$) it saturates at $\gamma_{\mathrm{dyn}}^{(k)} \to k$. Therefore, when steric terms are appreciable ($\gamma + 2\gamma_{\mathrm{b}} \gg k$), the dynamic contribution is relatively small; in practice (and in the systems analyzed below), $\gamma_{\mathrm{eff}}$ can be well approximated by summing static contributions from all relevant classes, with $\gamma_{\mathrm{dyn}}^{(k)}$ providing only a minor correction.

Equation (46) admits a natural kinetic interpretation: the scale factor $\tilde{\lambda}$ is the ratio of the effective, crowding-limited loop–growth speed to the intrinsic extrusion speed, i.e., $\tilde{\lambda} = \langle v \rangle / v$ (unity in the obstacle–free limit). Consistent with this view, single-molecule assays report $v \approx 1$ kb/s for human cohesin [Davidson et al., 2019], whereas genome-wide Hi-C recovery in HeLa cells yields a much smaller population-averaged $\langle v \rangle \approx 0.375$ kb/s [Rao et al., 2017]. Using Eq. (46) together with independently compiled densities and lifetimes of obstacles and cohesins (Table S1) to estimate $\gamma_{\mathrm{eff}}$, we obtain $\tilde{\lambda} \approx 0.37 \pm 0.07$ across the HeLa range. Thus, the mean loop length is typically only $\sim$ 30–50% of the intrinsic processivity $l_p$, reflecting substantial suppression by long-lived static barriers with a modest additional reduction from cohesin–cohesin encounters, in quantitative agreement with the in vivo vs. in vitro velocity discrepancy.

##### 2.3.2 WAPL depletion experiments

WAPL depletion provides an orthogonal test of the theory. Because WAPL promotes cohesin unloading from chromatin (Fig. S1a), its depletion is expected to increase cohesin residence time and to shift cohesin from the soluble pool onto DNA. In the model, these two effects act in different directions: they increase the bare processivity $l_p$ while reducing the mean spacing $d$ between chromatin-bound cohesins.

Figure S1b compares the predicted $\lambda(l_p)$ curves with experimental estimates for WT (red) and ΔWAPL (blue). The loop size used in this comparison was extracted from the experimental studies either from the position of the peak in the logarithmic derivative of the Hi-C contact probability or from polymer-model fits to Hi-C data, as summarized in Table S3. To compare with theory, we first constructed the WT curve using the cohesin spacing $d_{\rm WT} \approx 174$ kb, close to the experimental estimate (Table S1). With this value, the bidirectional prediction ($k = 2$, solid line) reproduces the observed WT loop size well. Repeating the same analysis for the ΔWAPL condition, we find that the experimental loop size is again captured by bidirectional extrusion, provided the effective cohesin spacing is reduced to $d_{\rm WAPL} \approx 104$ kb. To obtain the effective obstacle densities for the WT and ΔWAPL conditions, we recalculated the dimensionless parameters using the corresponding mean cohesin residence times (Table S3). Given an approximately 14-fold increase in the residence time (and, thus, $l_p$), this yields $\gamma_{\rm WT} \approx 4.0$ and $\gamma_{\rm WAPL} \approx 95.1$.

The genomic spacing and lifetime of the static barriers were assumed to remain unchanged upon WAPL depletion. Accordingly, the increase in cohesin residence time changes both the bare barrier density, $\gamma_b = l_p/d_b$, and the relative barrier turnover rate, $\alpha = \frac{\tau}{\tau_b}$. Starting from $\gamma_b = 4.0$ and $\alpha = 1.2$ in WT (Table S2), we obtain $\gamma_b^{\rm eff} = \frac{\gamma_b}{1+\alpha} \approx 1.8$ for WT and $\gamma_b^{\rm eff} \approx 3.15$ for ΔWAPL.

Finally, eq. (47) gives

$$\gamma_{\rm eff}^{\rm WT} \approx 4.1, \qquad \gamma_{\rm eff}^{\Delta\rm WAPL} \approx 51.6.$$

The ΔWAPL perturbation also allows one to estimate the fraction of soluble cohesin in WT. In the simplest interpretation, WAPL depletion drives most of the total cohesin pool onto chromatin, so that the mean spacing changes according to

$$d_{\rm WAPL} = \frac{N_b}{N_b + N_f}\, d_{\rm WT},$$

where $N_b$ and $N_f$ are the chromatin-bound and soluble cohesin numbers in WT. Therefore,

$$\frac{d_{\rm WAPL}}{d_{\rm WT}} = \frac{N_b}{N_b + N_f}$$

directly gives the fraction of cohesins already bound in WT. Using the fitted values, we obtain

$$\frac{N_b}{N_b + N_f} \approx \frac{104}{174} \approx 0.6.$$

This implies that about 60% of cohesins are chromatin-bound in WT, while the remaining $\sim 40\%$ belong to the soluble pool, consistent with the available experimental estimates (Table S1).

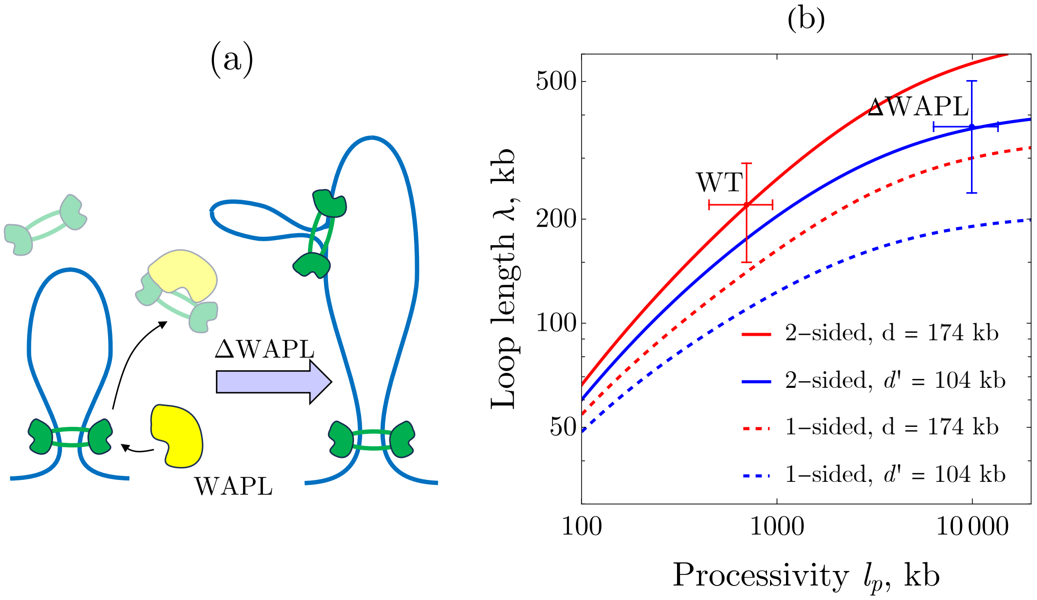

Figure S1: Effect of WAPL unloading on cohesin abundance and loop size. (a) Schematic of WAPL-mediated cohesin unloading. WAPL depletion (ΔWAPL) prolongs cohesin residence on chromatin and shifts cohesin from the soluble pool to DNA, increasing chromatin-bound abundance and reducing the mean spacing between complexes. (b) Predicted mean loop size $\lambda$ versus intrinsic processivity $l_p$. Red: wild type (WT). Blue: ΔWAPL. Solid lines: bidirectional extrusion; dashed lines: one-sided extrusion. Points (mean ± range) show experimental estimates for WT and ΔWAPL; sources are listed in Table S3. $\gamma_{\text{eff}} = 4.1$ for WT and $\gamma_{\text{eff}} = 51.6$ for ΔWAPL

### 3. Loop-length distribution

We first present a formal kinetic-equation approach, which yields the exact state-resolved stationary distributions of loop lengths. We then reformulate the same dynamics in terms of transport on a directed graph, which makes the origin of the resulting distribution classes transparent. In particular, the graph mapping shows directly why one-sided extrusion produces a single exponential distribution, whereas bidirectional extrusion generically gives a multiexponential, and therefore typically peaked, loop-length distribution.

#### 3.1 Kinetic equations approach

We now formulate the kinetic equations for the state-resolved probability densities. These equations describe the joint evolution of loop length and internal cohesin state under extrusion, blocking, obstacle release, loading, and dissociation.

##### 3.1.1 Unidirectional extrusion

The full kinetic equations for unidirectional extrusion were derived in Sec. 2.1, where they were used to derive the mean loop length. We briefly recall them here for completeness before proceeding to the bidirectional case

$$\frac{dA_0}{d\tilde{x}} = N_b\,\delta(\tilde{x}) - A_0\Bigg(1 + \sum_{i=1}^{n+1} \frac{\gamma_i}{1+\alpha_i}\Bigg), \tag{50}$$

$$A_i(\tilde{x}) = \frac{\gamma_i}{1+\alpha_i}\, A_0(\tilde{x}), \tag{51}$$

The total loop-length distribution is purely exponential,

$$N(\tilde{x}) = N_b(1+\gamma_{\text{eff}})\exp\Big[-(1+\gamma_{\text{eff}})\tilde{x}\Big], \tag{52}$$

with the mean $\tilde{\lambda} = (1+\gamma_{\text{eff}})^{-1}$, where $\gamma_{\text{eff}}$ collects the contributions of all obstacle classes, including the self-consistent dynamical correction due to other cohesins (Eq. (24)). Thus, in the one-sided case both the full distribution and the mean loop length are controlled by the same single parameter $\gamma_{\text{eff}}$.

##### 3.1.2 Bidirectional extrusion

**Stationary equations** We now turn to the *symmetric two–sided* extrusion regime, where both arms of a cohesin advance at equal instantaneous speeds and may be blocked independently on the left or on the right, with $n+1$ types of obstacles, including cohesin. In the stationary, time–integrated formulation the nonzero bound–state densities are: (i) the fully free state $A_{00}(\tilde{x})$; (ii) one–sidedly blocked states $A_i(\tilde{x})$, where the moving arm is arrested by an obstacle of class $i$; and (iii) the fully blocked states $A_{ij}(\tilde{x})$ with obstacles of classes $j$ and $i$ on the two sides. Here $\gamma_i = l_p/d_i$ and $\alpha_i = \tau/\tau_i$ are, respectively, the (dimensionless) encounter density and persistence of obstacle class $i$, as in the unidirectional case.

To derive the stationary loop-length distribution in the bidirectional case, we proceed as in Sec. 2, starting from the time-integrated master equations and then passing to the stationary limit. As before, the full set of state-resolved equations includes a source term at $\tilde{x} = 0$, which represents loading of cohesin into the zero-length state. However, for the purpose of determining the shape of the stationary distribution away from the loading point, it is sufficient to consider the region $\tilde{x} > 0$, where the $\delta$-function source is absent. In this region the problem reduces to a homogeneous system of linear balance equations:

$$\frac{dA_{00}}{d\tilde{x}} = -A_{00}\left(1 + \sum_{i=1}^{n+1} \gamma_i\right) + \sum_{i=1}^{n+1} \alpha_i\, A_i, \tag{53}$$

$$\frac{dA_i}{d\tilde{x}} = 2\gamma_i A_{00} - 2\Big(1+\alpha_i\Big)\left(1 + \sum_{j=1}^{n+1} \frac{\gamma_j}{2(1+\alpha_i+\alpha_j)}\right) A_i + 2\sum_{j=1}^{n+1} \frac{\alpha_j\gamma_i}{2(1+\alpha_i+\alpha_j)}\, A_j. \tag{54}$$

$$A_{ij}(\tilde{x}) = \frac{\gamma_j A_i(\tilde{x}) + \gamma_i A_j(\tilde{x})}{4\Big(1+\alpha_i+\alpha_j\Big)}. \tag{55}$$

The last line expresses the fully blocked, zero–velocity states $A_{ij}$ via algebraic (instantaneous) balance between the inflow from one–sidedly blocked states and outflow by obstacle unbinding.

**The distribution of states** For $\tilde{x} > 0$ (i.e. away from the $\delta$–source), the system (53)–(54) can be written compactly as

$$\frac{d}{d\tilde{x}}\begin{pmatrix} A_{00} \\ A_1 \\ \vdots \\ A_{n+1} \end{pmatrix} = \mathbb{M}\begin{pmatrix} A_{00} \\ A_1 \\ \vdots \\ A_{n+1} \end{pmatrix}, \qquad \frac{d}{d\tilde{x}}\mathbf{A}(\tilde{x}) = \mathbb{M}\,\mathbf{A}(\tilde{x}). \tag{56}$$

Because $\mathbb{M}$ is a constant matrix (for fixed parameters $\{\gamma_i, \alpha_i\}$), the solution for $\tilde{x} > 0$ is obtained by spectral decomposition. In the generic case, when the eigenvalues of $\mathbb{M}$ are pairwise distinct, the stationary solution is a superposition of simple exponentials,

$$\begin{pmatrix} A_{00} \\ A_1 \\ \vdots \\ A_{n+1} \end{pmatrix} = \sum_{m=1}^{n+2} B_m \mathbf{u_m} e^{-\beta_m \tilde{x}}, \qquad \beta_m \equiv -\mu_m(\mathbb{M}) > 0, \tag{57}$$

where $\{\mu_m\}$ are the eigenvalues of $\mathbb{M}$, $\{\mathbf{u_m}\}$ are the eigenvectors of $\mathbb{M}$, and the coefficients $\{B_m\}$ are fixed by the boundary condition at $\tilde{x} = 0$. The fully blocked components $A_{ij}$ then follow from Eq. (55).

Equation (57) is valid provided that no eigenvalue degeneracy occurs, i.e. for a non-defective $\mathbf{M}$. If two or more eigenvalues coincide, the corresponding solution is no longer a sum of pure exponentials only: polynomial prefactors appear, yielding terms of the form $\tilde{x}^p e^{-\beta\tilde{x}}$. In particular, the simplest degeneracy gives rise to $\tilde{x}e^{-\beta\tilde{x}}$, i.e. a gamma-distribution-type contribution. In the minimal examples from the next paragraphs we will show that for finite values of effective densities there is no degeneracy, however it can be present as limiting case. The loop-size PDF is then obtained as

$$N(\tilde{x}) = A_{00}(\tilde{x}) + \sum_{i=1}^{n+1} A_i(\tilde{x}) + \sum_{i,j=1}^{n+1} A_{ij}(\tilde{x}). \tag{58}$$

**Example: one obstacle type + transparent cohesin** We next consider a minimal nontrivial example of symmetric bidirectional extrusion in the presence of a single barrier type (class $b$) and transparent cohesins. We denote by $A_{00}$ the state in which both arms are mobile, by $A_b$ the total population of states in which one arm is blocked by obstacle and the other arm remains mobile, and by $A_{bb}$ the state in which both arms are blocked by obstacles. Here $\gamma_b$ is the dimensionless density of static barriers, and $\alpha = \tau/\tau_b$ the ratio of the cohesin residence time to the barrier lifetime.

This example is useful because it already captures an important qualitative feature of the full model: while the *mean* loop length can often be expressed through a single effective crowding parameter, the *full* stationary distribution generally depends separately on $\alpha$ and $\gamma_b$. In particular, obstacle persistence cannot in general be absorbed into a single renormalized density.

For a single obstacle class the master equations (53-55) read

$$\frac{\partial A_{00}}{\partial \tilde{x}} = -A_{00}\Big(1+\gamma_b\Big) + \alpha A_b, \tag{59}$$

$$\frac{\partial A_b}{\partial \tilde{x}} = 2\gamma_b A_{00} - \Big(2+2\alpha+\gamma_b\Big)A_b + 4\alpha A_{bb}. \tag{60}$$

$$A_{bb} = \frac{\gamma_b}{2(1+2\alpha)}\,A_b \tag{61}$$

Substituting the algebraic coupling into this ODE system, we obtain

$$\frac{\partial A_{00}}{\partial \tilde{x}} = -A_{00}\Big(1+\gamma_b\Big) + \alpha A_b, \tag{62}$$

$$\frac{\partial A_b}{\partial \tilde{x}} = 2\gamma_b A_{00} - \Big(2+2\alpha+\frac{\gamma_b}{1+2\alpha}\Big)A_b \tag{63}$$

$$A_{bb} = \frac{\gamma_b}{2(1+2\alpha)}\,A_b \tag{64}$$

The ODE-part of these equations can be written compactly in the matrix form as

$$\frac{d\mathbf{A}}{d\tilde{x}} = \mathbb{M}\,\mathbf{A}, \qquad \mathbf{A}(\tilde{x}) = \begin{pmatrix} A_{00}(\tilde{x}) \\ A_b(\tilde{x}) \end{pmatrix}, \tag{65}$$

with

$$\mathbb{M} = \begin{pmatrix} -(1+\gamma_b) & \alpha \\ 2\gamma_b & -\Big(2+2\alpha+\frac{\gamma_b}{1+2\alpha}\Big) \end{pmatrix}. \tag{66}$$

The solution is determined by two conditions. First,

$$A_b(0) = 0, \tag{67}$$

because at zero loop length the cohesin is loaded in the fully mobile state, with both arms initially unblocked. Second, normalization requires

$$\int_0^\infty \left(A_{00} + A_b + A_{bb}\right)\, d\tilde{x} = N_b. \tag{68}$$

The general solution is a linear combination of the eigenmodes of $\mathbb{M}$. Since the two eigenvalues are distinct, the total stationary loop-length distribution $N(\tilde{x}) = A_{00}(\tilde{x}) + A_b(\tilde{x}) + A_{bb}(\tilde{x})$ is a sum of two exponentials,

$$N(\tilde{x}) = N_b(C_1 e^{-\beta_1 \tilde{x}} + C_2 e^{-\beta_2 \tilde{x}}), \tag{69}$$

where the decay constants are

$$\beta_1 = 2(1+\alpha) + \gamma_b, \qquad \beta_2 = 1 + \frac{\gamma_b}{1+2\alpha}. \tag{70}$$

The coefficients $C_1$ and $C_2$ are fixed by the boundary and normalization conditions. Carrying out this calculation gives

$$C_1 = -\frac{\gamma_b(2\alpha + \gamma_b + 2)}{4\alpha^2 + 2\alpha(\gamma_b + 2) + 1}, \tag{71}$$

$$C_2 = \frac{(2\alpha + \gamma_b + 1)^2}{4\alpha^2 + 2\alpha(\gamma_b + 2) + 1}. \tag{72}$$

The corresponding mean loop length is

$$\lambda = \frac{1}{N_b}\int_0^\infty \tilde{x}\, N(\tilde{x})\, d\tilde{x} = \frac{1}{1 + \frac{\gamma_b}{2(1+\alpha)}}. \tag{73}$$

The same result is obtained from Eq. (44). In other words, at the level of the mean loop length, a barrier with finite lifetime is equivalent to an effective barrier with the renormalized density $\gamma_b/(1+\alpha)$.

At the level of the full distribution, however, the dependence on $\alpha$ and $\gamma_b$ remains irreducible: both the decay constants $\beta_{1,2}$ and their weights depend separately on these two parameters. Thus, unlike the mean loop length, the full stationary distribution cannot in general be collapsed onto a single effective crowding parameter.

A particularly instructive limit is $\gamma_b \gg 1$, where the structure of the solution changes qualitatively. In the next paragraph we show that, for long-lived barriers ($\alpha = 0$), the asymptotic approach of the two decay constants gives a $\Gamma$-distribution.

**Asymptotic emergence of a Γ-distribution** An important question is what happens when the decay constants in the modal solution become degenerate. Formally, exact degeneracy would require $\beta_1 = \beta_2$. Directly solving this condition shows that it would occur only at a negative value of $\gamma_b$ $\left(\gamma_b = -\frac{(1+2\alpha)^2}{2\alpha}\right)$, which is outside the physical parameter range. Thus, for finite positive parameters the eigenvalues remain distinct and the solution is, strictly speaking, always a sum of simple exponentials.

However, a physically meaningful near-degeneracy does arise in the singular limit

$$\alpha = 0, \qquad \gamma_b \gg 1,$$

corresponding to very dense and effectively permanent barriers. In this regime, the two eigenvalues approach one another asymptotically,

$$\beta_1, \beta_2 \to \gamma_b,$$

and the matrix $\mathbb{M}$ is dominated by the leading terms proportional to $\gamma_b$. Retaining only these dominant contributions gives

$$\mathbb{M} = \begin{pmatrix} -\gamma_b & 0 \\ 2\gamma_b & -\gamma_b \end{pmatrix}. \tag{74}$$

This matrix has a repeated eigenvalue

$$\mu_1 = \mu_2 = -\gamma_b,$$

so the corresponding solutions therefore contain a polynomial prefactor,

$$A(\tilde{x}) \sim (C_1 + C_2\tilde{x})e^{-\gamma_b \tilde{x}}.$$

Solving with the appropriate boundary conditions yields

$$A_{00}(\tilde{x}) \sim e^{-\gamma_b \tilde{x}}, \qquad A_b(\tilde{x}) \sim 2\gamma_b \tilde{x}\, e^{-\gamma_b \tilde{x}},$$

and, using the algebraic relation for the fully blocked state,

$$A_{bb}(\tilde{x}) \sim \gamma_b^2 \tilde{x}\, e^{-\gamma_b \tilde{x}}.$$

Summing over all populated states, the total loop-length distribution becomes

$$N(\tilde{x}) \sim e^{-\gamma_b \tilde{x}}\Big[1 + \gamma_b(2+\gamma_b)\tilde{x}\Big] \sim \gamma_b^2\, \tilde{x}\, e^{-\gamma_b \tilde{x}},$$

which is precisely the form of a Γ-distribution with shape parameter 2.

This asymptotic Γ-distribution has a simple physical origin. In the limit of very dense, permanent barriers ($\gamma_b \gg 1$, $\alpha = 0$), the two cohesin arms become effectively independent and extrude separate random genomic segments. The total loop length is then the sum of two independent exponentially distributed arm lengths, yielding a Γ-distribution with shape parameter 2. For any $\alpha > 0$, barriers are releasable, so this complete decoupling is lost even at large $\gamma_b$. The resulting residual coupling appears as a finite arm–arm correlation, which we analyze in Sec. 4. We now specialize to the minimal biologically relevant case: bidirectional extrusion with a single static obstacle class and opaque cohesins, for which $\alpha_c = 1$.

**Example: one obstacle + opaque cohesin** We now consider a minimal biologically relevant example: bidirectional extrusion in the presence of two blocker classes, namely a single static obstacle class ($b$) and other cohesins ($c$), treated as impermeable. This is the simplest setting that already retains the two qualitatively distinct sources of blocking present in vivo: long-lived external barriers (small $\alpha$) and steric encounters with other extruders ($\alpha = 1$). At the same time, it is physically instructive because it allows one to follow explicitly the effect of relatively short-lived obstacles (extruders), which produces nontrivial effects in the stationary loop-length distribution and in the synchronization of the two extrusion legs.

We denote by $A_{00}$ the state in which both arms are mobile, by $A_c$ the total population of states in which one arm is blocked by another cohesin and the other arm remains mobile, and by $A_b$ the analogous population for blocking by the static obstacle class. Similarly, $A_{cc}$ denotes the state in which both arms are blocked by cohesins, $A_{bc}$ the combined population of states in which one arm is blocked by a static obstacle and the other by a cohesin, and $A_{bb}$ the state in which both arms are blocked by static obstacles. Here and below, $\gamma$ denotes the dimensionless density of cohesins, $\gamma_b$ the dimensionless density of static barriers, and $\alpha = \tau/\tau_b$ the ratio of the cohesin residence time to the barrier lifetime.

In the stationary regime, the master equations reduce to the following system for the length-resolved state densities:

$$\frac{\partial A_{00}}{\partial \tilde{x}} = -A_{00}\Big(1+\gamma_c+\gamma_b\Big) + \Big[A_c + \alpha A_b\Big], \tag{75}$$

$$\frac{\partial A_c}{\partial \tilde{x}} = 2\gamma A_{00} - \Big(4+\gamma_c+\gamma_b\Big)A_c + 2\Big(2A_{cc} + \alpha A_{bc}\Big), \tag{76}$$

$$\frac{\partial A_b}{\partial \tilde{x}} = 2\gamma_b A_{00} - \Big(2+2\alpha+\gamma_c+\gamma_b\Big)A_b + 2\Big(A_{bc} + 2\alpha A_{bb}\Big). \tag{77}$$

The algebraic constraints give the blocked–state densities in terms of $A_b, A_c$:

$$A_{cc} = \frac{\gamma_c}{6}\,A_c, \tag{78}$$

$$A_{bb} = \frac{\gamma_b}{2(1+2\alpha)}\,A_b, \tag{79}$$

$$A_{bc} = \frac{\gamma_b A_c + \gamma_c A_b}{2(2+\alpha)}. \tag{80}$$

Substituting these into (76)–(77) yields the closed system

$$\frac{\partial A_{00}}{\partial \tilde{x}} = -A_{00}\Big(1+\gamma_c+\gamma_b\Big) + \Big[A_c + \alpha A_b\Big], \tag{81}$$

$$\frac{\partial A_c}{\partial \tilde{x}} = 2\gamma A_{00} - \left(4+\frac{\gamma_c}{3}+\frac{2\gamma_b}{2+\alpha}\right)A_c + \frac{\alpha\,\gamma_c}{2+\alpha}\,A_b, \tag{82}$$

$$\frac{\partial A_b}{\partial \tilde{x}} = 2\gamma_b A_{00} - \left(2+2\alpha+\frac{\gamma_b}{1+2\alpha}+\frac{(1+\alpha)\gamma_c}{2+\alpha}\right)A_b + \frac{\gamma_b}{2+\alpha}\,A_c. \tag{83}$$

For $\tilde{x} > 0$ equations (81)–(83) can be written as a linear system

$$\frac{d}{d\tilde{x}}\begin{pmatrix} A_{00} \\ A_c \\ A_b \end{pmatrix} = \mathbb{M}(\gamma,\gamma_b,\alpha)\begin{pmatrix} A_{00} \\ A_c \\ A_b \end{pmatrix}. \tag{84}$$

Let $\{\beta_1, \beta_2, \beta_3\}$ be the eigenvalues of $-\mathbb{M}$ (so that each decay rate $\beta_m > 0$ produces a term $e^{-\beta_m \tilde{x}}$ in the solution). If all eigenvalues are different, the general solution reads

$$\begin{pmatrix} A_{00} \\ A_c \\ A_b \end{pmatrix} = \sum_{m=1}^{3} B_m \mathbf{u_m} \, e^{-\beta_m \tilde{x}}, \tag{85}$$

where $\mathbf{u_m}$ are the eigenvectors of $\mathbb{M}$, with $A_{bb}, A_{cc}, A_{bc}$ recovered algebraically. The coefficients $B_m$ are fixed by boundary/normalization conditions (e.g. regularity as $\tilde{x} \to \infty$ and a chosen normalization of the total time–integrated probability).

Solving the characteristic polynomial of $\mathbb{M}$ gives three eigenvalues. In compact closed form they can be written as

$$\beta_1 = \frac{3 + \gamma_c + 2\alpha(3 + \gamma_c) + 3\gamma_b}{3 + 6\alpha}, \tag{86}$$

$$\beta_2 = S_1 + S_2, \tag{87}$$

$$\beta_3 = S_1 - S_2. \tag{88}$$

with

$$S_1 = \frac{12 + 4\alpha^3 + 3\gamma_c + 4\gamma_b + 2\alpha^2(11 + 2\gamma_c + \gamma_b) + \alpha(34 + 8\gamma_c + 9\gamma_b)}{2(2 + \alpha)(1 + 2\alpha)} \tag{89}$$

$$S_2 = \frac{(1 + 2\alpha)}{2(2 + \alpha)(1 + 2\alpha)} \times \tag{90}$$

$$\sqrt{4\alpha^4 + (4 + \gamma_c)^2 + 2\alpha\gamma_c(-2 + \gamma_b) - 8\alpha(2 + \gamma_b) + 4\alpha^3(2 + \gamma_b) + \alpha^2\Big(-12 - 4\gamma_c + 4\gamma_b + \gamma_b^2\Big)}, \tag{91}$$

In the small–$\alpha$ regime ($\alpha \ll 1$), these admit regular expansions;

$$\beta_1 = \left(\frac{\gamma_c}{3} + \gamma_b + 1\right) - 2\alpha\gamma_b \tag{92}$$

$$\beta_2 = (\gamma_c + \gamma_b + 4) - \frac{2\alpha\gamma_b}{\gamma + 4} \tag{93}$$

$$\beta_3 = \left(\frac{\gamma_c}{2} + \gamma_b + 2\right) + \frac{\alpha\,(\gamma_c^2 - 2\gamma(\gamma_b - 6) + 32)}{4(\gamma_c + 4)} \tag{94}$$

As in the previous example, one can verify by direct calculation that the decay constants $\beta_i$ do not become degenerate for any physical values of the parameters. Indeed, imposing pairwise equalities

$$\beta_1 = \beta_2, \qquad \beta_2 = \beta_3, \qquad \beta_1 = \beta_3$$

and solving any of these equations with respect to one of the model parameters (for example, $\gamma$) shows that all resulting roots are negative whenever $\gamma_b > 0$ and $\alpha > 0$.

Hence, in the physical parameter region the three decay constants remain distinct, and the stationary solution is a genuine sum of three simple exponentials, without polynomial prefactors.

Finally we obtain the expression for the distribution

$$N(\tilde{x}) = \sum_{m=1}^{3} C_m \, e^{-\beta_m \tilde{x}}, \tag{95}$$

where $C_m$ are the coefficients, directly obtained from $B_m$, according to

$$C_m = B_m \sum_{i=1}^{3} u_m^i (1 + F_i), \tag{96}$$

where we sum over the dynamic states of the system ($i = 1$ corresponds to $A_{00}$, $i = 2$ corresponds to $A_b$, $i = 3$ corresponds to $A_c$), and the factor $F_i$ expresses the algebraic connection between dynamic and blocked states. Explicitly, $F_i$ reads as

$$F_1 = 0, \tag{97}$$

$$F_2 = \frac{\gamma_b}{2(1 + 2\alpha)} + \frac{(\gamma + \gamma_{\text{dyn}})}{2(2 + \alpha)}, \tag{98}$$

$$F_3 = \frac{\gamma + \gamma_{\text{dyn}}}{6} + \frac{\gamma_b}{2(2 + \alpha)} \tag{99}$$

**Peak of the distribution** A natural question is under which conditions the stationary loop-length distribution develops a local maximum at finite $\tilde{x}$, rather than decaying monotonically from $\tilde{x} = 0$. In the present problem, the total loop length is the sum of two arm lengths, each of which is exponentially distributed; in general, the two arm lengths are correlated, and therefore the appearance of a peak in the distribution is not determined by a simple criterion.

In the simplest case of one type of static barrier and opaque cohesins (see previous paragraph), however, the stationary distribution is available in closed form. For this distribution (the sum of three exponentials), condition for the existence of a peak at finite $\tilde{x}$ is equivalent to the condition that the derivative at the origin be positive, i.e.

$$\left. \frac{dN(\tilde{x})}{d\tilde{x}} \right|_{\tilde{x}=0} > 0. \tag{100}$$

which immediately gives

$$\sum_{m=1}^{3} C_m \beta_m < 0. \tag{101}$$

Substituting the expressions for $C_m$ and $\beta_m$ obtained above, one finds that a peak is

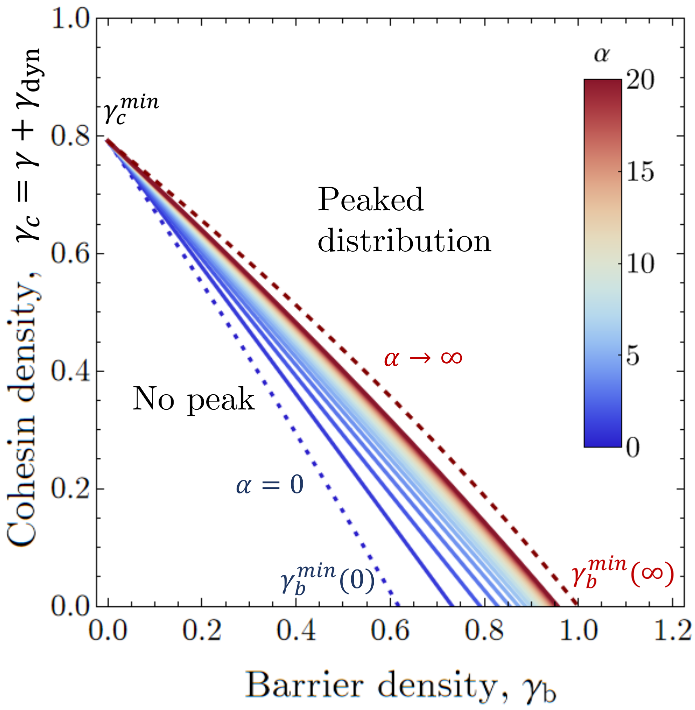

Figure S2: Phase diagram for the emergence of a peak in the stationary loop-length distribution. Solid lines show the boundary in the $(\gamma_c, \gamma_b)$ plane separating parameter regimes with and without a finite-$\tilde{x}$ maximum in the loop-length PDF, for different values of the barrier lifetime parameter $\alpha$, color-coded as indicated. Above the boundary, the stationary distribution is peaked, whereas below it the distribution decays monotonically from the origin. The dashed blue and red lines denote the limiting cases $\alpha = 0$ and $\alpha \to \infty$, respectively. The corresponding intercepts define the threshold values $\gamma_b^{\min}(0)$, $\gamma_b^{\min}(\infty)$, and $\gamma_c^{\min}$.

present provided

$$\gamma + \gamma_{\mathrm{dyn}} > \frac{\sqrt{3}F - 3\alpha(2\alpha + 4\gamma_b + 5) - 6\gamma_b - 6}{2(\alpha + 2)(2\alpha + 1)}. \tag{102}$$

where $F = \sqrt{(2\alpha + 1)\left(-4(2\alpha + 1)\left(\alpha^2 + \alpha - 2\right)\gamma_b - 4(\alpha - 1)^2\gamma_b^2 + 7(2\alpha + 1)(\alpha + 2)^2\right)}$.

The corresponding phase diagram is shown in Fig. S2.

Several limiting cases are worth noting. In the absence of static barriers ($\gamma_b = 0$), the minimal value of $\gamma_c = \gamma + \gamma_{\mathrm{dyn}}$ required for the appearance of a peak is

$$\gamma_c > \frac{\sqrt{21} - 3}{2} \approx 0.79. \tag{103}$$

Conversely, in the limit $\gamma_c \to 0$, the critical barrier density required for peak formation depends on $\alpha$. For long-lived barriers ($\alpha = 0$), one finds the threshold

$$\gamma_b > \varphi^{-1} = \frac{\sqrt{5} - 1}{2} \approx 0.62, \tag{104}$$

where $\varphi$ is the golden ratio. In the opposite limit $\alpha \to \infty$, corresponding to short-lived

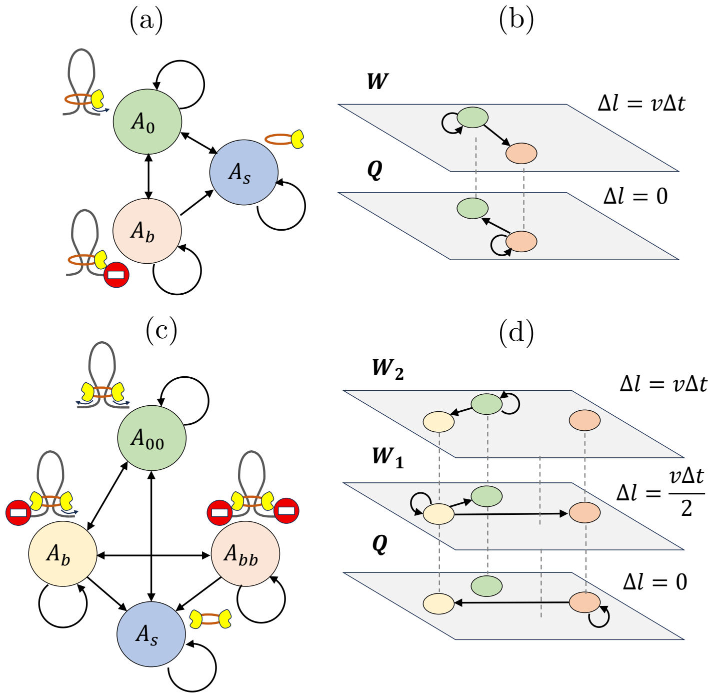

Figure S3: Graph and multigraph representations of one- and two-sided extrusion and a single obstacle class (either a barrier or another cohesin). (a) Transition graph for one-sided extrusion. $A_0$ denotes the freely extruding state, $A_b$ the blocked state, and $A_s$ the soluble cohesin pool. (b) Corresponding multigraph for one-sided extrusion. The transitions in panel (a) are separated into two layers according to the loop-length increment $\Delta l$: in layer $W$, transitions out of the mobile state ($A_0 \to A_0$, $A_0 \to A_b$) increase the loop length by $\Delta l = v\Delta t$; in layer $Q$, transitions out of the blocked state ($A_b \to A_b$, $A_b \to A_0$) leave the loop length unchanged, $\Delta l = 0$. (c) Transition graph for symmetric two-sided extrusion. $A_{00}$ denotes the state with both arms mobile, $A_b$ the state with one blocked arm, $A_{bb}$ the fully blocked state, and $A_s$ the soluble cohesin pool. (d) Corresponding multigraph for two-sided extrusion. The transitions in panel (c) are separated into three layers according to $\Delta l$: in layer $W_2$, both arms are mobile and the loop grows by $\Delta l = v\Delta t$; in layer $W_1$, only one arm remains mobile and the loop grows by $\Delta l = v\Delta t/2$; in layer $Q$, both arms are blocked and $\Delta l = 0$.

barriers, the critical value tends to

$$\gamma_b \to 1. \tag{105}$$

In this regime, however, the peak simultaneously approaches $x = 0$ and becomes progressively less pronounced, so that it ultimately loses practical observability.

#### 3.2 Mapping onto a graph

The kinetic equations derived above are convenient for obtaining exact stationary solutions, but they hide the simple geometric origin of the resulting loop-length distributions. A complementary viewpoint is to interpret the dynamics as transport on a discrete state graph: nodes represent internal cohesin states, while edges represent allowed transitions during one time step $\Delta t$ (Fig. S3a,c). The transition rules themselves were introduced in Sec. 1 (*Transitions between states*).

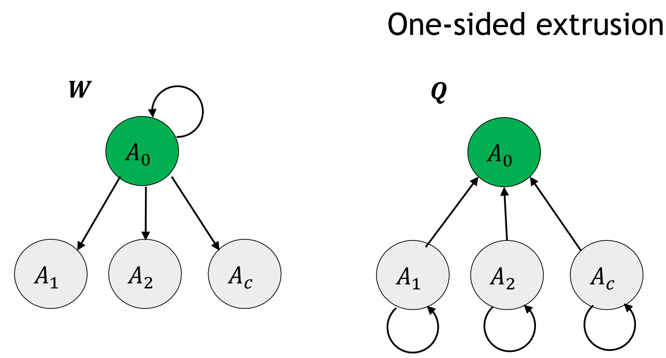

Figure S4: **W**-, **Q**-, and **L**-graph representations for one-sided extrusion with $n = 2$ barrier classes. Nodes denote the components of $\mathbf{A} = (A_0, A_1, A_2, A_c)^\mathsf{T}$: $A_0$ is the free state, $A_1$ and $A_2$ are states blocked by the two barrier classes, and $A_c$ is the state blocked by another cohesin. Nonzero edges are determined by the physical role of each graph: **W** and **L** advance the loop length $l$, whereas **Q** acts at fixed $l$.

This graph representation makes the origin of the stationary distributions transparent. In particular, it immediately explains why one-sided extrusion always produces a single exponential mode, whereas two-sided extrusion generically yields a sum of exponentials. The same construction also allows one to compare different microscopic implementations of two-sided extrusion—such as fully symmetric motion, random switching of the active leg, or collision-induced switching—within a common framework.

##### 3.2.1 Unidirectional extrusion

We first recast the one-sided extrusion model with $n + 1$ obstacle classes ($n$ classes of barriers plus the cohesin class) as a discrete-time transport process on a directed graph.

During a short time step $\Delta t$, the active arm advances by

$$\Delta l = v\,\Delta t$$

provided it remains mobile. To describe the dynamics with length resolution, we restrict attention to chromatin-bound states and introduce the vector

$$\mathbf{A}(l,t) = \Big(A_0(l,t), A_1(l,t), A_2(l,t), \ldots, A_{n+1}(l,t)\Big)^\mathsf{T}, \tag{106}$$

where $A_0(l,t)$ is the probability density of being in the free (moving) state with loop length $l$ at time $t$, and $A_i(l,t)$ is the density of being blocked by an obstacle of class $i$. The soluble pool $A_s(t)$ is not included explicitly in this length-resolved sector.

Within one time step, two types of transitions are possible. First, a mobile arm may encounter an obstacle and become blocked. Such *encounter transitions* are necessarily accompanied by loop growth, because the arm must advance in order to reach the obstacle. Second, a blocking obstacle may dissociate, thereby restoring the free state. Such *release transitions* do not change the loop length.

This naturally defines a layered representation of the state graph (a multigraph): transitions accompanied by loop growth are collected into the layer $\mathbf{W}$, whereas transitions at fixed loop length are collected into the layer $\mathbf{Q}$, see Fig. S3b. In this way, the original time-evolution graph (in Fig. S3a) is decomposed into an advective part and a conservative part. Cohesin dissociation can occur from any state and is therefore incorporated as a common survival factor in the corresponding propagators.

The one-step update for the bound-state sector can then be written as

$$\mathbf{A}(l+\Delta l, t+\Delta t) = \mathbf{W}\,\mathbf{A}(l,t) + \mathbf{Q}\,\mathbf{A}(l+\Delta l, t). \tag{107}$$

Here $\mathbf{W}$ contains transitions associated with forward motion from $l$ to $l+\Delta l$, namely propagation of the free state together with possible blocking during advection. By contrast, $\mathbf{Q}$ contains transitions at fixed loop length, namely release events that do not change $l$. Each transition carries the common factor $1-\Delta t/\tau$, equal to the probability that cohesin remains bound during the step.

Let $\gamma_i = v\tau/d_i$ be the dimensionless density of obstacle class $i$ and $\alpha_i = \tau/\tau_i$ the persistence ratio, so that the obstacle release rate is $1/\tau_i = \alpha_i/\tau$. To leading order in $\Delta t$, the encounter rate with class $i$ is $v/d_i = \gamma_i/\tau$, hence the probability to become blocked by class $i$ during $\Delta t$, conditioned on cohesin remaining bound, is $(\gamma_i/\tau)\Delta t$.

With this convention, Eq. (107) is specified by

$$\mathbf{W} = \left(1-\frac{\Delta t}{\tau}\right)\begin{pmatrix} 1-\frac{\Delta t}{\tau}\sum_{i=1}^{n+1}\gamma_i & 0 & 0 & \cdots & 0 \\ \frac{\Delta t}{\tau}\gamma_1 & 0 & 0 & \cdots & 0 \\ \frac{\Delta t}{\tau}\gamma_2 & 0 & 0 & \cdots & 0 \\ \vdots & \vdots & \vdots & \ddots & \vdots \\ \frac{\Delta t}{\tau}\gamma_{n+1} & 0 & 0 & \cdots & 0 \end{pmatrix}, \tag{108}$$

and

$$\mathbf{Q} = \left(1-\frac{\Delta t}{\tau}\right)\begin{pmatrix} 0 & \frac{\Delta t}{\tau_1} & \frac{\Delta t}{\tau_2} & \cdots & \frac{\Delta t}{\tau_{n+1}} \\ 0 & 1-\frac{\Delta t}{\tau_1} & 0 & \cdots & 0 \\ 0 & 0 & 1-\frac{\Delta t}{\tau_2} & \cdots & 0 \\ \vdots & \vdots & \vdots & \ddots & \vdots \\ 0 & 0 & 0 & \cdots & 1-\frac{\Delta t}{\tau_{n+1}} \end{pmatrix}. \tag{109}$$

The sum of each column equals the survival factor $1-\Delta t/\tau$, i.e., the total probability to remain bound upon leaving a given node.

In the stationary regime,

$$\mathbf{A}(l+\Delta l, t+\Delta t) = \mathbf{A}(l+\Delta l, t),$$

so Eq. (107) reduces to

$$\mathbf{A}(l+\Delta l) = (\mathbf{I}-\mathbf{Q})^{-1}\mathbf{W}\,\mathbf{A}(l) \equiv \mathbf{L}\,\mathbf{A}(l). \tag{110}$$

To first order in $\Delta l$, this gives

$$\mathbf{L} = \begin{pmatrix} 1-\left(1+\sum_i \frac{\gamma_i}{1+\alpha_i}\right)\frac{\Delta l}{l_p} & 0 & 0 & \cdots & 0 \\ \frac{\gamma_1}{1+\alpha_1} & 0 & 0 & \cdots & 0 \\ \frac{\gamma_2}{1+\alpha_2} & 0 & 0 & \cdots & 0 \\ \vdots & \vdots & \vdots & \ddots & \vdots \\ \frac{\gamma_{n+1}}{1+\alpha_{n+1}} & 0 & 0 & \cdots & 0 \end{pmatrix} + \mathcal{O}(\Delta l^2). \tag{111}$$

The loop-length problem is therefore mapped onto a survival process on the **L**-graph, with the initial node $A_0$ at $l=0$. Since only transitions out of the free state increase the loop length, the graph has a single propagating node and **L** has only one nonzero column (Fig. S4). This topology alone implies that the survival probability $N(l)$ has a single propagating mode and therefore a single exponential form. In other words, for one-sided extrusion the exponential loop-length distribution follows directly from the graph structure, independently of the detailed weights of **L**.

**Formal solution of the L-graph equation.** The transfer equation (110) is equivalent to the discrete kinetic relations

$$A_0(l+\Delta l) = \left[1-\left(1+\sum_i \frac{\gamma_i}{1+\alpha_i}\right)\frac{\Delta l}{l_p}\right]A_0(l), \tag{112}$$

$$A_i(l+\Delta l) = \frac{\gamma_i}{1+\alpha_i}\,A_0(l). \tag{113}$$

The prefactor in the first equation is the probability to avoid all termination events over one step $\Delta l$: the first contribution corresponds to cohesin dissociation, and the remaining ones to blocking by obstacle classes $i$. Thus the first equation is simply the survival law for the free state $A_0(l) = N(l)$.

Taking the limit $\Delta l \to 0$ gives

$$\frac{dN(l)}{dl} = -\frac{1}{l_p}\left(1+\sum_{i=1}^{n+1}\frac{\gamma_i}{1+\alpha_i}\right)N(l),$$

while the blocked populations become algebraically related to the free one,

$$A_i(l) = \frac{\gamma_i}{1+\alpha_i}N(l).$$

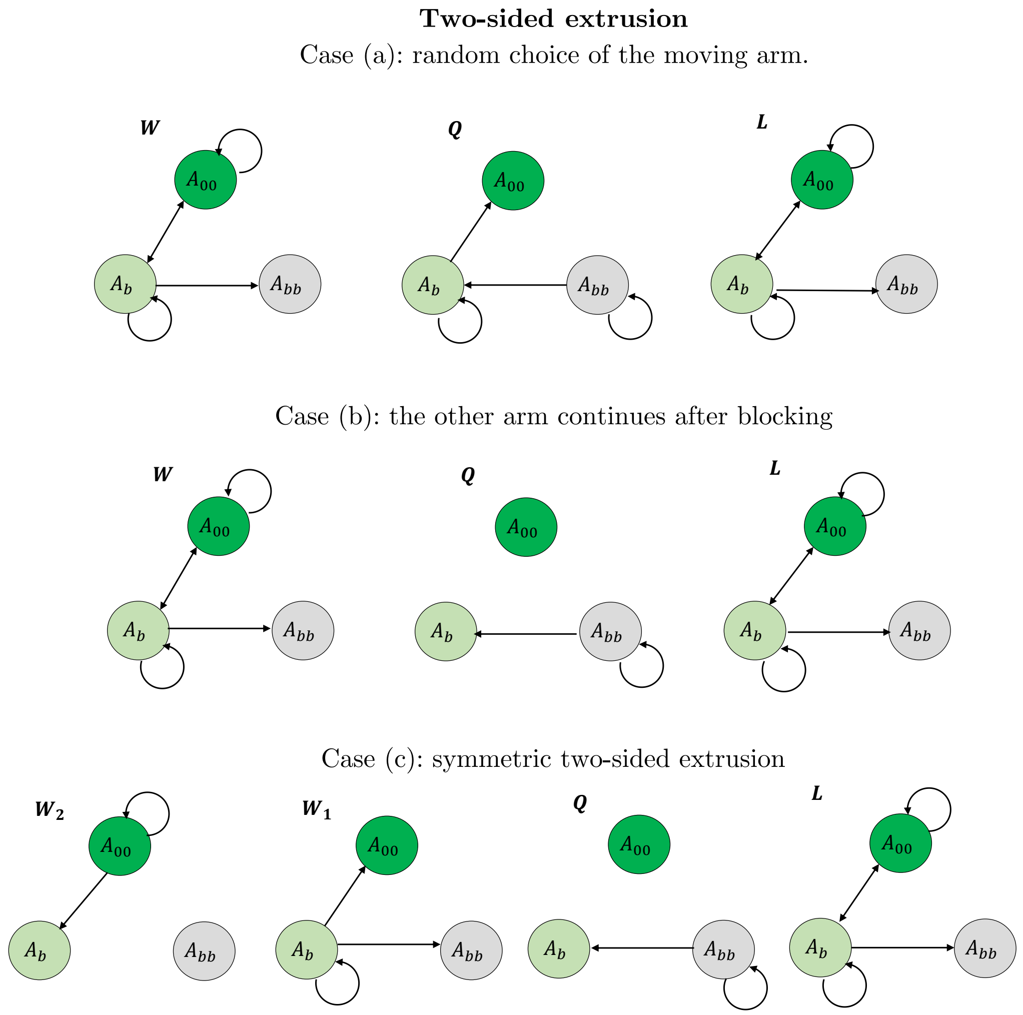

Figure S5: Two-sided extrusion under different rules of arm motion after a blocking event. Schematic representation of **W** and **Q** layers of transitions and finite **L**-graph for three scenarios for bidirectional loop extrusion. (a) Random choice of the moving arm: after loading, only one arm is advanced at each step, chosen stochastically. (b) Continued motion of the unblocked arm: if one arm is stalled by an obstacle, the other arm continues extruding. (c) Symmetric two-sided extrusion: both arms move simultaneously and contribute equally to loop growth.

Passing to the dimensionless coordinate $\tilde{x} = l/l_p$ reproduces the continuous equations (50–51) and the exponential solution with the mean length $\tilde{\lambda}$, Eq.(28).

##### 3.2.2 Bidirectional extrusion

The graph representation is particularly useful for bidirectional extrusion because the topology of the directed length-state graph **L** directly determines the structure of the loop-length distribution. In this representation, each state that still propagates in loop length contributes an independent dynamical mode. Accordingly, if the **L**-graph contains $m$ propagating nodes, the stationary distribution $N(x)$ is a linear combination of $m$ exponentials. The qualitative difference from one-sided extrusion therefore follows immediately from graph topology: one-sided extrusion has a single propagating node and hence a single exponential, whereas bidirectional extrusion generically contains additional propagating blocked states and therefore multiple exponential modes.

We now recast *bidirectional* extrusion in the same graph language and compare three microscopic implementations:

*(a) Alternating arm updates (random choice).* At each time step $\Delta t$, exactly one arm advances by $\Delta l = v\,\Delta t$, with the moving arm (left or right) chosen at random.

*(b) Backup arm (blocking-triggered switching).* Once one arm becomes blocked by an obstacle, the other arm continues advancing alone until it is blocked or the first arm is released.

*(c) Symmetric two-arm growth.* When unblocked, *both* arms advance simultaneously by $\Delta l/2$, so that the total loop length increases by $\Delta l$ per step. This is the symmetric two-sided mechanism used in the main text.

To illustrate the common structure of these mechanisms, we consider the simplest case: transparent cohesins and a single static barrier class $B$ with parameters $(\gamma_b, \alpha)$. In all three bidirectional variants (a)–(c), the corresponding **L**-graph contains two propagating nodes, associated with the free state and the singly blocked state (green nodes in Fig. S5). The stationary loop-length distribution must therefore contain two exponential modes,

$$N(\tilde{x}) = C_1\, e^{-\beta_1 \tilde{x}} + C_2\, e^{-\beta_2 \tilde{x}}, \tag{114}$$

where the decay rates $\beta_{1,2}$ are related to the eigenvalues $\nu_{12}$ of the **L**-graph (see below). Thus, the emergence of multiexponential, and generically peaked, loop-length statistics in bidirectional extrusion is a direct topological consequence of the **L**-graph.

**Case (a): random choice of the moving arm.** Here we consider the update rule: during each time step only one arm advances by $\Delta l$ (left or right with probability 1/2). Further for simplicity we consider the discrete increment $\Delta = \Delta l/l_p$ and a three–component state vector

$$\mathbf{A}(\tilde{x}) = \Big(A_{00}(\tilde{x}),\, A_b(\tilde{x}),\, A_{bb}(\tilde{x})\Big)^{\mathsf{T}},$$

where $A_{00}$ is the free (both arms mobile) state, $A_b$ is the one–sided blocked state, $A_{bb}$ is the fully blocked state. In this variant the one–step stationary update can be written as $\mathbf{A}(\tilde{x}+\Delta) = \mathbf{W}\,\mathbf{A}(\tilde{x}) + \mathbf{Q}\,\mathbf{A}(\tilde{x}+\Delta)$, cf. Eq. (107). The corresponding matrices are

$$\mathbf{W} = (1-\Delta)\begin{pmatrix} 1-\gamma_b\Delta & \frac{1}{2}\alpha\Delta & 0 \\ \gamma_b\Delta & \frac{1}{2}\Big(1-\Delta(\frac{1}{2}\gamma_b+\alpha)\Big) & 0 \\ 0 & \frac{1}{2}\gamma_b\Delta & 0 \end{pmatrix}, \tag{115}$$

$$\mathbf{Q} = (1-\Delta)\begin{pmatrix} 0 & \frac{1}{2}\alpha\Delta & 0 \\ 0 & \frac{1}{2}\Big(1-\Delta(\frac{1}{2}\gamma_b+\alpha)\Big) & 2\alpha\Delta \\ 0 & 0 & 1-2\alpha\Delta \end{pmatrix}. \tag{116}$$

As in Sec. 3.2.1, stationarity implies $\mathbf{A}(\tilde{x}+\Delta) = (\mathbf{I}-\mathbf{Q})^{-1}\mathbf{W}\,\mathbf{A}(\tilde{x}) \equiv \mathbf{L}\,\mathbf{A}(\tilde{x})$. Expanding to first order in $\Delta$ gives

$$\mathbf{L} \;=\; \begin{pmatrix} 1-(1+\gamma_b)\Delta & \alpha\Delta & 0 \\ 2\gamma_b\Delta & 1-\dfrac{2+6\alpha+4\alpha^2+\gamma_b}{1+2\alpha}\Delta & 0 \\ 0 & \dfrac{\gamma_b}{2(1+2\alpha)} & 0 \end{pmatrix} \;+\; \mathcal{O}(\Delta^2), \tag{117}$$

The two nontrivial eigenvalues of $\mathbf{L}$ are

$$\nu_1 = 1-(2+2\alpha+\gamma_b)\,\Delta, \quad \nu_2 = 1-\left(1+\frac{\gamma_b}{1+2\alpha}\right)\Delta \tag{118}$$

Passing to the continuum limit $\Delta \to 0$, an eigenvalue of the form $\nu = 1-\beta\,\Delta$ generates a mode $\propto e^{-\beta\tilde{x}}$. Therefore, the two decay constants (the "exponents") are

$$\beta_1 = 2+2\alpha+\gamma_b, \quad \beta_2 = 1+\frac{\gamma_b}{1+2\alpha} < \beta_1 \tag{119}$$

Accordingly, the stationary loop–length distribution is a linear combination of two exponentials, $N(\tilde{x}) = C_1 e^{-\beta_1\tilde{x}} + C_2 e^{-\beta_2\tilde{x}}$. Importantly, the decay constants $\beta_{1,2}$ coincide exactly with those obtained for the same case (transparent cohesin and one class of obstacle) in Sec.3.1.2 for symmetric bidirectional extrusion (equation (70)). Thus, the two models are statistically indistinguishable: symmetric bidirectional extrusion and extrusion with random leg switching share the same stationary statistics.

However we should note that the limit $\Delta \to 0$ used above corresponds to sufficiently rapid switching between the two arms relative to the cohesin residence time. Thus, the equivalence with symmetric two-sided extrusion applies only in the rapid-switching regime. For slower switching, individual trajectories remain effectively one-sided.

**Case (b): the other arm continues after blocking.** In this variant, arms also cannot work simultaneously, but the switching between arms is not random, it happens, when the growing arm collides with an obstacle. Using the same three–state basis $\mathbf{A}(\tilde{x}) = \big(A_{00}(\tilde{x}), A_b(\tilde{x}), A_{bb}(\tilde{x})\big)^{\mathsf{T}}$, the stationary one–step update $\mathbf{A}(\tilde{x}+\Delta) = \mathbf{W}\,\mathbf{A}(\tilde{x}) + \mathbf{Q}\,\mathbf{A}(\tilde{x}+\Delta)$ is characterized by

$$\mathbf{W} = (1-\Delta)\begin{pmatrix} 1-\Delta\gamma_b & \alpha\Delta & 0 \\ \gamma_b\Delta & 1-\Delta(\gamma_b+\alpha) & 0 \\ 0 & \gamma_b\Delta & 0 \end{pmatrix}, \qquad \mathbf{Q} = (1-\Delta)\begin{pmatrix} 0 & 0 & 0 \\ 0 & 0 & 2\alpha\Delta \\ 0 & 0 & 1-2\alpha\Delta \end{pmatrix}. \tag{120}$$

As before,

$$\mathbf{A}(\tilde{x}+\Delta) = (\mathbf{I}-\mathbf{Q})^{-1}\mathbf{W}\,\mathbf{A}(\tilde{x}) \equiv \mathbf{L}\,\mathbf{A}(\tilde{x}).$$

Expanding to first order in $\Delta$ yields

$$\mathbf{L} = \begin{pmatrix} 1-(1+\gamma_b)\Delta & \alpha\Delta & 0 \\ \gamma_b\Delta & 1-\dfrac{(1+3\alpha+2\alpha^2+\gamma_b)}{1+2\alpha}\Delta & 0 \\ 0 & \dfrac{\gamma_b}{1+2\alpha} & 0 \end{pmatrix} + \mathcal{O}(\Delta^2). \tag{121}$$

which matches the discrete operator quoted above.

The two nontrivial eigenvalues of $\mathbf{L}$ can be written in the compact form

$$\nu_{1,2} = 1 - \frac{\Delta}{2(1+2\alpha)}\Big(P \pm S\Big),$$

where

$$P = 2+5\alpha+2\alpha^2+2(1+\alpha)\gamma_b,$$

$$S = \sqrt{\alpha\Big(4\alpha\gamma_b^2+4(\alpha+1)(2\alpha+1)\gamma_b+\alpha(2\alpha+1)^2\Big)}.$$

The third eigenvalue is 0.

Although the expressions for $\nu_m$ and corresponding expressions for $\beta_m$ are more complex in the case (b) relative to case (a) and bidirectional symmetric extrusion, the functional shape of distributions remains the same - the solution is just the sum of two exponentials.

**Case (c): symmetric two–sided extrusion** In the symmetric bidirectional scheme, a loop can advance by one half of nondimensional arm step $\Delta/2$ (one arm moves) or by $\Delta$ (both arms move), depending on the internal state. In variants (a) and (b), the formalism was identical to the unidirectional case of Sec. 3.2.1. In variant (c) the discrete update involves *two* spatial increments because probability can arrive at length $\tilde{x}+\Delta$ either by (i) two-arm advancement from $l$ in one step, or (ii) one-arm advancement to $\tilde{x}+\Delta/2$ followed by the second-arm advancement to $\tilde{x}+\Delta$. This is conveniently written as a stationary two-step transfer in loop length:

$$\mathbf{A}(\tilde{x}+\Delta) = \mathbf{Q}\,\mathbf{A}(\tilde{x}+\Delta) + \mathbf{W}_1\,\mathbf{A}(\tilde{x}+\Delta/2) + \mathbf{W}_2\,\mathbf{A}(\tilde{x}), \tag{122}$$

where $\mathbf{A} = (A_{00}, A_b, A_{bb})^{\mathsf{T}}$ collects the densities of the free state, the one-sided blocked state, and the fully blocked state, respectively, which can be represented as a three-layer multigraph (Fig. S3d). For a single barrier class, the discrete-time matrices are:

$$\mathbf{W}_2 = (1-\Delta)\begin{pmatrix} 1-\gamma_b\Delta & 0 & 0 \\ \Delta\gamma_b & 0 & 0 \\ 0 & 0 & 0 \end{pmatrix}, \qquad \mathbf{W}_1 = (1-\Delta)\begin{pmatrix} 0 & \alpha\Delta & 0 \\ 0 & 1-\frac{1}{2}\gamma_b\Delta-\alpha\Delta & 0 \\ 0 & \frac{1}{2}\Delta\gamma_b & 0 \end{pmatrix}, \tag{123}$$

$$\mathbf{Q} = (1-\Delta)\begin{pmatrix} 0 & 0 & 0 \\ 0 & 0 & 2\alpha\Delta \\ 0 & 0 & 1-2\alpha\Delta \end{pmatrix}. \tag{124}$$

Here $\mathbf{Q}$ encodes transitions at *fixed* loop length, while $\mathbf{W}_1$ and $\mathbf{W}_2$ encode updates that occur while attempting to advance the loop length by $\Delta/2$ and $\Delta$, respectively. Equation (122) reduces to

$$(\mathbf{I}-\mathbf{Q})\,\mathbf{A}(\tilde{x}+\Delta) = \mathbf{W}_1\,\mathbf{A}(\tilde{x}+\Delta/2) + \mathbf{W}_2\,\mathbf{A}(\tilde{x}). \tag{125}$$

To pass to a continuum description in loop-length space, we expand $\mathbf{A}(\tilde{x}+\Delta)$ and $\mathbf{A}(\tilde{x}+\Delta/2)$ to first order in $\Delta$:

$$\mathbf{A}(\tilde{x}+\Delta) \simeq \mathbf{A}(\tilde{x}) + \Delta\,\frac{d\mathbf{A}}{d\tilde{x}}, \qquad \mathbf{A}\left(\tilde{x}+\frac{\Delta}{2}\right) \simeq \mathbf{A}(\tilde{x}) + \frac{\Delta}{2}\,\frac{d\mathbf{A}}{d\tilde{x}}.$$

Substituting these expansions into Eq. (125) gives

$$(\mathbf{I}-\mathbf{Q})\left(\mathbf{A}(\tilde{x}) + \Delta\,\frac{d\mathbf{A}}{d\tilde{x}}\right) = \mathbf{W}_1\left(\mathbf{A}(\tilde{x}) + \frac{\Delta}{2}\,\frac{d\mathbf{A}}{d\tilde{x}}\right) + \mathbf{W}_2\,\mathbf{A}(\tilde{x}),$$

Collecting the derivative terms, one obtains a first-order ODE in $\tilde{x}$:

$$\Delta\,\frac{d\mathbf{A}}{d\tilde{x}} = \left(\mathbf{I}-\mathbf{Q}-\frac{1}{2}\mathbf{W_1}\right)^{-1}(\mathbf{W_1}+\mathbf{W_2}+\mathbf{Q}-\mathbf{I})\,\mathbf{A}(\tilde{x}). \tag{126}$$

This naturally results in the following $\mathbf{L}$-graph operator:

$$\mathbf{L} = \left(\mathbf{I}-\mathbf{Q}-\frac{1}{2}\mathbf{W_1}\right)^{-1}(\mathbf{W_1}+\mathbf{W_2}+\mathbf{Q}-\mathbf{I}) + \mathbf{I}, \tag{127}$$

so that the continuum equation takes the compact form

$$\frac{d\mathbf{A}}{d\tilde{x}} = \frac{1}{\Delta}\Big(\mathbf{L}-\mathbf{I}\Big)\,\mathbf{A}(\tilde{x}), \tag{128}$$

corresponding to the discrete update $\mathbf{A}(\tilde{x}+\Delta) = \mathbf{L}\;\mathbf{A}(\tilde{x})$.

For subsequent analytical manipulations we should eliminate the fully blocked node $A_{bb}$. The dynamics reduces to a closed $2\times 2$ transfer on the moving subspace $(A_{00}, A_b)$, and the corresponding continuum generator has two eigenvalues. Equivalently, the loop-length PDF is a sum of two exponentials as in Eq. (114).

Keeping only the leading (first-order) terms in $\Delta$, one obtains the reduced transfer matrix for a single effective step $\Delta$ in the compact form

$$\mathbf{L}-\mathbf{I} \approx \Delta\begin{pmatrix} -(1+\gamma_b) & \alpha \\ 2\gamma_b & -\left(2+2\alpha+\frac{\gamma_b}{1+2\alpha}\right) \end{pmatrix}, \tag{129}$$

Passing to the continuum limit, we identify the decay rates (the exponential slopes in $N(\tilde{x})$):

$$\beta_1 = 2(1+\alpha) + \gamma_b, \qquad \beta_2 = 1 + \frac{\gamma_b}{1+2\alpha}. \tag{130}$$

which coincide with the result for the case of random arm switching (a), Eq.(119).

### 4. Correlation between the arms

The loop-length distributions discussed above already imply that, in the biologically relevant regime, two cohesin arms cannot be treated as fully independent. In particular, the blocked-loop PDF is generically peaked but does not reduce to the ideal Γ-distribution expected for two completely decoupled exponential arms, except in a singular limit of infinitely dense permanent barriers. This naturally raises the next question: how strongly are the two extrusion fronts correlated, and which obstacles control this correlation? To answer this, we now analyze the arm–arm correlation in the minimal two-sided model containing one class of static barriers and other cohesins, both acting as blockers.

We return to the two-coordinate formulation introduced in Sec. 2.2, in which the left and right arms are described separately by their dimensionless extensions $\tilde{\ell}$ and $\tilde{r}$. We specialize to the minimal biologically relevant case discussed in Sec. 3.2.2 (*Example: one obstacle + opaque cohesin*), namely a single static barrier class $b$ and other cohesins $c$, also treated as barriers. The stationary time-integrated equations then read

$$\frac{1}{2}\frac{\partial A_{00}}{\partial \tilde{\ell}} + \frac{1}{2}\frac{\partial A_{00}}{\partial \tilde{r}} = N_b \delta(\tilde{r}, \tilde{\ell}) - A_{00}\Big(1 + \Gamma_{\text{eff}}\Big) + (A_{c0} + A_{0c}) + \alpha(A_{b0} + A_{0b}), \tag{131}$$

$$\frac{1}{2}\frac{\partial A_{0c}}{\partial \tilde{\ell}} = \frac{\gamma_c}{2} A_{00} - \Big(2 + \tfrac{\Gamma_{\text{eff}}}{2}\Big) A_{0c} + A_{cc} + \alpha A_{bc}, \tag{132}$$

$$\frac{1}{2}\frac{\partial A_{c0}}{\partial \tilde{r}} = \frac{\gamma_c}{2} A_{00}` - \Big(2 + \tfrac{\Gamma_{\text{eff}}}{2}\Big) A_{c0} + A_{cc} + \alpha A_{cb}, \tag{133}$$

$$\frac{1}{2}\frac{\partial A_{0b}}{\partial \tilde{\ell}} = \frac{\gamma_b}{2} A_{00} - \Big(1 + \alpha + \tfrac{\Gamma_{\text{eff}}}{2}\Big) A_{0b} + A_{cb} + \alpha A_{bb}, \tag{134}$$

$$\frac{1}{2}\frac{\partial A_{b0}}{\partial \tilde{r}} = \frac{\gamma_b}{2} A_{00} - \Big(1 + \alpha + \tfrac{\Gamma_{\text{eff}}}{2}\Big) A_{b0} + A_{bc} + \alpha A_{bb}, \tag{135}$$

where

$$\Gamma_{\text{eff}} = \gamma_b + \gamma_c.$$

The fully blocked states satisfy the algebraic constraints

$$\frac{\gamma_c}{2}\Big(A_{0c} + A_{c0}\Big) - 3A_{cc} = 0, \tag{136}$$

$$\frac{\gamma_b}{2}\Big(A_{0b} + A_{b0}\Big) - (1 + 2\alpha)A_{bb} = 0, \tag{137}$$

$$\frac{\gamma_b}{2}A_{0c} + \frac{\gamma_c}{2}A_{b0} - (2 + \alpha)A_{bc} = 0, \tag{138}$$

$$\frac{\gamma_c}{2}A_{0b} + \frac{\gamma_b}{2}A_{c0} - (2 + \alpha)A_{cb} = 0. \tag{139}$$

To quantify the correlation structure, we introduce spatial moments of the integrated densities:

$$\begin{aligned}
A^P_{ij} &= \iint_0^\infty A_{ij}(\tilde{\ell}, \tilde{r})\, d\tilde{\ell}\, d\tilde{r}, \\
A^\ell_{ij} &= \iint_0^\infty \tilde{\ell}\, A_{ij}(\tilde{\ell}, \tilde{r})\, d\tilde{\ell}\, d\tilde{r}, \\
A^r_{ij} &= \iint_0^\infty \tilde{r}\, A_{ij}(\tilde{\ell}, \tilde{r})\, d\tilde{\ell}\, d\tilde{r}, \\
A^{\ell r}_{ij} &= \iint_0^\infty \tilde{\ell}\tilde{r}\, A_{ij}(\tilde{\ell}, \tilde{r})\, d\tilde{\ell}\, d\tilde{r}, \\
A^{\ell 2}_{ij} &= \iint_0^\infty \tilde{\ell}^2\, A_{ij}(\tilde{\ell}, \tilde{r})\, d\tilde{\ell}\, d\tilde{r}.
\end{aligned}$$

The total normalization is

$$\sum_{ij} A^P_{ij} = 1.$$

By symmetry,

$$\mathbb{E}[\tilde{\ell}] = \sum_{ij} A^\ell_{ij} = \mathbb{E}[\tilde{r}] = \sum_{ij} A^r_{ij},$$

while

$$\mathbb{E}[\tilde{\ell}\tilde{r}] = \sum_{ij} A^{\ell r}_{ij}, \qquad \mathrm{Var}[\tilde{\ell}] = \sum_{ij} A^{\ell 2}_{ij} - \mathbb{E}[\tilde{\ell}]^2.$$

The Pearson correlation coefficient is therefore

$$\rho = \frac{\mathbb{E}[\tilde{\ell}\tilde{r}] - \mathbb{E}[\tilde{\ell}]^2}{\mathrm{Var}[\tilde{\ell}]}.$$

All moments are obtained by multiplying the master equations by $\tilde{\ell}$, $\tilde{r}$, $\tilde{\ell}\tilde{r}$, or $\tilde{\ell}^2$, integrating over both coordinates, and solving the resulting linear algebraic systems, in direct analogy with the moment calculation for the mean loop size in Sec. 2.2. The resulting correlation coefficient is

$$\rho = \frac{f_1 + f_2}{4(\alpha + 1)^2\, (2\alpha + \gamma_b + \gamma_c + 4)\, (2\alpha(\gamma_c + 3) + 3\gamma_b + \gamma_c + 3)}, \tag{140}$$

where

$$f_1 = (2\alpha+1)(\alpha+1)^2(\gamma_c+12)(2\alpha+\gamma_c+4), \tag{141}$$

$$f_2 = 12\alpha^2\gamma_b^2 + 2\Big[\alpha\Big(\alpha(\alpha+8)+3\Big)\gamma_c + 6\alpha\Big(\alpha(4\alpha+9)+4\Big)+6\Big]\gamma_b. \tag{142}$$

#### 4.1 Limiting cases

Before turning to the general case, it is useful to examine several limiting regimes.

If cohesin–cohesin encounters are rare ($\gamma_c \ll 1$), the correlation is controlled primarily by static barriers. In the limit of very dense, long-lived barriers ($\gamma_b \to \infty$, $\alpha \to 0$), the spectrum becomes degenerate and the loop-length PDF approaches the Γ-distribution discussed above. In the same limit $\rho \to 0$, showing that the two arms become effectively independent.

At the opposite extreme, for very rare or very short-lived barriers ($\gamma_b \to 0$ or $\alpha \to \infty$), the loop-length distribution reduces to a simple exponential and $\rho \to 1$: the two arms remain almost perfectly synchronized because they scarcely feel either barriers or other extruders.

For finite $\alpha$, however, the two arms always retain a finite memory of one another, and the correlation coefficient remains bounded from below:

$$\rho_{\min} = \frac{\alpha^2}{(1+\alpha)^2}, \tag{143}$$

corresponding to the limit of very dense static barriers, $\gamma_b \to \infty$. Thus, as long as barriers can eventually detach, the two arms never become fully independent. A particularly important special case is $\alpha = 1$, which corresponds to cohesin-like obstacle lifetimes and gives

$$\rho_{\min} = \frac{1}{4}. \tag{144}$$

This is exactly the lower bound reached in the system of opaque cohesins without static barriers ($\gamma_b \to 0$) at high cohesin density. Hence even strong cohesin crowding cannot fully decorrelate the two arms.

#### 4.2 Origin of the crowding-induced correlation floor $\rho_{\min}$

The limiting value $\rho_{\min}$ (Eq. (143)) can be derived directly in the high-crowding regime. Let $T$ denote the lifetime of the tagged cohesin, measured in units of its mean lifetime $\tau$. Since cohesin dissociation is Poissonian,

$$\langle T\rangle = \mathrm{Var}(T) = 1. \tag{145}$$

At high obstacle density, the time required for an arm to traverse the short distance to the neighboring obstacle is negligible compared with the time spent waiting for a blocking obstacle to dissociate. Each arm therefore performs a renewal process: after crossing an initial gap, it remains blocked until the neighboring obstacle dissociates, crosses the next gap, and becomes blocked again.

For a blocker with mean lifetime $\tau_b$, define $\alpha = \tau/\tau_b$. Conditional on the tagged-cohesin lifetime $T$, the numbers of blocker-release events on the two sides are independent Poisson variables,

$$N_\ell|T \sim \text{Poisson}(\alpha T), \qquad N_r|T \sim \text{Poisson}(\alpha T). \tag{146}$$

Let the successive gaps be independent exponential variables with mean $d$ and variance $d^2$. The final left-arm extension is then

$$\ell = \sum_{i=0}^{N_\ell} X_i, \tag{147}$$

and analogously for $r$. The initial term accounts for the gap crossed before the first collision. Standard formulas for a random sum give

$$\langle \ell|T\rangle = d(1+\alpha T), \tag{148}$$

$$\text{Var}(\ell|T) = d^2(1+2\alpha T). \tag{149}$$

At fixed $T$, the two arms are statistically independent, so

$$\text{Cov}(\ell, r|T) = 0. \tag{150}$$

Their unconditional covariance arises entirely from the shared lifetime $T$:

$$\begin{aligned}\text{Cov}(\ell, r) &= \text{Cov}(\langle \ell|T\rangle, \langle r|T\rangle) \\ &= d^2\alpha^2\,\text{Var}(T) = d^2\alpha^2. \end{aligned} \tag{151}$$

The variance of either arm follows from the law of total variance,

$$\begin{aligned}\text{Var}(\ell) &= \langle \text{Var}(\ell|T)\rangle + \text{Var}(\langle \ell|T\rangle) \\ &= d^2(1+2\alpha) + d^2\alpha^2 = d^2(1+\alpha)^2. \end{aligned} \tag{152}$$

Consequently,

$$\rho_\infty = \frac{\text{Cov}(\ell, r)}{\sqrt{\text{Var}(\ell)\,\text{Var}(r)}} = \left(\frac{\alpha}{1+\alpha}\right)^2. \tag{153}$$

For collisions between identical cohesins, the blocker and the tagged cohesin have the same mean lifetime, so $\alpha = 1$, yielding

$$\rho_{\min} = \frac{1}{4}. \tag{154}$$

Thus, $1/4$ is not the probability of a particular instantaneous blocking state. Rather, it is the fraction of the total arm-length variance generated by the lifetime shared by the two arms. Long-lived static barriers correspond to $\alpha \ll 1$, for which this shared contribution vanishes and $\rho_\infty \to 0$.

#### 4.3 Synchronization–desynchronization transition

When both static barriers and extruders are present, the dependence of $\rho$ on cohesin density becomes qualitatively richer. For long-lived static barriers ($\alpha < 1$), increasing $\gamma_c$ at fixed $\gamma_b$ can either decrease or increase the arm–arm correlation, defining two distinct regimes. The boundary between them is a synchronization–desynchronization transition (SDT), at which the response of $\rho$ to increasing cohesin density changes sign.

To understand the origin of this transition, it is useful to consider a system that initially contains only static barriers and then gradually add cohesins. At $\gamma_c = 0$, the correlation coefficient is determined entirely by the barrier parameters:

$$\rho_{\gamma_c=0} = \frac{(4+\gamma_b)(12+\gamma_b) + 12\alpha\gamma_b}{4(4+\gamma_b+\alpha\gamma_b)(3+\gamma_b+\alpha\gamma_b)}. \tag{155}$$

Equation (155) already reveals two distinct mechanisms. At low barrier density, $\gamma_b < \gamma_b^{\mathrm{cr}}$, the barrier-only correlation remains close to unity: static barriers are too sparse to substantially decorrelate the two arms. In this case, adding cohesins simply introduces extra steric obstacles, so $\rho$ decreases with increasing $\gamma_c$ and approaches the universal steric bound $\rho_{\min} = 1/4$. We therefore refer to this regime as *cohesin-dominated* or *desynchronizing*.

The situation is reversed at high barrier density, $\gamma_b > \gamma_b^{\mathrm{cr}}$ (Fig. 4 in the main text). For long-lived barriers, the barrier-only value $\rho_{\gamma_c=0}$ can become very small: persistent pauses at static barriers nearly decouple the two arms. In this regime, adding cohesins does not act like adding more static barriers. Unlike long-lived external blockers, cohesins cannot reduce the correlation below the steric bound $1/4$. Consequently, once $\rho_{\gamma_c=0} < 1/4$, increasing $\gamma_c$ must *increase* the correlation, driving the system upward toward the cohesin-controlled asymptote. Physically, this reflects a statistical replacement of long-lived barriers by shorter-lived cohesin blockers. We therefore call this regime *barrier-dominated* or *synchronizing*.

The critical barrier density $\gamma_b^{\mathrm{cr}}$ is defined by the condition that the barrier-only correlation equals the high-cohesin asymptote,

$$\rho(\gamma_b, \alpha, \gamma_c = 0) = \frac{1}{4}. \tag{156}$$

Substituting Eq. (155) gives

$$\gamma_b^{\mathrm{cr}} = \frac{3(1+\alpha)^2(1+2\alpha)}{1+2\alpha-3\alpha^2}. \tag{157}$$

The minimal value of $\gamma_b^{\rm cr}$ is reached for infinitely long-lived barriers, $\alpha = 0$, giving

$$\gamma_b^{\rm cr}(0) = 3. \tag{158}$$

As $\alpha$ increases, barriers become more transient, so the transition shifts to larger $\gamma_b$: a higher barrier density is required to offset the shorter lifetime. Thus $\gamma_b < 3$ provides a simple mnemonic for the cohesin-dominated regime, since below this value static barriers are never dense enough to produce the synchronization–desynchronization transition, regardless of persistence.

Thus, for $\alpha < 1$, the SDT line $\gamma_b = \gamma_b^{\rm cr}$ separates two distinct dynamical regimes. Below it, correlations are controlled mainly by cohesin–cohesin encounters; above it, they are controlled mainly by persistent external barriers. For short-lived barriers ($\alpha > 1$), the denominator in $\gamma_b^{\rm cr}$ changes sign and the transition disappears: such barriers cannot suppress the correlation strongly enough to push the system below the cohesin-controlled bound 1/4. In that case, increasing cohesin density can only reduce the arm–arm correlation.

Under interphase conditions, CTCF barriers are relatively short-lived, with $\alpha$ close to or above unity (Table S2), so only the cohesin-dominated regime is expected. MCM barriers are much more persistent ($\alpha \approx 0$), but their density appears too low to exceed the threshold $\gamma_b^{\rm cr}(0)$. Thus neither class of barriers is expected to drive the system into the barrier-dominated regime in interphase. The physiological system therefore most likely resides in the cohesin-dominated sector, where the arm–arm correlation remains bounded below by 1/4. This residual synchronization explains why the loop-length PDF of the real system never reaches the fully decoupled Γ-distribution limit.

#### 4.4 Arm correlation in a minimal alternating-arm extrusion model

To determine whether intrinsic switching between the active cohesin arms can reproduce the correlation $\rho = 1/4$, we consider a minimal symmetric alternating-arm model. At any instant, only one arm extrudes, and the active state switches from the left arm to the right arm, or vice versa, as a Poisson process with rate $\omega$. Cohesin dissociates independently after an exponentially distributed residence time $T$ with mean $\tau$, and either arm is initially active with equal probability.

We introduce a telegraph variable $s(t) = +1$ when the left arm is active and $s(t) = -1$ when the right arm is active. The accumulated extensions of the two arms can then be written as

$$\ell = \frac{v}{2}(T + Z), \qquad r = \frac{v}{2}(T - Z), \qquad Z = \int_0^T s(t)\, dt, \tag{159}$$

where $v$ is the loop-growth speed. For symmetric Poisson switching,

$$\langle s(t)s(t')\rangle = \exp\left[-2\omega|t - t'|\right]. \tag{160}$$

Averaging the integrated telegraph process over the exponential residence-time distribution gives

$$\mathrm{Var}(Z) = \frac{2\tau^2}{1 + 2\omega\tau}, \tag{161}$$

whereas $\mathrm{Var}(T) = \tau^2$. Since $T$ and $Z$ are uncorrelated by left–right symmetry, the Pearson correlation between the accumulated arm extensions is

$$\rho_{\mathrm{sw}} = \frac{\mathrm{Cov}(\ell, r)}{\sqrt{\mathrm{Var}(\ell)\,\mathrm{Var}(r)}} = \frac{\mathrm{Var}(T) - \mathrm{Var}(Z)}{\mathrm{Var}(T) + \mathrm{Var}(Z)} = \frac{2\omega\tau - 1}{2\omega\tau + 3}. \tag{162}$$

This expression has transparent limiting cases. Perfect synchronization, $\rho \to 1$, is recovered only in the limit of infinitely rapid switching, $\omega\tau \to \infty$, when the two arms sample nearly equal fractions of the same cohesin residence time. The correlation vanishes at $\omega\tau = 1/2$, where the positive covariance induced by the shared residence time is exactly balanced by the negative covariance arising from competition between the two mutually exclusive extrusion states.

For slower switching, $\omega\tau < 1/2$, the model predicts a negative arm–arm correlation. This regime is physically meaningful: because only one arm can move at a given time, an extrusion trajectory that spends an unusually long time extending one arm necessarily spends less time extending the other. Fluctuations in the division of the total residence time $T$ between the two arms therefore dominate over the positive correlation generated by their common lifetime.

In the no-switching limit, $\omega\tau \to 0$, one randomly selected arm remains active throughout the entire trajectory while the other does not advance, and Eq.(162) yields $\rho \to -1/3$. Indeed, in this case the possible arm extensions are $(\ell, r) = (vT, 0)$ and $(0, vT)$ with equal probability. Although $\ell r = 0$ for every trajectory and therefore $\langle \ell r \rangle = 0$, the Pearson covariance is negative because both arms have positive mean extensions: $\mathrm{Cov}(\ell, r) = \langle \ell r \rangle - \langle \ell \rangle \langle r \rangle = -v^2\tau^2/4$. For an exponentially distributed residence time, $\mathrm{Var}(\ell) = \mathrm{Var}(r) = 3v^2\tau^2/4$, which gives $\rho = -1/3$. Thus, negative correlation reflects the mutual exclusivity of arm motion. Increasing the switching rate $\omega\tau > 0$ progressively suppresses this left–right imbalance, drives the correlation through zero at $\omega\tau = 1/2$, and ultimately produces effectively synchronized two-arm extrusion.

Setting $\rho_{\mathrm{sw}} = 1/4$ yields

$$\omega\tau = \frac{7}{6}. \tag{163}$$

Thus, a minimal alternating-arm mechanism reproduces $\rho = 1/4$ when cohesin undergoes, on average, $7/6 \simeq 1.17$ switching events during one residence time. This equivalence concerns the arm correlation alone: additional roadblocks or cohesin–cohesin encounters would generally modify the relation between $\rho$ and the intrinsic switching rate.

### 5. Experimental parameters of extrusion

Table S1 summarizes extrusion parameters reported in different experimental studies. We consider two classes of barrier proteins, CTCF and MCM. CTCF is the best-established and most extensively studied barrier to loop extrusion. MCM, by contrast, has been investigated much less systematically; however, Dequeker et al., 2022 showed that it can also act as a barrier to extrusion and may contribute substantially because of its broad abundance in cells. Our estimates likewise suggest that its effect can be significant, since its effective density is comparable to that of CTCF (Table S2).

Most of the experimental data compiled here for cohesin correspond to HeLa cells. For the barriers (CTCF and MCM) the data are compiled from the datasets of human (mostly HeLa) and mouse cell due to the limited number of studies.

When multiple literature estimates were available, their arithmetic mean was used as the representative value. If uncertainties were reported for most of the contributing estimates (at least half), the uncertainty of the mean was obtained by propagation of the reported errors; otherwise, the sample standard deviation across the literature estimates was used. Uncertainties of subsequently derived model parameters were propagated from these input uncertainties.

The nondimensional parameters estimated for the G1 phase in HeLa cells are listed in Table S2. These values were calculated using the definitions introduced in Sec. 1 together with the averaged experimental estimates summarized in Table S1. For obstacle classes with finite permeability, the effective spacing entering the definition of $\gamma_i$ was taken to be

$$d_i = \frac{\tilde{d}_i}{p_i}, \tag{164}$$

where $\tilde{d}_i$ is the mean genomic spacing between barriers of class $i$, and $p_i$ is the corresponding barrier stopping probability. Equivalently, only a fraction $p_i$ of nominal barriers acts as effective obstacles for extrusion, so that the corresponding dimensionless density is reduced accordingly.

The parameters for the ΔWAPL condition (Table S3) were estimated separately from the main HeLa parameter set. Because the number of available ΔWAPL measurements is limited, and because these measurements were obtained in several different cell types, we average the reported values across systems rather than restricting the analysis to a single cell line.

### 6. Numerical modelling of active extrusion

To validate and complement the analytical results, we built a stochastic, time-resolved simulator of cohesin-driven loop extrusion on a one-dimensional chromatin substrate. The simulator explicitly includes cohesin loading and unloading with finite residence time, one-

Table S1: **Experimental estimates of loop-extrusion parameters used in this work.** The table summarizes literature values for cohesin, CTCF, and MCM residence times, genomic separations, permeabilities, extrusion velocity, and loop length. Most data for cohesin correspond to interphase in HeLa cells; the main exception is MCM, whose barrier properties are currently available primarily for mESCs.

| Protein | Object | Parameter | Value | Error | Method | Source |
|---|---|---|---|---|---|---|
| Cohesin | HeLa | Residence time $\tau$, s | 172 | 36 | FRAP | Brunner et al., 2025 |
| | | | 450[a] | - | FRAP | Wutz et al., 2020 |
| | | | 822 | 132 | FRAP | Holzmann et al., 2019 |
| | | | 418[b] | - | FRAP | Wutz et al., 2017 |
| | | | **465** | **37** | | **Average** |
| | human cohesin, *in vitro* | Velocity $v$, kb/s | 1.0 | 0.4 [c] | in vitro single-molecule fluorescence / TIRF microscopy | Davidson et al., 2019 |
| | HeLa | Velocity $v$, kb/s | 0.85 | - | model fit to Hi-C | Tortora & Fudenberg, 2026 |
| | | | **0.93** | **0.2** | | **Average** |
| | HeLa | Processivity, $l_p$, kb | **431** | **99** | | $l_p = \tau v$ |
| | HeLa | Separation, $d$, kb | 150 | 40 | Average across methods (LC-MS/iFRAP) | Holzmann et al., 2019 |
| | human/mouse datasets | | 165 | 30 | from position of Hi-C minimum | Polovnikov & Starkov, 2026 |
| | | | **158** | **25** | | **Average** |
| | HeLa | Relative number of chromatin-bound cohesins, % | 59 % | 12 % | spot-bleach | Brunner et al., 2025 |
| | | | 60% | 30% | Eq.(6) of main text | |
| CTCF | HeLa | Residence time $\tau_{\mathrm{CTCF}}$, s | 139 | 22 | FRAP | Brunner et al., 2025 |
| | mESC+human U2OS | | 150 | - | average between single-molecule tracking and FRAP + kinetic model | Hansen et al., 2017 |
| | purified CTCF + naked DNA, *in vitro* | | 855[d] | - | single-molecule fluorescence | Davidson et al., 2023 |
| | | | **380** | **410** | | **Average** |
| | HeLa | Raw separation $\tilde{d}_{\mathrm{CTCF}}$, kb | 177 | 34 | FCS | Brunner et al., 2025 |
| | mESC | | 75[e] | 10 | recalc from Hansen et al., 2017 and Cattoglio et al., 2019) | Rahmaninejad et al., 2025 |
| | | | **126** | **18** | | **Average** |
| | HeLa | Stopping probability, $p_{\mathrm{CTCF}}$ | 0.31[f] | 0.06 | orientation-averaged single-molecule loop-extrusion assay | Davidson et al., 2023 |
| | | Separation $d_{\mathrm{CTCF}}$, kb | 406 | 97 | | Calculations (164) |
| MCM | mouse zygotes, HCT116 | Residence time $\tau_{\mathrm{MCM}}$ | >6 hours | | | Dequeker et al., 2022 |
| | paternal mouse zygote | Raw separation $\tilde{d}_{\mathrm{MCM}}$, kb | 75 | - | simulation fit | Dequeker et al., 2022 |
| | HeLa | | 52[g] | - | mass-spectrometry + immunoblotting | Dequeker et al., 2022,Kulak et al., 2014; |
| | | | **64** | **16** | | **Average** |
| | paternal mouse zygote | Stopping probability, $p_{\mathrm{MCM}}$ | 0.2 | - | simulation fit | Dequeker et al., 2022 |
| | | Separation $d_{\mathrm{MCM}}$, kb | 320 | 80 | | Calculations (164) |
| Whole system | HeLa | Loop length, $\lambda$, kb | 200 | - | Hi-C + HiCCUPS | Wutz et al., 2020 |
| | | | 85 | - | nuclease digestion | Jackson et al., 1990 |
| | | | 265 | - | Hi-C + HiCCUPS | Wutz et al., 2017 |
| | | | **183** | **91** | | **Average** |
| | HCT116 | Mean cohesin velocity, $\langle v \rangle$, kb/s | 0.375 | | Hi-C recovery | Rao et al., 2017 |

[a] WT cohesin residence time was estimated for the dynamic cohesin pool as an abundance-weighted average of STAG1- and STAG2-cohesin residence times reported by Wutz et al. (2020)

[b] Wutz et al. reported an iFRAP half-time; the mean exponential residence time was calculated as $\tau = t_{1/2}/\ln 2$.

[c] Recalculated from Supplementary data Davidson et al., 2019

[d] Davidson et al. reported two CTCF dwell-time components, with $t_{1/2} = 1.2$ min (69%) and 29.2 min (31%). Half-times were converted to exponential mean lifetimes, $\tau = t_{1/2}/\ln 2$, and population-weighted, giving $\tau_{\mathrm{CTCF}} \simeq 855$ s.

[e] Calculated from the absolute CTCF abundance reported by Cattoglio et al. and the specifically bound fraction measured by Hansen et al., giving approximately one specifically bound CTCF per 75 kb.

[f] Calculated by averaging the stopping probabilities measured for the two CTCF orientations in the single-molecule assay of Davidson et al.

[g] Dequeker et al. combined the MCM abundance measured by mass spectrometry with the chromatin-bound fraction determined by quantitative immunoblotting to estimate approximately one MCM double hexamer per 52 kb in HeLa cells.

Table S2: Estimated dimensionless extrusion parameters for HeLa cells. The table summarizes the nondimensional parameters inferred from the experimental values listed in Table S1. $\gamma_b^{\text{eff}} = \gamma_{\text{CTCF}}/(1+\alpha_{\text{CTCF}}) + \gamma_{\text{MCM}}/(1+\alpha_{\text{MCM}})$, $\gamma_{\text{dyn}}$ was calculated according to Eq. (43) for $k = 2$, $\gamma_{\text{eff}} = \gamma_b^{\text{eff}} + (\gamma + \gamma_{\text{dyn}})/2$ the effective barrier density. Errors were obtained using the uncertainties in Table S1.

| | $\gamma$ | $\gamma_{\text{CTCF}}$ | $\gamma_{\text{MCM}}$ | $\alpha_{\text{CTCF}}$ | $\alpha_{\text{MCM}}$ | $\gamma_b^{\text{eff}}$ | $\gamma_{\text{dyn}}$ | $\gamma_{\text{eff}}$ |
|---|---|---|---|---|---|---|---|---|
| Value | 2.7 | 1.1 | 1.3 | 1.2 | $\approx 0$ | 1.8 | 0.4 | 3.4 |
| Error | 0.6 | 0.7 | 0.4 | 1.3 | - | 0.6 | 0.1 | 0.7 |

Table S3: Estimated extrusion parameters for the ΔWAPL condition. The table summarizes experimental measurements of cohesin residence time and loop size for ΔWAPL and the corresponding wild-type controls, compiled across several cell types.

| | **Object** | **Parameter** | **ΔWAPL** | **WT** | **Method** | **Source** |
|---|---|---|---|---|---|---|
| Cohesin | HeLa | Residence time $\tau$, s | 2502 | 418 | iFRAP, recalc from $t_{1/2}$ | Wutz et al., 2017 |
| | HeLa | | 1080 | 480 | FRAP | Kueng et al., 2006 |
| | MEF | | 32400 | 1500 | FRAP | Tedeschi et al., 2013 |
| | HAP1 | | 4518 | 576 | FRAP, fitted residence time | Haarhuis et al., 2017 |
| | | **Average** | **10125** | **743** | | |
| | HeLa | Separation $d$, kb | - | 150 | average across methods | Holzmann et al., 2019 |
| | MESC | | - | 187 | average across FCM and "in gel" FCM | Cattoglio et al., 2019 |
| | mouse and human datasets | | - | 165 | from Hi-C minimum | Polovnikov & Starkov, 2026 |
| | | **Average** | **-** | **167** | | |
| Whole system | HeLa | Loop length, $\lambda$, kb | 387 | 265 | Hi-C + HiCCUPS | Wutz et al., 2017 |
| | mouse oocyte | | 390 | 335 | Hi-C + P(s) curve | Silva et al., 2020 |
| | mouse zygote | | 120 | 65 | snHi-C, simulation fit | Gassler et al., 2017 |
| | MESC | | 200 [a] | 160 | Micro-C | Hsieh et al., 2022 |
| | HAP1 | | 575 | 370 | Hi-C + HiCCUPS | Haarhuis et al., 2017 |
| | mESC | | 550 | 125 | Hi-C + RCP analysis | Liu et al., 2025 |
| | | **Average** | **370** | **220** | | |

[a] retrieved from the log-derivative of the contact probability (extended Fig 4e) [Hsieh et al., 2022]

sided or symmetric two-sided extrusion, steric blocking between neighboring loops, and transient positionally immobile roadblocks with tunable spacing and lifetime. From the resulting trajectories, we extract loop-length distributions and mean loop lengths and compare them with the analytical theory.

#### 6.1 Parameters and derived quantities

The numerical system can be specified by the loop-growth speed $v$, cohesin residence time $\tau$, processivity $l_p = v\tau$, cohesin spacing $d$, barrier residence time $\tau_b$, and barrier spacing $d_b$. For comparison with the analytical theory, simulations were performed in dimensionless units with

$$v = 1, \qquad \tau = 1, \qquad l_p = v\tau = 1.$$

The independently varied control parameters were therefore

$$\gamma = \frac{l_p}{d}, \qquad \gamma_b = \frac{l_p}{d_b}, \qquad \alpha = \frac{\tau}{\tau_b},$$

together with the extrusion mode $k \in \{1, 2\}$. In obstacle–free conditions, the per–arm speed is

$$v_{\text{arm}} = \begin{cases} v, & k = 1 \text{ (one–sided)}, \\ \frac{v}{2}, & k = 2 \text{ (two–sided)}. \end{cases}$$

Hence an unblocked loop grows at total rate $v$ in both modes. Time is discretized with a fixed step $\Delta t$, giving a per–arm length increment $\Delta l = v_{\text{arm}} \Delta t$.

The chromatin axis is represented as a 1D segment of length $L$ (kb). To control the mean extruder density, we set $\gamma = v\,\tau/d \Rightarrow d = v\,\tau/\gamma$. Two additional quantities controlled the simulation size: the total number of cohesins, $N_c^{\text{tot}} = 6000$, and the target mean number of cohesins simultaneously bound to chromatin, $n_c = 200$. These determine the total simulation time and the length of the simulated chromatin segment,

$$T = \frac{N_c^{\text{tot}} \tau}{n_c}, \qquad L = n_c d = n_c \frac{l_p}{\gamma}.$$

The time step was fixed at

$$\Delta t = 0.005.$$

Table S4: Parameters of the stochastic extrusion simulations. Simulations were performed in dimensionless units with $v = 1$, $\tau = 1$, and $l_p = v\tau = 1$. The independently varied parameters were $\gamma$, $\gamma_b$, $\alpha$, and the extrusion mode $k$.

| Parameter | Meaning | Definition | Value / range |
|---|---|---|---|
| $v$ | loop-growth speed | – | 1 |
| $\tau$ | cohesin residence time | – | 1 |
| $l_p$ | processivity | $l_p = v\tau$ | 1 |
| $d$ | mean cohesin spacing | $d = l_p/\gamma$ | $1/\gamma$ |
| $d_b$ | mean barrier spacing | $d_b = l_p/\gamma_b$ | $1/\gamma_b$ |
| $\tau_b$ | barrier residence time | $\tau_b = \tau/\alpha$ | $1/\alpha$ |
| $\gamma$ | cohesin density parameter | $\gamma = l_p/d$ | varied |
| $\gamma_b$ | barrier density parameter | $\gamma_b = l_p/d_b$ | varied |
| $\alpha$ | relative barrier turnover rate | $\alpha = \tau/\tau_b$ | varied |
| $k$ | extrusion mode | – | 1 or 2 |
| $\Delta t$ | simulation time step | – | 0.005 |
| $l = v_{\text{arm}}\Delta t$ | mode-dependent | | |
| $N$ | total number of simulated cohesins | $N \equiv N_c^{\text{tot}}$ | 6000 |
| $n_c$ | target mean number of bound cohesins | – | 200 |
| $T$ | total simulation time | $T = N\tau/n_c$ | 30 |
| $L$ | simulated chromatin length | $L = n_c d = n_c l_p/\gamma$ | $200/\gamma$ |

**Loading of cohesins** Before the start of simulation for each cohesin $m = 1, \ldots, N$, we sample:

(i) a start time $t_m^{\text{start}} \sim \text{Uniform}(0, T)$;

(ii) a residence time $T_m \sim \text{Exp}(\tau)$, which determines $t_m^{\text{end}} = t_m^{\text{start}} + T_m$, truncated at $T$ if necessary;

(iii) a loading position $x_m \sim \text{Uniform}(0, L)$ along the 1D axis of length $L$;

(iv) a direction variable $\sigma_m$: for $k = 1$, $\sigma_m \sim \text{Bernoulli}(1/2)$, where $\sigma_m = 0$ corresponds to leftward motion and $\sigma_m = 1$ to rightward motion; for $k = 2$, we set $\sigma_m = 1/2$, corresponding to symmetric bidirectional extrusion.

Each cohesin also carries two dynamical variables: the right-arm length $r_m$ and the left-arm length $\ell_m$. Upon loading, both are initialized to zero,

$$r_m = \ell_m = 0.$$

**Steric blocking between loops** At each time step $t$, we construct the list of loaded cohesins, where cohesin $m$ is considered loaded if

$$t_m^{\text{start}} \leq t < t_m^{\text{end}}.$$

The loaded cohesins are then sorted by their loading positions along the axis. Let the sorted list be

$$\{m_i, x_{m_i}, \ell_{m_i}, r_{m_i}, \sigma_{m_i}\}_{i=1}^{n(t)},$$

where $n(t)$ is the number of loaded cohesins at time $t$.

At each step, each arm attempts to advance by

$$\Delta r_m = v\sigma_m \Delta t, \qquad \Delta \ell_m = v(1 - \sigma_m)\Delta t,$$

provided it is not blocked by another cohesin or by a static barrier.

We define the instantaneous loop edges as

$$E_i^R = x_{m_i} + r_{m_i}, \qquad E_i^L = x_{m_i} - \ell_{m_i}.$$

The right arm of cohesin $m_i$ is allowed to elongate only if it does not overlap with the left arm of the next cohesin $m_{i+1}$,

$$E_i^R + \Delta r_{m_i} < E_{i+1}^L.$$

Similarly, the left arm is allowed to elongate only if it does not overlap with the right arm of the previous cohesin $m_{i-1}$,

$$E_i^L - \Delta \ell_{m_i} > E_{i-1}^R.$$

If the corresponding inequality is violated, that arm is marked as *blocked* for the current step; growth may resume once the neighboring cohesin moves away.

For the first and last cohesins in the sorted list, the absent left or right neighbor is ignored, respectively.

When $t \geq t_m^{\text{end}}$, cohesin $m$ dissociates, its loop is removed, and it no longer constrains neighboring loops.

**Transient static barriers** Static barriers are introduced in the same way as cohesins, by assigning to each barrier both a genomic position and a finite lifetime. Specifically, before the start of simulation for each barrier $j$, we sample:

(i) a position $b_j \sim \text{Uniform}(0, s)$ along the 1D axis of length $L$;

(ii) a start time $t_j^{\text{start}} \sim \text{Uniform}(0, T)$;

(iii) a lifetime $T_j^{(b)} \sim \text{Exp}(\tau_b)$, which determines the removal time

$$t_j^{\text{end}} = t_j^{\text{start}} + T_j^{(b)},$$

truncated at $T$ if necessary.

Thus, at each time step $t$, a barrier $j$ is considered active if

$$t_j^{\text{start}} \leq t < t_j^{\text{end}}.$$

In this way, barriers are distributed uniformly along the genomic axis and undergo stochastic binding and unbinding in time, fully analogously to cohesins.

For each cohesin arm, we then test whether the attempted update would cross the nearest active barrier in the corresponding direction of motion. Denoting by $b_i^{\text{next}}$ the nearest active barrier ahead of the right arm, the right-arm update is permitted only if

$$E_i^R + \Delta r_{m_i} < b_i^{\text{next}}.$$

Similarly, if $b_i^{\text{prev}}$ denotes the nearest active barrier ahead of the left arm in the leftward direction, then the left-arm update is permitted only if

$$E_i^L - \Delta \ell_{m_i} > b_i^{\text{prev}}.$$

If the corresponding condition is violated, that arm is marked as *blocked* for the current step. Growth may resume once the blocking barrier becomes inactive.

In the numerical implementation, collision detection was performed within a tolerance of $2\Delta l$ around the current position of the moving loop edge.

**Recorded observables** At specified sampling times, we record for each active cohesin $m$ the total loop length

$$\lambda_m(t) = \ell_m(t) + r_m(t),$$

which is then used to construct empirical probability density functions $N(\tilde{x})$. The values of $\gamma$, $\gamma_b$, and $\alpha$ were varied independently over the ranges specified in the corresponding parameter sweeps.

#### 6.2 Comparison with the analytical theory

To assess the accuracy and practical range of the mean-field model, we performed stochastic simulations of loop extrusion and compared the outputs to the closed–form predictions for (i) loop–length distributions and (ii) the mean loop scale $\tilde{\lambda}$ as functions of the control parameters. Unless stated otherwise, each datum in the figures below is the average over 5 independent runs at fixed $(\gamma, \gamma_b, \alpha, k)$; error bars denote the sample standard deviation (STD) across runs. In all experiments we used a single static–barrier class, so that $\gamma_b = l_p/d_b$ and $\alpha = \tau/\tau_b$.

##### 6.2.1 Loop–length distributions

**One–sided extrusion ($k$=1).** The analytical prediction for unidirectional extrusion is a purely exponential distribution,

$$N(\tilde{x}) = \beta \exp(-\beta \tilde{x}), \qquad \beta = 1 + \gamma_{\text{eff}},$$

see Eq. (24) for $k = 1$.

In the simulations, we binned $\tilde{x}$ for all loops sampled over the last half of the simulation time and plotted the corresponding empirical PDF. The distributions were compared with the analytical prediction in two ways.

First, the analytical exponential distributions with slopes determined from Eq. (27) were directly compared with the simulated distributions (Fig. S6). As seen in the figure, the simulated distributions have an approximately constant slope on a semi-log scale and closely follow the analytical prediction.

Second, the simulated distributions were fitted with exponentials, and the fitted slopes were compared with the analytical value $1+\gamma_{\text{eff}}$. The results are summarized in Table S5. The fitted slopes for simulations on Fig. S6 are presented in caption.

**Two–sided extrusion ($k$=2).** For symmetric bidirectional extrusion, the theory predicts a sum of three exponentials (Eq. (85)), which generically produces a pronounced peak at finite length followed by an exponential tail. For several parameter settings we overlaid the predicted PDF $N(\tilde{x}) = \sum_{m=1}^{3} C_m e^{-\beta_m \tilde{x}}$ on the empirical histogram. The values of coefficients $\beta_m$, $C_m$ for analytical PDF were calculated according to 3.2.2 in paragraph *Example: one obstacle + opaque cohesin.* For all considered sets of parameters the simulated distributions are very similar to the analytical ones (Figure S7).

##### 6.2.2 Mean loop scale as a function of parameters

We next tested the dependence of the mean dimensionless loop length $\tilde{\lambda} = \langle \tilde{x} \rangle$ on (i) extruder density $\gamma$, (ii) barrier density $\gamma_b$, and (iii) barrier persistence $\alpha$ for both extrusion modes. For each sweep, the varied parameter was scanned on a grid while the remaining two were held fixed. At every tested set of parameters we ran 5 simulations. Simulation

Table S5: Simulated vs analytical slopes of the distribution for one-sided extrusion.

| $\gamma$ | $\gamma_b$ | $\alpha$ | Simulated slope, $\beta_{\text{sim}}$ | Theoretical slope, $1+\gamma_{\text{eff}}$ | Difference, % |
|---|---|---|---|---|---|
| 0.1 | 0.01 | 0.1 | 1.09 | 1.08 | 0.26% |
| 1 | 1 | 0.1 | 2.56 | 2.51 | 2.21% |
| 1 | 2 | 0.1 | 3.22 | 3.39 | 5.11% |
| 2 | 2 | 0.1 | 4.16 | 3.94 | 5.33% |
| 3 | 2 | 0.1 | 4.59 | 4.49 | 2.32% |
| 4 | 2 | 0.1 | 5.24 | 5.02 | 4.49% |
| 5 | 1 | 0.1 | 4.76 | 4.68 | 1.83% |
| 5 | 2 | 0.1 | 5.41 | 5.54 | 2.49% |
| 10 | 2 | 0.1 | 8.55 | 8.13 | 5.28% |
| 1 | 0.1 | 0.1 | 1.76 | 1.74 | 1.22% |
| 1 | 1 | 0.1 | 2.59 | 2.51 | 3.22% |
| 1 | 2 | 0.1 | 3.53 | 3.39 | 3.97% |
| 1 | 3 | 0.1 | 4.15 | 4.29 | 3.12% |
| 1 | 5 | 0.1 | 5.92 | 6.09 | 2.78% |
| 1 | 10 | 0.1 | 10.17 | 10.61 | 4.16% |
| 1 | 2 | 0.1 | 3.53 | 3.39 | 3.97% |
| 1 | 2 | 0.2 | 3.11 | 3.24 | 3.98% |
| 1 | 2 | 0.5 | 3.07 | 2.92 | 5.16% |
| 1 | 2 | 1 | 2.74 | 2.60 | 5.53% |
| 1 | 2 | 2 | 2.23 | 2.28 | 1.86% |
| 1 | 2 | 5 | 2.03 | 1.96 | 3.48% |

estimates were compared to the analytical expressions for loop length given by Eq. (28) for unidirectional extrusion and by Eq. (44)) for bidirectional extrusion.

Figure S8 summarizes the results. In all three parametric sweeps ($\gamma$, $\gamma_b$, $\alpha$) the simulated $\tilde{\lambda}$ closely follows the theoretical curves for both $k$=1 and $k$=2. As predicted theoretically, two–sided extrusion yields systematically larger $\tilde{\lambda}$ at identical $(\gamma, \gamma_b, \alpha)$ due to continued growth by the unblocked arm. Increasing either $\gamma$ or $\gamma_b$ shortens loops, while decreasing $\alpha$ (long–lived barriers) also reduces $\tilde{\lambda}$; the measured trends and absolute values agree with theory within the error bars.

### 7. Fountains

#### 7.1 Numerical solution of the two-arm extrusion equations

The stationary two-arm extrusion equations (131)–(139) were solved numerically in the first quadrant of the dimensionless arm-extension plane,

$$\tilde{\ell} \geq 0, \qquad \tilde{r} \geq 0.$$

Here $\tilde{\ell} = \ell/l_p$ and $\tilde{r} = r/l_p$, where $l_p$ is the bare processivity scale. The goal of the numerical calculation was to obtain a point-source contact kernel, $K(\tilde{\ell}, \tilde{r})$, which gives the probability density of observing a contact between two loci reached by the left and

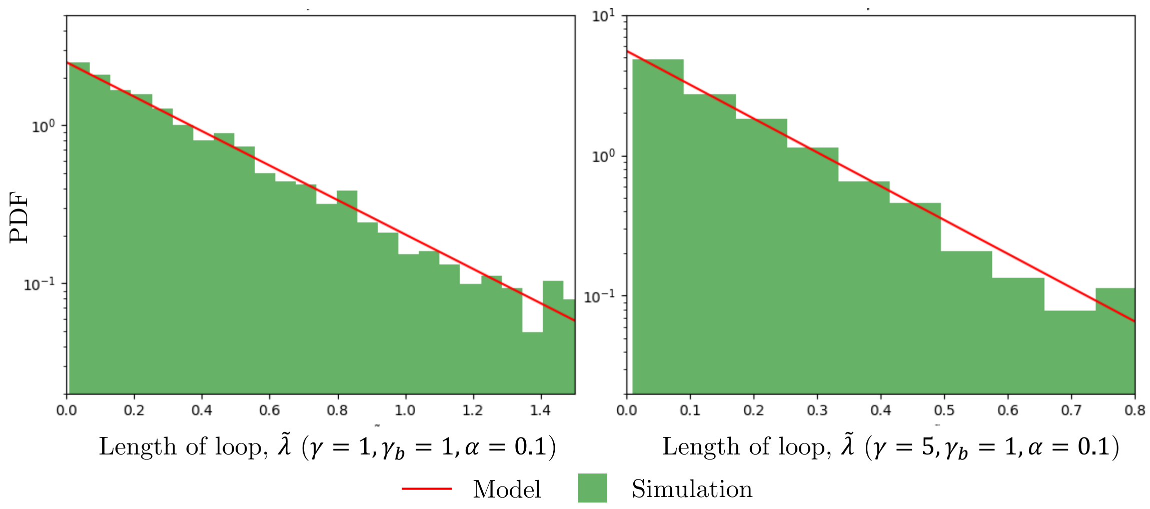

Figure S6: Loop–length distributions, one–sided case ($k$=1). Empirical PDF (green bars) on a semi–log scale vs. the exponential with rate $1 + \gamma_{\text{eff}}$ (line). The empirical slopes ($\beta_{\text{sim}} = 2.56$ on the left, $\beta_{\text{sim}} = 4.76$ on the right) are close to theoretical ones ($1 + \gamma_{\text{eff}} = 2.51$ on the left, $1 + \gamma_{\text{eff}} = 4.68$ on the right). Parameters of the system are shown in the panel.

right arms of a cohesin complex loaded at a fixed position.

The state vector contained the propagating states

$$\mathbf{A} = (A_{00}, A_{0c}, A_{c0}, A_{0b}, A_{b0})\,,$$

where 0 denotes an actively extruding arm, $c$ denotes blockage by another cohesin, and $b$ denotes blockage by a static genomic barrier. The fully blocked states,

$$A_{cc}, \quad A_{bb}, \quad A_{bc}, \quad A_{cb},$$

were not treated as independent dynamic variables. Instead, they were updated locally from the algebraic closure relations after each propagation step. This reduces the numerical problem to a first-order transport-balance system for the five propagating fields.

The equations were discretized on a uniform square grid,

$$\tilde{\ell}_i = ih, \qquad \tilde{r}_j = jh, \qquad i, j = 0, \ldots, N,$$

where $h = L_{\max}/N$. We used

$$L_{\max} = 2.$$

Because the stationary equations are first-order transport equations, they were solved by an upwind marching scheme. The field $A_{00}$, whose transport operator contains derivatives with respect to both arm coordinates, was propagated along diagonal grid directions. The one-sided blocked fields $A_{0c}$ and $A_{0b}$ were propagated along the left-arm coordinate, whereas $A_{c0}$ and $A_{b0}$ were propagated along the right-arm coordinate. At each grid point,

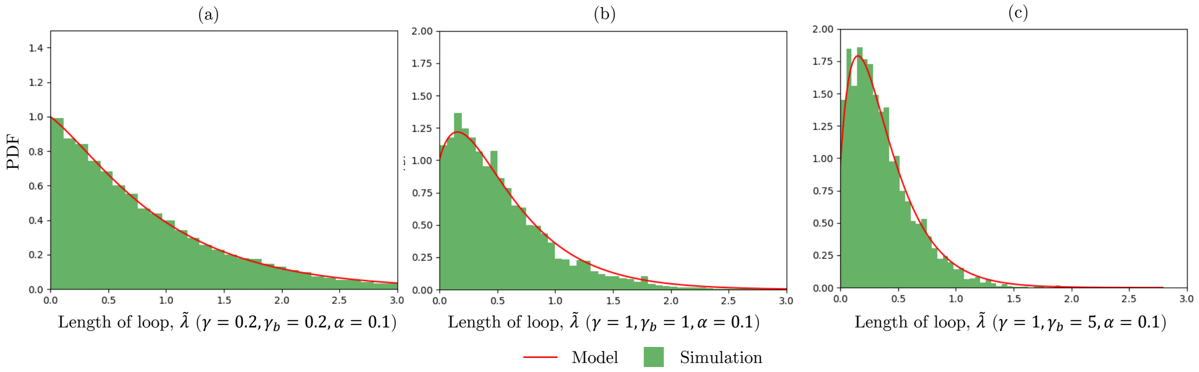

Figure S7: Loop–length distributions, two–sided case ($k$=2). Empirical PDF (green bars) vs. analytical three–mode mixture (red line). Corresponding parameters are stated in the figure. The coefficients of distribution were calculated analytically using the expressions in 3.2.2.

.

the local source and sink terms were evaluated using already computed upwind values, and the algebraic blocked states were then recomputed from the closure equations.

The point-source loading event was implemented as a localized source at the origin. Numerically, the singular source was represented by a narrow regularized loading profile whose width was much smaller than the spatial scale resolved in the final Hi-C comparison. We verified that reducing this regularization width did not change the fitted fountain contours within the resolution of the aggregate maps. In the obstacle-free limit, the boundary condition reduces to the expected exponential propagation of an unblocked extruding complex; this limit was used as a numerical consistency check.

After solving for the propagating states, we constructed the total point-source contact kernel as the sum of all contact-producing states, including both propagating and fully blocked configurations:

$$K(\tilde{\ell}, \tilde{r}) = A_{00} + A_{0c} + A_{c0} + A_{0b} + A_{b0} + A_{cc} + A_{bb} + A_{bc} + A_{cb}.$$

Thus, $K$ represents the time-integrated probability density that a cohesin loaded at the origin generates a contact with arm extensions $(\tilde{\ell}, \tilde{r})$, irrespective of the final blocking state of the two arms.

To compare the point-source solution with aggregate Hi-C fountains, we converted it into a genomic contact map by convolving $K$ with a finite loading-position distribution. For two genomic coordinates $x_1 < x_2$, the predicted contact probability was

$$P(x_1, x_2) = \int_{x_1}^{x_2} K\left(\frac{f - x_1}{l_p}, \frac{x_2 - f}{l_p}\right) \rho(f)\, df.$$

Here $f$ is the genomic position at which cohesin is loaded, and the two arguments of $K$ are the left and right arm extensions required to connect the loading site to the loci $x_1$

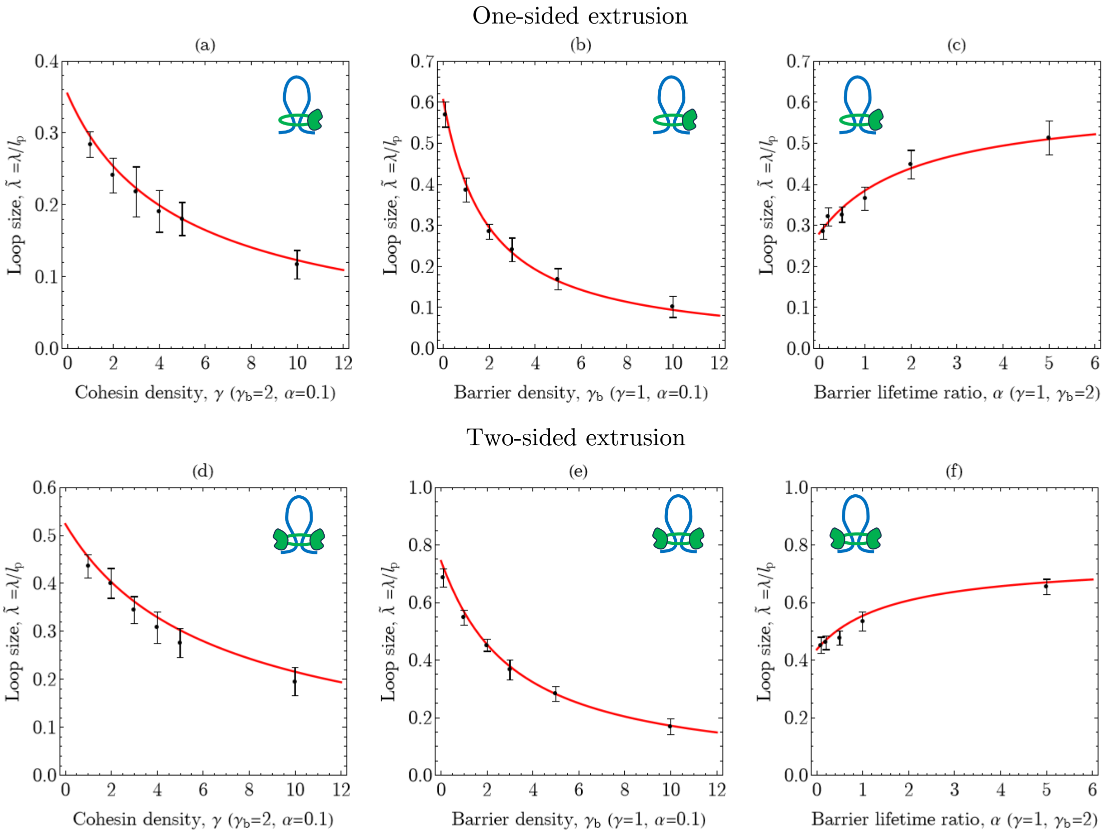

Figure S8: Mean loop scale $\tilde{\lambda}$ vs. parameters for one-sided and two-sided extrusion. Left: $\tilde{\lambda}(\gamma)$ at fixed $(\gamma_b, \alpha)$; Middle: $\tilde{\lambda}(\gamma_b)$ at fixed $(\gamma, \alpha)$; Right: $\tilde{\lambda}(\alpha)$ at fixed $(\gamma, \gamma_b)$. Points: simulation mean over 5 runs; error bars: STD across runs; lines: analytical prediction. The fixed parameters are stated in the figures.

and $x_2$. The loading-position density was taken to be Gaussian,

$$\rho(f) = \frac{1}{\sqrt{2\pi}\sigma_p} \exp\left[-\frac{(f - f_0)^2}{2\sigma_p^2}\right],$$

where $f_0$ is the fountain base and $\sigma_p$ is the physical width of the loading zone.

For numerical evaluation, the convolution integral was computed on a regular grid in genomic coordinates. At each grid point $(x_1, x_2)$, the integration variable $f$ was sampled between the two loci, and the corresponding dimensionless arm lengths were used to interpolate $K$ from the numerical $(\tilde{\ell}, \tilde{r})$ grid. Contributions outside the computed domain were set to zero. The resulting map $P(x_1, x_2)$ was normalized by its maximum value before comparison with the normalized aggregate Hi-C signal.

All fountain-profile fits reported in the main text used the minimal parameter set

$$\theta = (l_p, \sigma_p, \gamma_c),$$

where $l_p$ sets the genomic scale of the fountain, $\sigma_p$ controls the width of the loading zone, and $\gamma_c$ controls the effective local cohesin-crowding strength. The barrier parameter $\gamma_b$ was fixed to the reference value $\gamma_b = 1.8$, and the fits were performed at $\alpha = 0$ (corres-

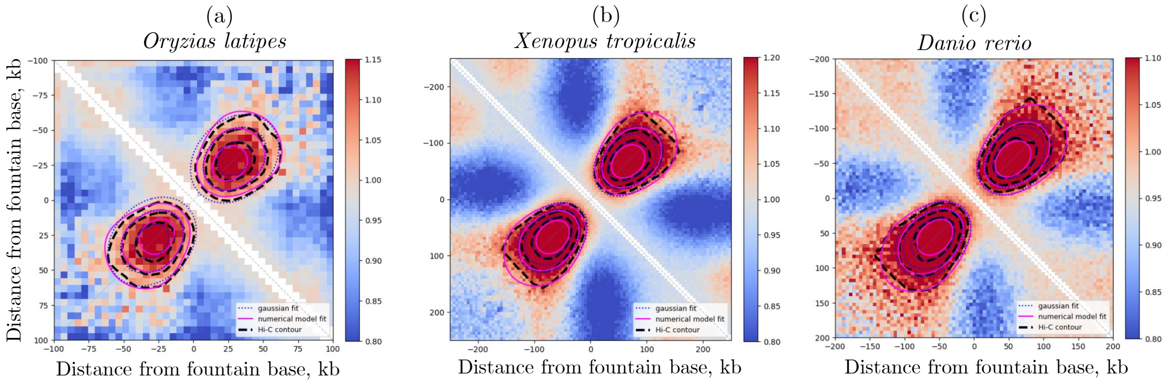

Figure S9: Theoretical and Gaussian fits to extrusion fountains across vertebrates. Aggregated observed/expected Hi-C fountain profiles for **(a)** Medaka (*Oryzias latipes*), **(b)** western clawed frog (*Xenopus tropicalis*), and **(c)** zebrafish (*Danio rerio*). Heatmaps show the normalized Hi-C contact intensity as a function of genomic distance from the inferred fountain base. Dashed black contours trace the experimental fountain signal, dotted blue contours show two-dimensional Gaussian fits to the central fountain region, and solid magenta contours show fits of the numerical kinetic model for the joint distribution of the two cohesin-arm extensions. Both approaches reproduce the characteristic fountain anisotropy, while the numerical model additionally captures the broader profile of the extrusion-associated signal. Fitted parameters, arm–arm correlations, and normalized root-mean-square errors within the central 80-kb and full fitting windows are reported in Table S7.

ponding to $\gamma_b^{\text{eff}} = 1.8$, Table S2). For each candidate parameter set, the numerical system was solved from scratch, convolved with the loading distribution, rescaled to genomic coordinates, and interpolated onto the same grid as the aggregate Hi-C map.

#### 7.2 Construction of aggregate fountain maps

Fountain annotations and Hi-C maps were taken from Galitsyna et al., 2026. We analyzed three vertebrate datasets: *Oryzias latipes*, *Xenopus tropicalis*, and *Danio rerio*. Hi-C signals around fountain bases were extracted at 5-kb resolution using species-specific flanking windows of 200 kb for *Danio rerio*, 250 kb for *Xenopus tropicalis*, and 100 kb for *Oryzias latipes*. All Hi-C maps were analyzed using `cooltools` library.

For each extracted region, the Hi-C contact map was converted to an observed/expected signal by dividing the observed contact frequency by the expected contact probability at the corresponding genomic separation. This normalization removes the average distance-dependent decay of Hi-C contacts and isolates the anisotropic fountain-associated enrichment. Individual base-centered maps were then averaged to obtain one aggregate fountain map per species. The aggregate maps included 1885 fountains for *Oryzias latipes*, 1733 fountains for *Xenopus tropicalis*, and 1460 fountains for *Danio rerio*.

All fitting was performed in the base-centered coordinates. The horizontal and vertical coordinates of the aggregate map are denoted by $x$ and $y$ (in the Gaussian model), with the fountain base placed at the origin. Unless stated otherwise, we fitted the upper-triangular part of the aggregate map, corresponding to contacts between loci located on opposite sides of the fountain base. In this convention, genomic distances to two loci at different sides from the origin can be interpreted as the left and right extrusion-arm extensions. To analyze the shape features of the aggregated fountain, we smoothed the fountain shape and visualized its level contours. For smoothing we used a 6 kb Gaussian kernel. Next, the standard `contour` function of the `matplotlib` library was used to construct the level contours.

Also we computed the NRMSE between the original and smoothed maps in the same fitting window,

$$\mathrm{NRMSE}_{\mathrm{smooth}} = \sqrt{\frac{1}{N}\sum_{i=1}^{N}[H_i - H_i^{\mathrm{smooth}}]^2}.$$

This quantity estimates the small-scale variability of the aggregate map.

#### 7.3 Gaussian fitting of aggregate fountains

As a model-light description of the central fountain geometry, we fitted the aggregate observed/expected maps with an anisotropic two-dimensional Gaussian profile. The fit was performed in base-centered coordinates using the model

$$G(x, y) = C + A \exp\left[-\frac{(x-y)^2}{a^2} - \frac{(x+y-2p)^2}{b^2}\right]. \tag{165}$$

Here $A$ is the amplitude, $C$ is a local background offset, $p$ is the displacement of the fountain maximum from the base along the fountain axis, and $a$ and $b$ are the Gaussian widths perpendicular and parallel to the fountain axis, respectively.

The Gaussian parameters were estimated by nonlinear least-squares fitting using `scipy.optimize.least_squares`. The fit was restricted to upper-triangular part of the aggregate map and excluded pixels close to the main Hi-C diagonal. This exclusion avoids contamination from the strong short-range diagonal signal, which is not part of the fountain feature. To make the fit robust to distal asymmetric tails and background variation, the fitting procedure was performed in two steps. First, a broad window around the expected fountain maximum was used to obtain an initial estimate of the peak position and Gaussian widths. Second, the fit was repeated in a near-peak window centered on the fitted maximum. The second step was used for the final estimate of the central fountain anisotropy.

Because the central enriched pixels are most informative about the fountain axis and width, high-signal pixels were upweighted during fitting. In addition, the distal tail along the major axis was downweighted to prevent asymmetric long-range signal from dominating the estimate of the central Gaussian shape.

The fitted Gaussian widths were converted into an empirical arm–arm correlation coefficient. If the two arm extensions have equal variances and Pearson correlation $\rho$, the variances along the major and minor axes of the fountain are proportional to $1 + \rho$ and

$1-\rho$, respectively. Therefore, for Gaussian widths $b$ and $a$,

$$\rho_{\text{gauss}} = \frac{b^2 - a^2}{b^2 + a^2}.$$

This value was used as a phenomenological estimate of the correlation between the accumulated left and right arm extensions.

##### 7.3.1 Estimation of Gaussian fit robustness

First, we estimated $\rho_{\text{gauss}}$ from the anisotropy of the aggregated fountain within an 80-kb window. To assess the robustness of this estimate, we repeated the analysis using different window sizes (Table S6). The resulting variation in $\rho_{\text{gauss}}$ was quite small. The standard error (SE) of the correlation coefficient was $0.01 - 0.03$, comparable in magnitude to the variation of the estimates across different window sizes. The SE was calculated using the local uncertainty of the fitted Gaussian widths $a$ and $b$ around the optimum, estimated from the least-squares fit and propagated to $\rho_{\text{gauss}}$.

We further tested the sensitivity of the estimated correlation coefficient to the bootstrap procedure. The bootstrap estimates remained consistent with the original values, indicating that the result is robust to resampling. The results are reported in Table S6. Standard deviations between bootstrap samples were small and only weakly dependent on the bootstrap sample size: we used samples of 50 bootstrap replicates, and increasing this number up to 500 did not substantially change the estimates. For each bootstrap replicate, individual fountains were sampled with replacement, aggregated and refitted by Gaussian model.

However, these statistical errors do not account for the uncertainty associated with degeneracy of Gaussian fit parameters. To quantify this, we scanned Gaussian shape parameters $\theta = (a, b, p)$ around the optimum, refitted the linear parameters $A$ and $C$ for each parameter set, and retained near-optimal solutions whose NRMSE remained within the tolerance defined by the empirical difference between the smoothed and raw aggregate fountain ($\text{NRMSE}_{\text{smooth}} \approx 3\%$). Near-optimal solutions were defined by the threshold

$$\text{NRMSE}^2(\theta) \leq \text{NRMSE}^2_{\text{min}} + \text{NRMSE}^2_{\text{smooth}}. \tag{166}$$

where $\text{NRMSE}_{\text{min}}$ is the NRMSE for the optimal parameters of the fit, the reported uncertainty in $\rho_{\text{Gauss}}$ in Table S7 is the range across this set. The same NRMSE-tolerance logic was used to estimate parameter uncertainty for the numerical-model fits.

This yielded an uncertainty of 0.11-0.13 in $\rho_{\text{gauss}}$ (Table S7, main text), substantially larger than the standard error. Thus, the uncertainty of the correlation coefficient is dominated by experimental noise and should be assessed relative to the noise level in the aggregated Hi-C map rather than from the statistical error alone.

Table S6: Robustness of Gaussian-fit correlation estimates. For each species, the Gaussian fit was repeated using different local fitting-window sizes. The table reports the fitted Gaussian correlation coefficient $\rho_{\text{Gauss}}$ for aggregate fountain, the local standard error estimated from uncertainties of parameters $a$ and $b$ of fit, and the bootstrap mean and standard deviation. Bootstrap data were estimated from 50 bootstrap replicates

| Species | Window (kb) | $\rho_{\text{Gauss}}$ | SE | Bootstrap mean | Bootstrap SD |
|---|---|---|---|---|---|
| *Oryzias latipes* | 60 | 0.244 | 0.025 | 0.250 | 0.019 |
| | 80 | 0.251 | 0.027 | 0.254 | 0.017 |
| | 100 | 0.279 | 0.032 | 0.283 | 0.018 |
| | 130 | 0.247 | 0.02 | 0.252 | 0.017 |
| *Xenopus tropicalis* | 60 | 0.334 | 0.009 | 0.331 | 0.010 |
| | 80 | 0.299 | 0.010 | 0.300 | 0.009 |
| | 100 | 0.297 | 0.015 | 0.300 | 0.011 |
| | 130 | 0.325 | 0.009 | 0.326 | 0.009 |
| *Danio rerio* | 60 | 0.319 | 0.015 | 0.326 | 0.018 |
| | 80 | 0.299 | 0.017 | 0.310 | 0.019 |
| | 100 | 0.288 | 0.023 | 0.300 | 0.020 |
| | 130 | 0.314 | 0.016 | 0.324 | 0.018 |

#### 7.4 Fitting aggregate fountains with the numerical extrusion model

The complete aggregate fountain profiles were also fitted by the numerical two-arm extrusion model. For each candidate parameter set, we first solved the stationary two-arm equations on the dimensionless $(\tilde{\ell}, \tilde{r})$ grid and constructed the point-source contact kernel $K(\tilde{\ell}, \tilde{r})$. This kernel was then convolved with a Gaussian loading-position distribution, rescaled to genomic coordinates using the extrusion length scale $l_p$, and interpolated onto the same coordinate grid as the aggregate Hi-C map. As in the Gaussian fitting procedure, the resulting model profile $P(x, y)$ was compared with the experimental map after the rescaling with offset, $A + C \cdot P(x, y)$, where the offset $A$ and amplitude $C$ were optimized by least-squares minimization for each candidate parameter set. The fit quality was quantified by the normalized root-mean-square error,

$$\text{NRMSE} = \sqrt{\frac{1}{N}\sum_{i=1}^{N} [H_i - P_i]^2},$$

where $H_i$ and $P_i$ are the normalized Hi-C and model signals at pixel $i$, respectively, and the sum runs over the selected fitting window. The near-peak error, denoted $\text{NRMSE}^{80\,\text{kb}}$, was computed in the central 80-kb near-peak window used for the fitting. The full-window error, $\text{NRMSE}^{\text{full}}$, was computed over the complete fitting window and was used as a diagnostic of how well the model reproduced the global fountain profile.

Model parameters were optimized using a bounded coordinate-descent procedure. The search was initialized at $l_p = 500\text{kb}$, $\tilde{\sigma}_p = 0.01$, and $\gamma_c = 5$, with parameter bounds $100\text{kb} \leq l_p \leq 10000\text{kb}$, $0.003 \leq \tilde{\sigma}_p \leq 0.06$, and $0 \leq \gamma_c \leq 40$. The initial coordinate steps were 300 kb, 0.001, and 1 for $l_p$, $\tilde{\sigma}_p$, and $\gamma_c$, respectively. At each iteration, the current parameter set and the two neighboring points obtained by adding or subtracting the corresponding step along each parameter axis were evaluated, while the remaining parameters were kept fixed. Candidate values outside the prescribed bounds were clipped to the nearest boundary.

If none of the coordinate moves produced sufficient improvement, all step sizes were multiplied by 0.5. Optimization was terminated when the steps reached $10^{-4}$ for $\tilde{\sigma}_p$, and 0.1 for $\gamma_c$. The same initialization, bounds, and convergence criteria were used for all aggregate maps.

For each species, the optimized numerical profile was used to compute the corresponding theoretical arm–arm correlation coefficient $\rho_{\mathrm{num}}$. Unlike $\rho_{\mathrm{gauss}}$, which is inferred from the local ellipticity of the central Gaussian approximation, $\rho_{\mathrm{num}}$ was calculated from the fitted kinetic parameter $\gamma_c$ using the analytical expression for $\rho(\gamma_b, \gamma_c, \alpha)$, Eq. (140). This provides an independent consistency check between the local Gaussian anisotropy and the full numerical extrusion-model fit.

Table S7: Parameters, arm–arm correlations and goodness of fit for the theoretical and Gaussian descriptions of extrusion fountains. The numerical joint arm-extension distribution was fitted to the aggregated observed/expected fountain profiles for *Oryzias latipes*, *Xenopus tropicalis* and *Danio rerio*. The local cohesin-density parameter $\gamma_c$ was optimized separately for each organism, whereas the static-roadblock contribution was fixed at $\gamma_b = 1.8$, based on the genome-wide estimate for HeLa cells, and the roadblocks were treated as long-lived ($\alpha = 0$). The corresponding arm–arm correlation, $\rho_{\mathrm{num}}$, was not fitted independently but calculated from the optimized model parameters using Eq. 140. The model-light estimate $\rho_{\mathrm{gauss}}$ was obtained independently from the anisotropy of a two-dimensional Gaussian fitted to the central fountain signal. Reported uncertainties denote the fitting uncertainty for the corresponding parameter or derived correlation. Goodness of fit is quantified by the normalized root-mean-square error (NRMSE). $\mathrm{NRMSE}^{80\,\mathrm{kb}}$ is calculated within the central 80-kb region surrounding the fountain peak and therefore emphasizes local anisotropy, whereas $\mathrm{NRMSE}^{\mathrm{full}}$ is calculated over the complete fitting window and assesses the global fountain profile. The similar values of $\rho_{\mathrm{num}}$ and $\rho_{\mathrm{gauss}}$ provide an internal consistency check between the full kinetic model and the model-light Gaussian inference.

| | *Oryzias latipes* | *Xenopus tropicalis* | *Danio rerio* |
|---|---|---|---|
| $\gamma_c$ | $14.4 \pm 3.5$ | $12.3 \pm 5.8$ | $10.1 \pm 4.7$ |
| $\sigma_p$, kb | $18 \pm 5$ | $45 \pm 12$ | $37 \pm 8$ |
| $l_p$, kb | $1700 \pm 500$ | $3800 \pm 900$ | $3200 \pm 700$ |
| $\rho_{\mathrm{num}}$ | $0.27 \pm 0.01$ | $0.28 \pm 0.02$ | $0.28 \pm 0.02$ |
| $\rho_{\mathrm{gauss}}$ | $0.25 \pm 0.11$ | $0.30 \pm 0.11$ | $0.30 \pm 0.13$ |
| $\mathrm{NRMSE}^{80\,\mathrm{kb}}_{\mathrm{num}}$ | 0.09 | 0.08 | 0.07 |
| $\mathrm{NRMSE}^{\mathrm{full}}_{\mathrm{num}}$ | 0.09 | 0.11 | 0.09 |
| $\mathrm{NRMSE}^{80\,\mathrm{kb}}_{\mathrm{gauss}}$ | 0.10 | 0.08 | 0.11 |
| $\mathrm{NRMSE}^{\mathrm{full}}_{\mathrm{gauss}}$ | 0.10 | 0.17 | 0.16 |

Uncertainty of the fitted numerical parameters was assessed by profile-NRMSE analysis. For each parameter in

$$\theta = (l_p, \sigma_p, \gamma_c),$$

we fixed that parameter across a grid of values and re-optimized the remaining two parameters using the same coordinate-descent procedure. This produced a profile curve NRMSE($\theta_j$), where $\theta_j$ is the parameter held fixed during the scan. Flat profile curves indicate poor identifiability, whereas sharply increasing profiles away from the optimum indicate that the parameter is strongly constrained by the fountain shape.

The accepted range of each parameter was defined using a data-driven error floor derived from the aggregate Hi-C map itself. Parameter values were considered practically indistinguishable from the optimum if the increase in mean squared error did not exceed this smoothing-derived floor:

$$\mathrm{NRMSE}^2(\theta) \leq \mathrm{NRMSE}^2_{\mathrm{min}} + \mathrm{NRMSE}^2_{\mathrm{smooth}}.$$

Here NRMSE($\theta$) is the minimum profile NRMSE obtained after re-optimizing the remaining parameters. The value $\mathrm{NRMSE}_{\mathrm{smooth}}$ characterizes small scale variability of aggregated fountain that the smooth numerical model is not expected to reproduce. The uncertainties of parameters estimated this way are reported in Table S7.

To assess the robustness of the inferred $\rho_{\mathrm{num}}$, we evaluated its sensitivity to the choice of $\gamma_b$ (Table S8). We additionally considered the minimal and maximal value $\gamma_b$ in the independently determined range (Table S2) with $\alpha = 0$. The correlation coefficient changes only slightly ($\sim 10\%$) in this range of $\gamma_b$.

Table S8: Sensitivity of the inferred correlation coefficient $\rho_{\mathrm{num}}$ to the choice of the blocking parameter $\gamma_b$ with $\alpha = 0$. $\gamma_b$ varied in the range, corresponding to independently determined uncertainty (Table S2). Values are shown for three species; uncertainties correspond to the estimated uncertainty of $\rho_{\mathrm{num}}$. The reference value used in the main analysis is $\gamma_b = 1.8$.

| $\gamma_b$ | *Oryzias latipes* | *Xenopus tropicalis* | *Danio rerio* |
|---|---|---|---|
| 1.1 | $0.30 \pm 0.02$ | $0.31 \pm 0.03$ | $0.31 \pm 0.03$ |
| 1.8 | $0.27 \pm 0.01$ | $0.28 \pm 0.02$ | $0.28 \pm 0.02$ |
| 2.5 | $0.25 \pm 0.01$ | $0.25 \pm 0.01$ | $0.25 \pm 0.01$ |

### 8. Estimation of loop-length distribution parameters

An important limitation of the estimation of kinetic parameters directly from loop-length distributions is their poor identifiability. Even in the simplest model of bidirectional extrusion with a single type of extrusion barrier (Sec. 3.1.2, *Example: one obstacle and opaque cohesin*), the analytical loop-length distribution contains three exponential contributions whose characteristic scales and relative weights are controlled by four model parameters ($\alpha$, $\gamma$, $\gamma_b$, and $l_p$). In contrast, the experimental distributions contain only a limited number of robust features that can be used for parameter identification, primarily a peak at short-to-intermediate loop lengths and an approximately exponential tail. Moreover, the experimental distributions are relatively noisy and vary between replicates (Fig. S12b). Thus, the experimental profiles do not contain sufficient independent information to constrain all model parameters with equal precision.

Parameter identifiability is further limited because individual model parameters do not affect the distribution independently. As follows from the expression for the effective loop length (Eq. 46), parameters such as $\gamma$, $\gamma_b$, and $\alpha$ enter the characteristic length scale through effective combinations. Changes in these parameters can therefore partially compensate for one another. As a result, substantially different parameter sets may produce very similar loop-length profiles, giving rise to broad, correlated regions of parameter space rather than sharply defined minima for individual parameters (Fig. S12d–f).

For this reason, we do not infer the model parameters directly from the loop-length distributions. Instead, we use independently estimated parameter values as a reference set, construct the corresponding theoretical distributions, and quantitatively compare them with the experimental data (Fig. 2 in the main text, Fig. S12, Table S9). In this comparison, the model parameters were fixed at their independently estimated values (Tables S1 and S2): $\gamma = 2.7$, $\gamma_b^{\mathrm{eff}} = 1.8$, $l_p = 431$ kb, and $\alpha = 1.2$. Table S9 lists the experimental contact-capture datasets used for comparison with the theoretical loop-length distributions and summarizes the corresponding NMAE values.

To assess robustness to parameter uncertainty, we separately examined how strongly the predicted distributions changed when the model parameters were varied within their experimentally constrained ranges (Tables S1 and S2). Figure S13 shows the resulting sensitivity of the theoretical distributions. The characteristic peak at nonzero loop length is preserved throughout the explored parameter ranges. Variation in processivity primarily rescales the genomic length axis and shifts the peak position, whereas changes in $\gamma$, $\gamma_b$, and $\alpha$ mainly affect the shape and tail of the distribution.

Table S9: Experimental datasets used for comparison with the theoretical loop-length distributions. NMAE denotes the normalized mean absolute error between the experimental loop-length distribution and the theoretical prediction with independently determined parameters $\gamma = 2.7$, $\gamma_b^{\mathrm{eff}} = 1.8$, $\alpha = 1.2$, $l_p = 431$ kb (Table S1, S2).

| No. | Cell line | Protein | Method | Source accession | NMAE |
|---|---|---|---|---|---|
| 1 | K562 | CTCF | MNase HiChIP | GSE285087 | 11.3% |
| 2 | K562 | CTCF | ChIA-PET | GSM970216 | 9.5% |
| 3 | GM12878 | CTCF | ChIA-PET | GSM1872886 | 11.6% |
| 4 | GM12878 | RAD21 | ChIA-PET | GSM1436265 | 9.0% |
| 5 | HeLa | CTCF | ChIA-PET | GSM1872888 | 12.8% |

### References

Brunner, A., Morero, N. R., Zhang, W., Hossain, M. J., Lampe, M., Pflaumer, H., et al. (2025). Quantitative imaging of loop extruders rebuilding interphase genome architecture after mitosis. *Journal of Cell Biology*, 224(3), e202405169.

Cattoglio, C., Pustova, I., Walther, N., Ho, J. J., Hantsche-Grininger, M., et al. (2019). Determining cellular CTCF and cohesin abundances to constrain 3D genome models. *Elife*, 8, e40164.

Davidson, I. F., Bauer, B., Goetz, D., Tang, W., Wutz, G., & Peters, J. M. (2019). DNA loop extrusion by human cohesin. *Science*, 366(6471), 1338–1345.

Davidson, I. F., Barth, R., Zaczek, M., van der Torre, J., Tang, W., Nagasaka, K., Janissen, R., Kerssemakers, J., Wutz, G., Dekker, C., et al. (2023). CTCF is a DNA-tension-dependent barrier to cohesin-mediated loop extrusion. *Nature*, 616(7958), 822–827.

Dequeker, B. J., et al. (2022). MCM complexes are barriers that restrict cohesin-mediated loop extrusion. *Nature*, 606(7912), 197–203.

Galitsyna, A., et al. (2026). Extrusion fountains are hallmarks of chromosome organization emerging upon zygotic genome activation. *Nature Communications*, 17(1), 2787.

Gassler, J., Brandão, H. B., Imakaev, M., Flyamer, I. M., Ladstätter, S., Bickmore, W. A., et al. (2017). A mechanism of cohesin-dependent loop extrusion organizes zygotic genome architecture. *The EMBO journal*, 36(24), 3600–3618.

Haarhuis, J. H., van der Weide, R. H., Blomen, V. A., Yáñez-Cuna, J. O., Amendola, M., van Ruiten, M. S., Krijger, P. H., Teunissen, H., Medema, R. H., van Steensel, B., et al. (2017). The cohesin release factor WAPL restricts chromatin loop extension. *Cell*, 169(4), 693–707.

Hansen, A. S., Pustova, I., Cattoglio, C., Tjian, R., & Darzacq, X. (2017). Ctcf and cohesin regulate chromatin loop stability with distinct dynamics. *elife*, 6, e25776.

Holzmann, J., Politi, A. Z., Nagasaka, K., Hantsche-Grininger, M., Walther, N., Koch, B., et al. (2019). Absolute quantification of cohesin, CTCF and their regulators in human cells. *Elife*, 8, e46269.

Hsieh, T. H. S., Cattoglio, C., Slobodyanyuk, E., Hansen, A. S., Darzacq, X., & Tjian, R. (2022). Enhancer-promoter interactions and transcription are maintained upon acute loss of CTCF, cohesin, WAPL, and YY1. *Nature Genetics*, 54(12), 1919–1932.

Jackson, D., Dickinson, P., & Cook, P. (1990). The size of chromatin loops in hela cells. *The EMBO journal*, 9(2), 567–571.

Kueng, S., Hegemann, B., Peters, B. H., Lipp, J. J., Schleiffer, A., Mechtler, K., & Peters, J.-M. (2006). Wapl controls the dynamic association of cohesin with chromatin. *Cell*, 127(5), 955–967.

Kulak, N. A., Pichler, G., Paron, I., Nagaraj, N., & Mann, M. (2014). Minimal, encapsulated proteomic-sample processing applied to copy-number estimation in eukaryotic cells. *Nature Methods*, 11(3), 319–324.

Liu, N. Q., et al. (2025). Extrusion fountains are restricted by WAPL-dependent cohesin release and CTCF barriers. *Nucleic Acids Research*, 53(12), gkaf549.

Polovnikov, K., & Starkov, D. (2026). A universal polymer signature in Hi-C resolves cohesin loop density and supports monomeric extrusion. *Proceedings of the National Academy of Sciences of the United States of America*, 123(16), e2534385123.

Rahmaninejad, H., Xiao, Y., Tortora, M. M., & Fudenberg, G. (2025). Dynamic barriers modulate cohesin positioning and genome folding at fixed occupancy. *Genome Research*, 35(8), 1745–1757.

Rao, S. S., Huang, S. C., St Hilaire, B. G., Engreitz, J. M., Perez, E. M., et al. (2017). Cohesin loss eliminates all loop domains. *Cell*, 171(2), 305–320.

Silva, M. C., Powell, S., Ladstätter, S., Gassler, J., Stocsits, R., Tedeschi, A., Peters, J.-M., & Tachibana, K. (2020). Wapl releases scc1-cohesin and regulates chromosome structure and segregation in mouse oocytes. *Journal of Cell Biology*, 219(4), e201906100.

Tedeschi, A., Wutz, G., Huet, S., Jaritz, M., Wuensche, A., Schirghuber, E., Davidson, I. F., Tang, W., Cisneros, D. A., Bhaskara, V., et al. (2013). WAPL is an essential regulator of chromatin structure and chromosome segregation. *Nature*, 501(7468), 564–568.

Tortora, M. M., & Fudenberg, G. (2026). The physical chemistry of interphase loop extrusion. *Cell Genomics*, 6(3).

Wutz, G., et al. (2017). Topologically associating domains and chromatin loops depend on cohesin and are regulated by CTCF, WAPL, and PDS5 proteins. *The EMBO journal*, 36(24), 3573–3599.

Wutz, G., Ladurner, R., St Hilaire, B. G., Stocsits, R. R., Nagasaka, K., Pignard, B., Sanborn, A., Tang, W., Várnai, C., Ivanov, M. P., et al. (2020). ESCO1 and CTCF enable formation of long chromatin loops by protecting cohesin$^{\mathrm{STAG1}}$ from WAPL. *Elife*, 9, e52091.

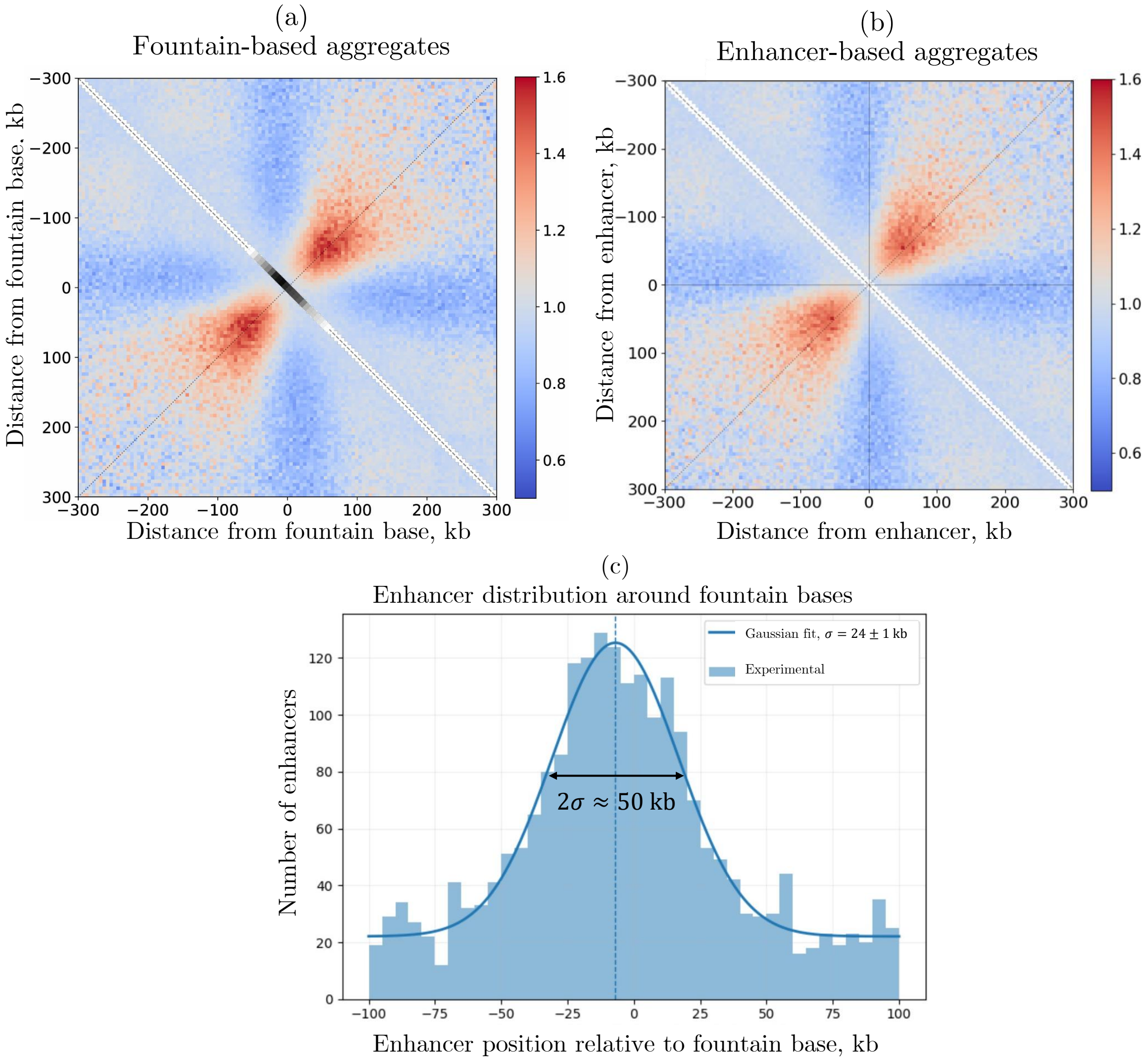

Figure S10: Realignment to the nearest enhancer weakens the aggregated fountain signal. (a) Fountain-centered aggregate obtained by aligning loci to the annotated fountain bases. (b) Aggregate of the same fountains after realignment to the nearest enhancer. The reduced signal and broader pattern after enhancer-based alignment indicate that individual enhancers are genuinely offset from the fountain bases rather than marking their precise centers. (c) Distribution of enhancer positions relative to fountain bases. The distribution is well described by a Gaussian with $\sigma = 24 \pm 1$ kb, corresponding to a characteristic width $2\sigma \approx 50$ kb and indicating that enhancer enrichment extends over a finite region around the fountain base rather than being localized to a single position. Data are from *Danio rerio*, Galitsyna et al, 2026.

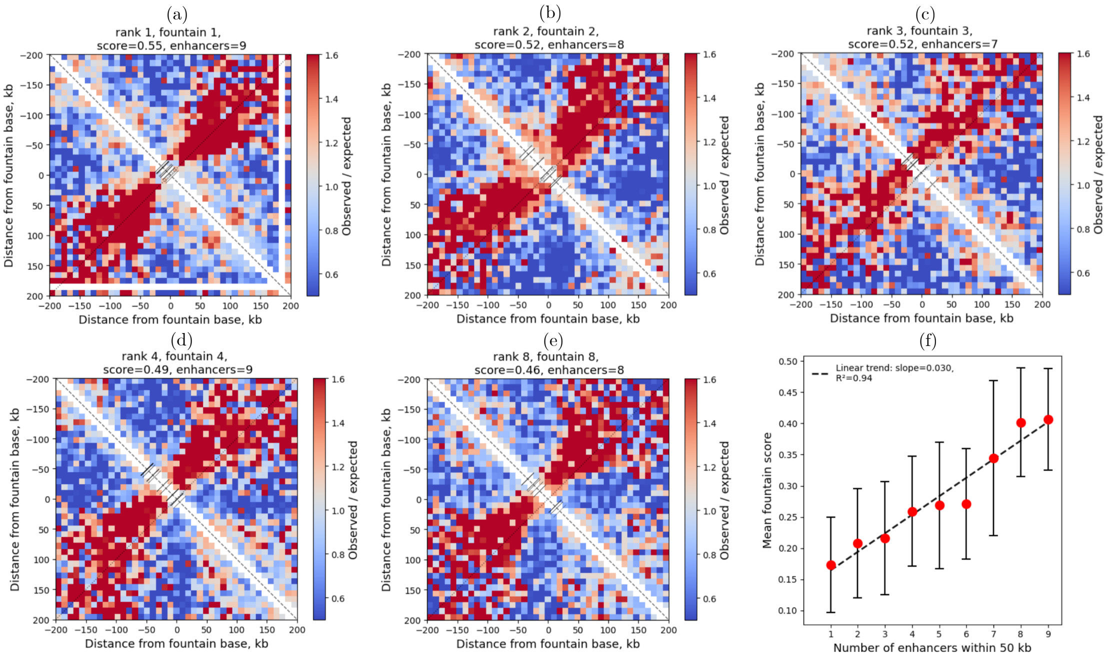

Figure S11: Fountain strength increases with the number of nearby enhancers. (a–e) Representative high-scoring fountains in *Danio rerio*, shown as observed/expected Hi-C maps centered on the inferred fountain base. Gray marks indicate enhancer positions within ±50 kb of the base. Fountain rank, score, and the number of nearby enhancers are indicated above each map. (f) Mean fountain score as a function of the number of enhancers within ±50 kb of the fountain base. Points show group means and error bars indicate variability across fountains. The dashed line shows the linear fit (slope = 0.030, $R^2 = 0.94$), revealing a strong positive association between fountain strength and local enhancer abundance.

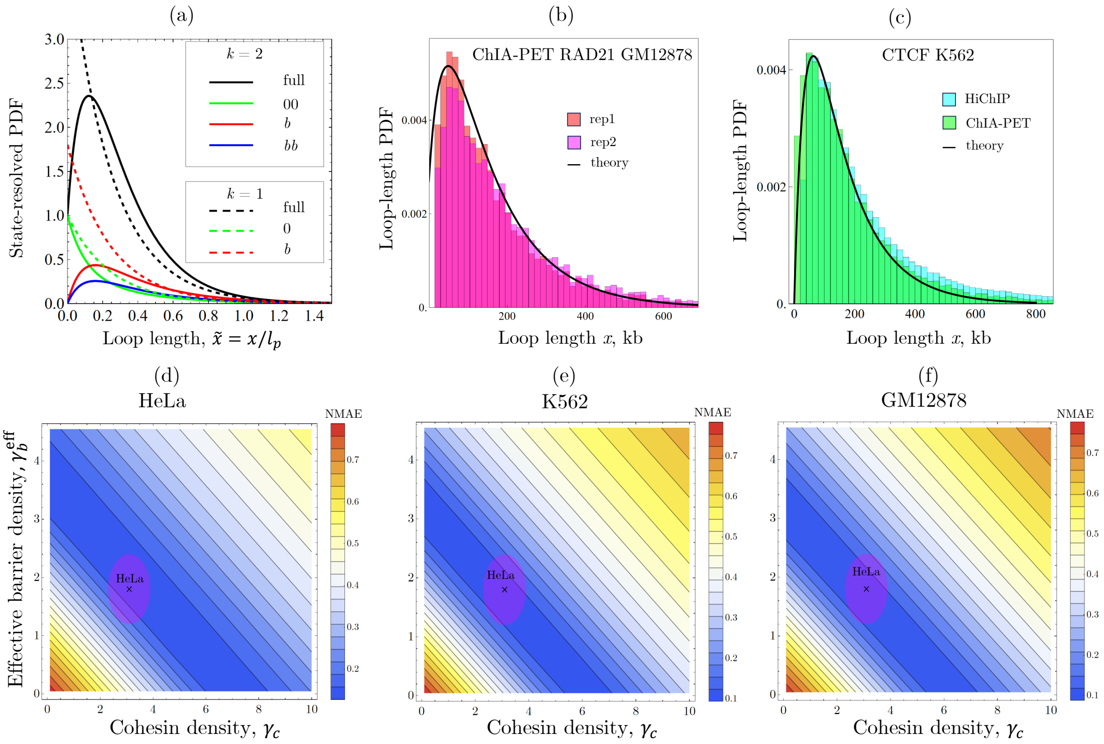

Figure S12: State-resolved loop-length distributions, cross-assay reproducibility, and parameter degeneracy. (a) State-resolved theoretical loop-length distributions for one-sided ($k = 1$, dashed) and two-sided ($k = 2$, solid) extrusion. The full distribution is normalized, while the individual state-resolved components are shown with their relative statistical weights and are not normalized separately. (b) RAD21 ChIA-PET loop-length distributions for two GM12878 replicates compared with the full theoretical distribution $N(x)$ evaluated using the independently estimated parameters. The NMAE between replicates is 16.9%, whereas the NMAE between theory and the replicate-averaged distribution is 9.0%. (c) CTCF loop-length distributions in K562 cells measured independently by MNase HiChIP and PET-weighted ChIA-PET, compared with the same theoretical fully blocked distribution $N_{bb}(x)$. The corresponding NMAEs are 11.3% and 9.5%, respectively. (d–f) NMAE landscapes as functions of cohesin density $\gamma_c$ and effective barrier density $\gamma_b^{\text{eff}}$ at fixed $\alpha = 1.2$ for HeLa, K562, and GM12878, respectively. For HeLa, the NMAE is calculated from CTCF ChIA-PET; for K562, it is averaged over CTCF ChIA-PET and MNase HiChIP; and for GM12878, over CTCF and RAD21 ChIA-PET. The cross marks the independently estimated HeLa parameters, $\gamma_c = \gamma + \gamma_{\text{dyn}}$ , $\gamma = 2.7$, $\gamma_{\text{dyn}} = 0.4$ and $\gamma_b^{\text{eff}} = 1.8$ (Table S2), and the pink ellipse denotes their uncertainty range. The broad low-NMAE valleys illustrate the strong parameter degeneracy of the loop-length distributions, which prevents unique inference of $\gamma$ and $\gamma_b^{\text{eff}}$ from these data alone. Nevertheless, the independently estimated parameter region lies within the low-error regime across all three cell lines.

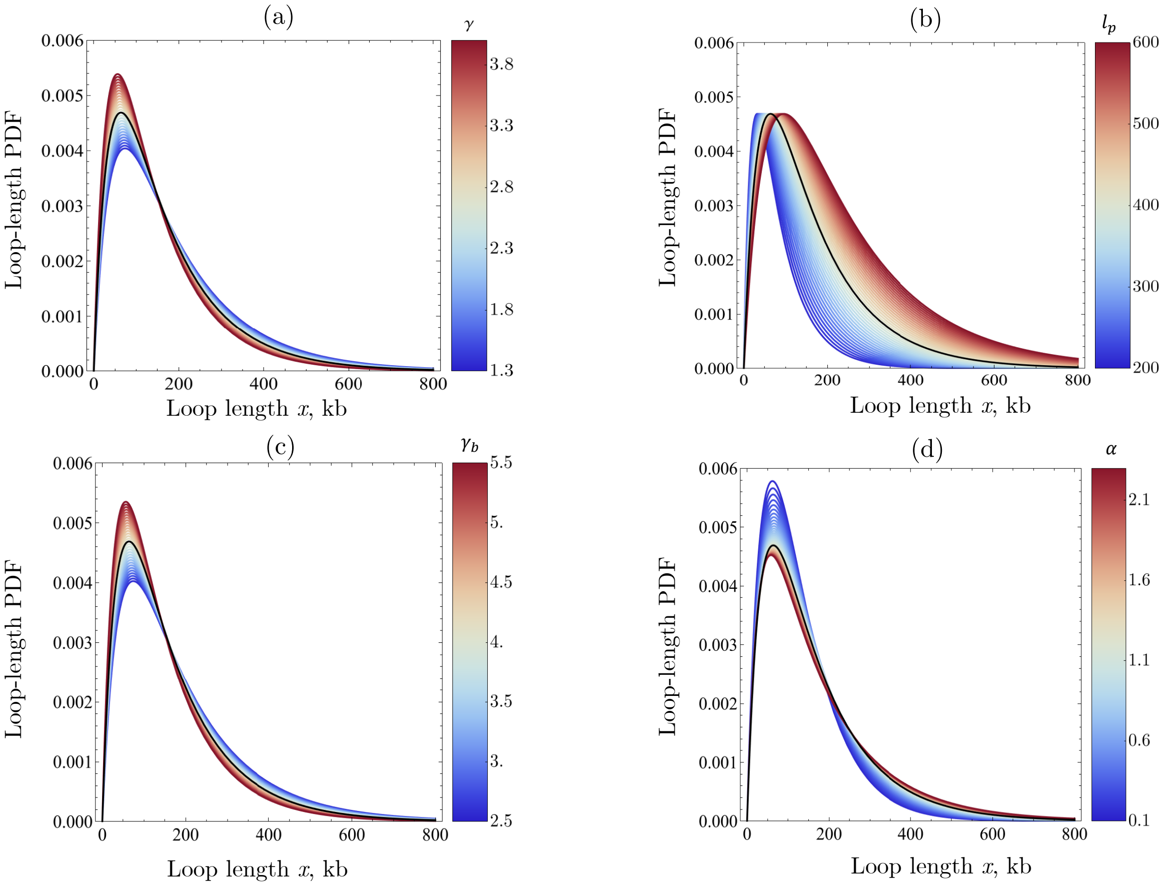

Figure S13: Robustness of the predicted doubly blocked loop-length distribution to experimental parameter uncertainty. Theoretical two-sided distributions $N_{bb}(x)$ obtained by varying one model parameter at a time over its experimentally estimated uncertainty range, while holding the remaining parameters at their central HeLa values (Tables S1,S2). (a) Cohesin density $\gamma$. (b) Bare processivity $l_p = v\tau$ (Table S1). (c) Barrier density $\gamma_b$. For the single effective barrier class used in the loop-length calculation, $\gamma_b$ is obtained from the independently estimated effective density as $\gamma_b = \gamma_b^{\text{eff}}(1+\alpha)$, using $\gamma_b^{\text{eff}} = 1.8 \pm 0.7$ and fixing $\alpha = 1.2$ at its central value. (d) Barrier persistence $\alpha$. The color scale in each panel indicates the varied parameter, and the black curve shows the prediction at the central HeLa parameter values. Across all experimentally constrained variations, the characteristic maximum at nonzero loop length is preserved; uncertainty in $l_p$ primarily shifts the genomic length scale, whereas $\gamma$, $\gamma_b$, and $\alpha$ mainly modulate the shape and tail of the distribution.
